# TGS1 controls snRNA 3’ end processing, prevents neurodegeneration and ameliorates *SMN*-dependent neurological phenotypes in vivo

**DOI:** 10.1101/2020.10.27.356782

**Authors:** Lu Chen, Caitlin M. Roake, Paolo Maccallini, Francesca Bavasso, Roozbeh Dehghannasiri, Pamela Santonicola, Natalia Mendoza-Ferreira, Livia Scatolini, Ludovico Rizzuti, Alessandro Esposito, Ivan Gallotta, Sofia Francia, Stefano Cacchione, Alessandra Galati, Valeria Palumbo, Gian Gaetano Tartaglia, Alessio Colantoni, Gabriele Proietti, Yunming Wu, Matthias Hammerschmidt, Cristiano De Pittà, Gabriele Sales, Julia Salzman, Livio Pellizzoni, Brunhilde Wirth, Elia Di Schiavi, Maurizio Gatti, Steven E. Artandi, Grazia D. Raffa

## Abstract

Trimethylguanosine synthase 1 (TGS1) is a highly conserved enzyme that converts the 5’ mono-methylguanosine cap of snRNAs to a trimethylguanosine cap. Here, we show that loss of TGS1 in *C. elegans, D. melanogaster* and *D. rerio* results in neurological phenotypes similar to those caused by Survival Motor Neuron (SMN) deficiency. Importantly, expression of human TGS1 ameliorates the *SMN*-dependent neurological phenotypes in both flies and worms, revealing that TGS1 can partly counteract the effects of SMN deficiency. TGS1 loss in HeLa cells leads to the accumulation of immature U2 and U4atac snRNAs with long 3’ tails that are often uridylated. snRNAs with defective 3’ terminations also accumulate in *Drosophila Tgs1* mutants. Consistent with defective snRNA maturation, *TGS1* and *SMN* mutant cells also exhibit partially overlapping transcriptome alterations that include aberrantly spliced and readthrough transcripts. Together, these results identify a neuroprotective function for TGS1 and reinforce the view that defective snRNA maturation affects neuronal viability and function.

## INTRODUCTION

The correct processing of small noncoding RNAs is essential to prevent severe neurodegenerative conditions (1). Small nuclear RNAs (snRNAs) are components of the spliceosomal snRNPs. Defects in the snRNP assembly pathway have been observed in Spinal Muscular Atrophy (SMA), a neurological disease caused by deficiency of the Survival Motor Neuron (SMN) protein (2). Another neurodegenerative disorder, Pontocerebellar hypoplasia type 7 (PCH7), is caused by mutations in the TOE1 deadenylase (Target Of EGR1) that prevents accumulation of 3’-end-extended pre-snRNAs (3).

The Sm-class snRNAs include components of both the major (U1, U2, U4, and U5) and the minor (U11, U12, U4atac, and U5) spliceosome. These snRNAs are transcribed by RNA polymerase II and the formation of their 3’ ends is co-transcriptionally controlled (4, 5). Nascent U1 and U2 snRNA transcripts are subject to cleavage by the Integrator complex, resulting in precursors carrying extended 3’ ends that are trimmed after export to the cytoplasm (6–8).

All snRNAs transcribed by Pol II acquire an m^7^-monomethylguanosine cap (m^7^G/MMG cap) at their 5’ end. Immediately after transcription, the monomethylated snRNA precursors bind the Cap Binding Complex (CBC) and are exported to the cytoplasm (9, 10), where they bind the Gemin5 subunit of the SMN complex (11) that mediates their assembly with the seven Sm proteins (Sm core) to form snRNPs (12). The snRNA precursors associated with Gemin5 have extended 3’ tails that can be oligoadenylated or uridylated (11). The U2 snRNA precursors have 3’ extended tails including 12 genomic-encoded nucleotides, which are removed by the TOE1 deadenylase (3).

Following interaction with SMN and the Sm core, the MMG cap of snRNAs is converted to a 2,2,7-trimethylguanosine cap (henceforth abbreviated with TMG cap) by Trimethylguanosine Synthase 1 (TGS1) (13), which physically interacts with SmB and SMN (14). Re-entry of the snRNPs into the nucleus is mediated by both the TMG cap and the Sm core (15, 16). In somatic cells of *Xenopus* and mammals, the Sm signal is both necessary and sufficient for nuclear import, while the TMG cap accelerates the rate of import (17–19). Once in the nucleus, the snRNPs concentrate in the Cajal bodies (CBs) and undergo further maturation steps (20).

3’ end processing of snRNAs is thought to occur after TMG cap formation (21) but whether TGS1 and cap hypermethylation have roles in subsequent steps of snRNP processing is unknown. This is a relevant biomedical issue, as abnormal accumulations of 3’ end extended snRNA precursors have been observed in PCH7 patients (3) and in cells deficient for SMN (11). To elucidate the role of *TGS1* in snRNA maturation and its functional relationships with *SMN*, here we explored the phenotypic consequences of TGS1 loss in human cells and in the *Drosophila*, *C. elegans* and *D. rerio* model organisms. Collectively, our results indicate that TGS1 plays a conserved function in 3’ processing of snRNAs and acts as a neuroprotective factor that can counteract the phenotypic consequences of SMN deficiency.

## MATERIALS AND METHODS

### C.elegans strains

Nematodes were grown and handled following standard procedures, under uncrowded conditions, at 20°C, on NGM (Nematode Growth Medium) agar plates seeded with *Escherichia coli* strain OP50(22). Wild-type animals used in this work were *C. elegans* variety Bristol, strain N2. The transgenic strains used in this work were: NA1330 *gbIs4* [GBF109 p*unc-25::cesmn-1(RNAi)*(200 ng/μL); GB301 p*chs-2::GFP* (10 ng/μL)] III; NA1617 *gbEx593* [GBF109 p*unc-25::cesmn-1(RNAi)* (10 ng/μL); GB301 p*chs-2::GFP* (10 ng/μL)]; NA1541 *gbEx568* [GBF321 p*unc-25::cekal-1(RNAi)* (10 ng/μL); GB301 p*chs-2::GFP* (10 ng/μL)]; NA1252 *gbEx504* [GBF322 p*unc-119*::*dsRED* (20 ng/μL); p*elt- 2*::RFP (30 ng/μL)](23); EG1285 *oxIs12* [p*unc-47*::GFP; *lin-15*(+)] X. EG2185 and N2 were provided by the *Caenorhabditis* Genetics Center (CGC), funded by NIH Office of Research Infrastructure Programs (P40 OD010440); *Is[*p*unc-47::RFP]* was kindly provided by K. Shen (Stanford University, USA).

A complete description of the transgenes obtained in this work is reported in the table below.

| Strain | Transgene name | Transgene description | Notes |
| --- | --- | --- | --- |
| NA1539 | *gbEx579* | [GBF332 *punc-25::cetgs-1(RNAi)*(10 ng/μL); GB301 *pchs-2::GFP* (10 ng/μL)] | *cetgs-1* knock-down in D-type MNs |
| NA1972 | *gbEx647* | [GB335 *punc-119::hTGS1* (20 ng/μL); *podr-1::RFP* (30 ng/μL)] | *hTGS1* overexpression in all neurons |
| NA1981 | *gbEx648* | [GB338 *punc-119::cetgs-1* (20 ng/μL); *podr-1::RFP* (30 ng/μL)]; *gbls4* III | *ceTGS-1* overexpression in all neurons with *smn-1* knock-down in D-type MNs |
| NA1533 | *gbEx576* | [GBF329 *pcetgs-1a::GFP* (50 ng/μl)] | GFP expression under *cetgs-1* isoform *a* promoter |
| NA1535 | *gbEx577* | [GBF330 *pcetgs-1b::GFP* (50 ng/μl)] | GFP expression under *cetgs-1* isoform *b* promoter |

### Transgenic *C.elegans* strains

The constructs GBF329 (p*cetgs-1a*::GFP) and GBF330 (p*cetgs-1b*::GFP) for analyzing the expression of *cetgs-1* isoforms were created by PCR-fusion (24) of two DNA fragments: the presumptive promoters of *cetgs-1* and the GFP sequence. The two promoters of *cetgs-1* isoforms (*a* or *b*) have been chosen upstream of the ATG of the two isoforms, to span 360 bp (*cetgs-1a*) and 960 bp(*cetgs-1b*), respectively. The putative regulatory regions were amplified by PCR using as template wild-type genomic DNA. The GFP, followed by the 3’UTR of the *unc-54* gene to increase the stability of the constructs, was amplified from plasmid pPD95.75, kindly provided by A. Fire (Stanford University, USA). The rescue plasmids (GB335 p*unc-119*:: *hTGS1* and GB338 p*unc- 119*::*cetgs-1*) for pan-neuronal expression were created by cloning *hTGS1* or *cetgs-1* CDS into the plasmid pBY103, kindly provided by O. Hobert (Columbia University, New York, USA), which contains the pan-neuronal promoter p*unc-119*. *hTGS1* and *cetgs-1* CDS were amplified by PCR from cDNA libraries and cloned in BamHI site of pBY103. D-type motoneuron-specific RNAi transgenic lines were obtained as previously described (23, 25). To obtain the specific knock-down of *cetgs-1* we used a short form of *unc-25*/GAD promoter (180 bp), which is specifically expressed from embryonic to adult stages only in the 19 D-type motor neurons in the ventral cord and not in other GABAergic neurons (RME, AVL, DVB and RIS) (23, 26). For gene silencing we amplified, from genomic DNA, the same exon-rich region that was used for the RNAi plasmid library prepared for RNAi by feeding experiments (27). Exon-rich regions were amplified in two separate PCR reactions to obtain the sense and antisense fragments. The *unc-25* promoter was amplified using specific primers. The promoter was subsequently fused to each orientation of the target gene by PCR fusion using internal primers. All primers are available upon request. Germ line transformation was accomplished as described (28) by injecting into the gonad of adult animals a DNA mixture containing a transgenic construct together with a phenotypic marker for selection of transgenic progeny. As co-injection marker we used p*odr-1::*RFP, kindly provided by C. Bargmann, The Rockfeller University, New York, USA (RFP expression in AWC and AWB neurons) at 30 ng/μL and GB301 p*chs-2*::GFP (GFP expression in the glandular pharyngeal g1 and g2 cells, and in m3 and m4 myoepithelial cells)(29) at 10 ng/μL. At least two independent transgenic lines were examined for each experiment. Genetic crosses were made to transfer transgenes to the appropriate genetic background. In all cases, the presence of the desired transgenes was verified by phenotyping. Two independent clones with the same genotype were examined after each cross and the mean of the two clones has been reported in the results. The p*unc- 25* promoter, used for neuron-specific silencing of *cetgs-1* and *cesmn-1* is expressed in 19 neurons among a total of 959 somatic cells and approximately 2000 germ cells in each animal (30), so it is estimated that only 0.6% of total cells have been silenced for *cetgs-1* and *cesmn-1* in each animal (23). Thus, changes in gene expression compared to wild type cannot be appreciated in whole animals. In contrast, *cetgs-1* overexpression is driven in all 302 neurons thanks to the pan-neuronal promoter *punc-119* and through the formation of extrachromosomal arrays, which carry a variable number of copies (between 80 and 300 of the gene of interest (31).

### *C.elegans* behavioral assay

Well-fed, young adult animals were used for backward movement assay (32) to test D-type motor neurons function. The assay was performed blind on NGM plates, 6 cm in diameter, seeded with bacteria. Using an eyelash, the animal was touched first on the tail to induce a forward movement and then on the head to test for backward movement. A defective movement was scored when animals were unable to fully move backward. For each data set, the percentage of animals with normal locomotion among the total number of tested animals was calculated. One-way ANOVA test was used for statistical analysis.

### *C.elegans* microscopy analysis

Animals were immobilized in 0.01% tetramisole hydrochloride (Sigma-Aldrich) on 4% agar pads and visualized using Zeiss Axioskop or Leica DM6B microscopes. All microscopes were equipped with epifluorescence and DIC Nomarski optics and images were collected with Leica digital cameras Hamamatsu C11440. Confocal images have been collected using DMi8 confocal microscope.

### Zebrafish experiments

Zebrafish procedures were approved by the local animal protection committee LANUV NRW; reference 84-02.04.2012.A251. Experiments for caudal primary motor neuron (CaP-MN) analysis were performed in the wild-type TL/EK (*Tupfel long fin* / *Ekkwill*) line.

### Zebrafish injection

Control (non-targeting) and *tgs1* MOs were purchased from (GeneTools, LLC) using the *tgs1* XM_003197865.5 sequence as reference. MO sequences are reported in Table S11. For TGS1 mRNA injection, human *TGS1* cDNA (NM_001317902) was cloned into an N-terminal flag-pCS2+ mRNA expression vector. In vitro transcription of 5’ capped mRNAs was performed using the mMACHINE SP6 Transcription Kit (Ambion) following manufacturer’s protocol and as previously described in (33, 34). Embryos from TL/EK wild-type crossings were used to visualize the CaP-MN phenotype. Embryos were injected with the respective dose of MOs or mRNA in and aqueous solution containing 0.05% PhenolRed and 0.05% Rhodamine-Dextran (Sigma-Aldrich). 6-7hr after injection, embryos were sorted according to homogeneity of the rhodamine fluorescence signal.

### Semi-quantitative RT-PCR of zebrafish samples

RNA isolation from zebrafish larvae (∼34 hours post fertilization - hpf) was performed following the procedure described in (35). RNA was extracted using the RNeasy kit (Qiagen) and concentrations were determined by RiboGreen method (Life Technologies). 600ng RNA were reversely transcribed to cDNA with the Quantitect Reverse Transcription Kit (Qiagen). The RT-PCR experiments were performed as previously described in (36), Multiplex analyses were optimized at 30 cycles to amplify both spliced and unspliced *tgs1* transcripts. In addition to the multiplex analyses, single analyses were performed in the linear phase (22-26 cycles) to ensure reliable quantitative measurements and normalization against endogenous control (*ef1a*).

We observed a clear-dose dependent increase of unspliced transcripts, with increasing Morpholino concentration. Primers were designed using Primer-BLAST and purchased from Integrated DNA technologies (IDT). Primers sequences (Drer-tgs1 and Drer ef1α) are reported in Table S11. Amplification products from semi-quantitative RT-PCR were gel-purified using the QIAquick gel extraction kit (Qiagen) and Sanger sequenced. Densitometric analyses were performed with ImageLab 5.2.1 (BioRAD).

### Immunostaining of zebrafish caudal primary motor neurons

Zebrafish larvae immunostaining was performed as previously described in (33, 34). Briefly, ∼34 hpf zebrafish were manually dechorionated, fixed in 4% PFA-PBS, and permeabilized by Proteinase K digestion of whole larvae. To visualize CaP-MNs, fish larvae were blocked in blocking solution (PBS 0.1% Tween + 5% FCS, 2% BSA and 1% DMSO) and posteriorly incubated overnight at 4°C with mouse α- Znp1 (Synaptotagmin) antibody diluted 1:150 in blocking solution. After five washes of 1 hour with washing solution (PBS 0.1% Tween + 1%FCS + 1%BSA), the secondary mouse α- AlexaFluor488 antibody was diluted 1:250 in blocking solution and incubated at 4°C overnight. Following 5 washes of 20 minutes each, stained fish were stored in 80% glycerol in PBS at 4°C.

For imaging of CaP-MN, fish were laterally embedded in 2% Low Melting Agarose (LMA) microslides under the binocular microscope. Slides were analysed with the fluoresce microscope AxioImagerM2 (Zeiss). The first 10 motor axons posterior to the yolk sac were considered for quantification. Based on overall appearance CaP-MN axons were classified as: normal, truncation (shortened axonal projection) or atrophy (absent axonal projection). Based on terminal branching, axons were classified as Normal, Mild (branching ventral from midline), Medium (2-3 or more branches at ventral or midline) or Severe (>3 branches ventral or dorsal from midline). Brigthfield images of zebrafish larvae were acquired with a Leica S8AP0 binocular attached to an AxioCam ERc5s camera (Zeiss).

### *Drosophila* strains and transgenic constructs

*The UAS-Smn RNAi* construct [P{TRiP.HMC03832}attP40 (UAS-SmnRNAi)] and the *nsyb-GAL4* driver were obtained from the Bloomington Stock Center. The deficiency that removes the *Smn* gene (the *Smn^X7^* allele) is gift from Dr. Artavanis-Tsakonas (37). The *UAS-GFP-dTgs1* strain carries the pPGW*-Tgs1* construct, generated by cloning the *dTgs1* CDS into the pPGW destination vector (Stock number 1077, Drosophila Genomics Resource Center, supported by NIH grant 2P40OD010949), using the Gateway technology (Thermo Fisher Scientific); *the UAS-mst* construct (*UAS-CTRL*) encoding the Misato protein is ppGW-GFP-Mst (38). The *UAS-hTGS1* construct carrying human *TGS1* carries the full-length human *TGS1* gene cloned into the pUAST-attB vector (39). Transgenic flies were obtained by injecting the constructs into either the *w^1118^* or the *y^1^ v^1^; P{CaryP}attP40* (2L, 25C6) strains. All embryo injections were carried out by BestGene (Chino Hills, CA, USA). The *dTgs1^R1^* mutant allele were generated by CRISPR/Cas9 genome editing and described in (40). The *dTgs1^CB0^* mutant allele is the P{PTT-GB}moiCB02140 insertion (41, 42).The Oregon-R strain was used as a wild type control. All flies were reared according to standard procedures at 25°C. Lethal mutations were balanced over either *TM6B, Hu, Tb* or *CyO-TbA, Cy, Tb* (43)*. All* genetic markers and special chromosomes are described in detail in FlyBase (http://www.flybase.org).

### Cell culture, transfections, generation of *TGS1-* and *SMN-* CRISPR HeLa cell lines and Transductions

HeLa S3 cells were cultured in DMEM supplemented with 10% fetal bovine serum and Penicillin-Streptomycin at 37°C, in 5% CO_2_. Lipofectamine 2000 (Life Technologies) was used for all cDNA transfection experiments. TGS1 or SMN CRISPR cells were generated by transfection of HeLa cells with pSpCas9-2A-GFP (PX458) plasmid (44) encoding 3x FLAG Cas9-T2A-GFP and guide RNAs to the *TGS1* locus. (*TGS1-1*: AGAGAAACATTTCCGCCACG; TGS1-2: TGTCAGAGCGTATCTTCAGC)(45) or the SMN locus (SMN-1: CACCCCACTTACTATCATGC). GFP-positive cells were single-cell sorted into 96 well plates by FACS, and clones carrying mutations affecting TGS1 expression were screened by immunoblotting using anti TGS1 or anti- SMN antibody. respectively.

To generate *TGS1* mutant cells stably expressing FLAG-TGS1 (*TGS1* rescued cells), we transfected 293T cells with the pCDH-CMV-PURO-3xFLAG-TGS1 construct (FLAG- TGS1) and packaging constructs; 48 hours later, viral supernatant was collected and concentrated using Retro-X (Clontech). HeLa TGS1-mutant clones were transduced in the presence of 5 μg/mL polybrene and selected in 2 μg/mL puromycin.

For *TGS1* knockdown, UMUC 3 cells (46) were cultured in EMEM EBSS supplemented with 2mM Glutamine, 0.1mM non-essential amino acids, 10% fetal bovine serum, 1.5g/L sodium bicarbonate, 1mM sodium pyruvate and treated for up to 10 days (every 72 hrs) with 50 nM SMARTpool: siGENOME TGS1 siRNA or ON-TARGET plus Non-targeting siRNA using Dharmafect I (Dharmacon). The *PAPD5 KO* and *TCAB1 K1* cell lines are described in (47) and (48). Cumulative population doubling values were determined over a period of 30 days. 1.5 x 10^4^ cells/well were seeded in 6-well plates in duplicates; cells were counted after 72 hrs.

### Western blotting

For western blotting, 10–50 µg of protein extracts were separated by SDS-PAGE, transferred onto PVDF membrane (GE Healthcare, RPN303F), and blotted according to standard procedures. 5% milk in PBST (0.1% Tween) was used for all blocking and antibody incubation steps; PBST (0.1% Tween) was used for all washes. Detection was performed with the SuperSignal™ West Pico Chemiluminescent Substrate (Thermo); images were acquired with Chemidoc (Biorad) and analyzed using the QuantityOne image analysis software (Biorad), or with the Odyssey infrared imaging system (LiCoR). We used the following primary antibodies: FLAG mouse monoclonal (clone M2), Sigma, F1804, 1:1000; PIMT/TGS1: Bethyl A300-814A (lot 1), 1:1500; β-Tubulin: mouse monoclonal (clone D-10) Santa Cruz, SC-5274; SMN: mouse monoclonal Antibody (clone 2B1), 05-1532 Sigma-Aldrich, 1:1000; Coilin: mouse monoclonal [IH10], abcam, (ab87913); CBP20: Rabbit NCBP2, Bethyl laboratory, A302-553A; GAPDH: rabbit, CST, D16H11. Primary antibodies were detected with: HRP-conjugated anti-Mouse or anti-Rabbit 1:5000 (GE Health Care).

### Immunofluorescence staining (IF)

IF experiments were carried out on cells grown on coverslips. Cells were fixed with 4% paraformaldehyde and permeabilized with 0.5% Triton X-100 in PBS. Coverslips were incubated with primary antibodies in 3%BSA for 1 hr at room temperature: monoclonal anti-SMN (05-1532 Sigma-Aldrich clone 2B1, 1:1000); rabbit anti- Coilin (H300 Santa Cruz SC-32860, 200 ug/mL); rabbit anti- TCAB1 ((49), 25 ng/mL). Coverslips were washed 3x with PBS and incubated with secondary Alexa Fluor conjugated antibodies (Jackson Immunoresearch). Coverslips were washed 3x in PBS, counterstained in a 300 nM DAPI solution and mounted in ProLong Gold Anti-fade Mountant. Images were captured on a Leica wide-field fluorescence microscope and processed using Leica LAS AF and Photoshop.

### RNA immunoprecipitation of TMG capped RNA

Trizol-purified RNA was subjected to IP with the R1131 anti-TMG cap specific antibody(50) . 50 µL of protein G beads were washed with PBS and blocked with 20 µg tRNA and 20 µg BSA, then washed with NT2 buffer (50mM Tris/HCl pH 7.5; 150mM NaCl; 1mM MgCl2; 0.05% NP40; 1mM DTT; 100U/mL RNasin; 400µM VRC, Vanadyl ribonucleoside complexes) and coupled O/N to either 20 µL of anti-TMG cap antibody or 1 µg IgG in 250 µL NT2 buffer. Beads were then briefly washed with NT2 buffer and incubated with Trizol-purified RNA in 250 µL NT2 buffer for 2 hrs at 4 °C, while rotating. Beads were then washed 6 times for 2’ while rotating with NT2 buffer and precipitated RNA was Trizol-extracted. Both TMG-immunopurified and input RNAs were then treated with DNase and subjected to reverse transcription and RT-qPCR.

### Quantitative Real Time PCR (RT-qPCR)

Total RNA was extracted with Trizol reagent (Ambion), treated with Ambion™ DNase I (RNase-free) extracted with phenol/chloroform. The integrity of RNA samples was evaluated by gel electrophoresis. 1 µg of intact RNA (with a 28S:18S rRNA ratio = 2:1) was reverse transcribed with the RevertAid H Minus Reverse Transcriptase kit (Thermo Scientific, EP0451). Real-time PCR reactions were performed with the Brilliant II SYBR® Green QPCR Master Mix (Agilent, 600828). The relative quantification in gene expression was carried out using the 2- ΔΔCt method (51). Using this method, we obtained the fold changes in gene expression normalized to the 5.8S gene (the amplification efficiencies were not significantly different for target and reference among all samples). A total of 3 experiments were performed for three biological replicates and the significance was assessed by one-way or two-way ANOVA: For RT-qPCR of immunoprecipitated RNA, samples were processed as above; fold change was calculated by normalizing each RIP sample to the relative input.

In RIP-qPCR analysis the values reported in the histogram bars represent the fold enrichment of RNA in each IP relative to TMG IP from control samples (which was normalized to 1) and were calculated as follows:

fold enrichment of RNA in IP, relative to control IP = 2(-ΔΔCt [RIPtest sample/ RIPCTR], where

ΔΔCt [RIPtest sample/ RIPCTR] =ΔCt [RIP]test sample − ΔCt [RIP]CTR and

ΔCt [RIP]test sample = (Ct [IP] test sample − Ct [Input] test sample −Log2 (Input/IP dilution factor))

Numbers in parentheses represent the percentage of RNA in TMG IP eluates from control samples, relative to input, calculated as: = 2(-ΔCt [RIPtest sample]).

For *C. elegans* experiments, worm pellets were flash-frozen three times before RNA extraction in TRI Reagent® (Merck, Darmstadt, Germany), according to the manufacturer’s protocol. RNA was treated with DNA-free™ DNA Removal (Invitrogen™). Reverse transcription was performed with SuperScript Reverse Transcriptase (Invitrogen™) and Real-time PCR reactions were performed using the Power SYBR Green PCR Master Mix (Applied Biosystems) on the 7900HT Fast Real Time PCR System (Applied Biosystems), following standard procedures. *cetgs-1 e*xpression levels were normalized to the average level of the housekeeping gene *ceact-1*. Each experiment assay was performed in triplicate in three independent experiments Primers are listed in Supplementary table S11.

### Cell fractionation and Northern Blotting

Nuclear extracts (NE) and cytosolic extracts (S100) were obtained using the Dignam protocol as shown in (52). For northern blotting, total RNA is extracted by resuspending pelleted cells with TRIzol reagent. Nuclear and cytoplasmic RNA were extracted from 4µg NE and 12 µg S100 respectively with 5x volume of TRIzol. Precipitated RNAs were dried and resuspended sequentially in 5ul ddH2O, and then with 5ul RNA loading dye, containing 0.1X TBE, 50 mM EDTA, 0.1% Bromophenol blue, 0.1% Xylene Cyanol, 93% formamide. Boiled and rapidly chilled RNAs were fractionated in a 18×24cm 6% Urea-PAGE (19:1) gel, running at 45V for more than 20 hours until the Bromophenol Blue run-off. The gel was then transferred to Hybond-N nitrocellulose membrane under 400mA, for 2 hours in the cold room. Post-transfer gel was stained by ethidium bromide for ribosomal RNAs as loading control. UV-crosslinked membrane was blocked and hybridized in ULTRAhyb, in a 42°C oven overnight, with P^32^ end-labeled oligonucleotide probes (sequence listed in Table S11). The washed membrane was exposed to a phosphor-imager screen, scanned by a Typhoon scanner, and quantitated with the TotalLab software.

### Single molecule fish (Yn in *situ*)

Single molecule FISH protocol was carried out according to the protocol described in (Wu et al., 2021. Yn-situ: a robust single RNA molecule in situ detection method. bioRxiv preprint: https://doi.org/10.1101/2021.10.20.465061). Oligonucleotides used as *Yn situ* probes for U1 and U2 snRNAs are listed in Table S11.

### Nascent RNAend-Seq and steady-state 3’ RACE-Seq

For the isolation of nascent RNA from HeLa cells, cells were pulsed with 1 hr 4SU 250 µM or 4 hr 4SU 50 µM. For chase experiments, 4SU was removed, the cells were washed in PBS, and 2.5 mM uridine containing media was added. Cells were harvested and cell pellets resuspended in Trizol. RNA was extracted using standard Trizol protocol. 100 µg RNA was biotinylated with Biotin-HPDP (Pierce 21341) at 0.5 mg/mL in 40% DMF and 10 mM Tris pH 7.4, 1 mM EDTA, for 1.5 hours at room temperature. In vitro transcribed luciferase RNA transcribed in the presence of 4SU was spiked into the biotinylation mixture for a final concentration of 0.01 ng/µL. RNA was extracted using isopropanol/ethanol precipitation. Biotinylated RNA was immunoprecipitated from total RNA using mMacs Streptavidin Kit (Miltenyi 130-074-101). The depleted fraction was recovered using isopropanol/ethanol precipitation. Biotinylated RNA was released from beads using 100 mM DTT and cleaned using RNeasy MinElute Kit (QIAGEN 74204). Half of eluted RNA or 600 ng of depleted RNA was ligated to 5 µM of 50 adenylated, 30 blocked adaptor (Universal miRNA cloning linker, NEB S1315S) with 250 units of T4 RNA ligase truncated KQ (NEB M0373S), 25% PEG 8000, and 1 uL RnaseOUT (ThermoFisher 10777019) in a 20 µL reaction at 25 degrees for 16 hours. Ligated RNA was cleaned up with RNA clean and concentrator columns (Clontech 740955.50) and DNase treatment, cDNA was synthesized with universal primer and SuperScript III (ThermoFisher 18080093). Amplification was carried out with Phusion (New England Biosystems M0530) and primer sets universal/specific for the RNA of interest. PCR products were directly run on an 8% PAGE gel and visualized with SYBR Gold (ThermoFisher S-11494), or subject to AMPure XP beads (Beckman Coulter A63881) for PCR cleanup and library preparation. Libraries were prepped using Kapa Hyperprep Kit (Kapa KK8504), quantified with Qubit and bioanalyzer, and run on Illumina miSeq at the Stanford Functional Genomics Facility. Reads were paired using fastq-join tool at Galaxy (http://usegalaxy.org). Reads were binned into the various forms of each specific snRNA using custom python scripts (https://cmroake.people.stanford.edu/linkspython-scripts) and the number of reads in each bin was normalized to total snRNA reads. Primer sequences can be found in table S11. Data has been deposited at SRA with BioProject ID PRJNA628085.

### Transcriptome library preparation and RNA-sequencing

Total RNA was extracted with Trizol reagent (Ambion), treated with Ambion™ DNase I (RNase-free) extracted with phenol/chloroform. RNA samples were quantified and quality-tested by Agilent 2100 Bioanalyzer RNA assay (Agilent technologies, Santa Clara, CA). TruSeq Stranded mRNA kit (Illumina, San Diego, CA) has been used for library preparation following the manufacturer’s instructions (library type: fr-firstrand). Final libraries were checked with both Qubit 2.0 Fluorometer (Invitrogen, Carlsbad, CA) and Agilent Bioanalyzer DNA assay. Libraries were then prepared for sequencing and sequenced on paired-end 150 bp mode on NovaSeq 6000 (Illumina, San Diego, CA).

The number of reads (in millions) produced for each sample is listed in the table below.

| **Sample name** | **Reads (M)** |
| --- | --- |
| ID1232_1-1A-CTR1 | 311.29 |
| ID1232_2-1B-CTR2 | 310.44 |
| ID1232_3-2A-TGS1 M1 | 299.96 |
| ID1232_4-3A-TGS1 M2 | 314.74 |
| ID1232_5-4A-SMN C1 | 358.39 |
| ID1232_6-5A-SMN C2 | 272.79 |

Primary bioinformatic analysis includes:

*Base calling and demultiplexing*. Processing raw data for both format conversion and demultiplexing by Bcl2Fastq 2.20 version of the Illumina pipeline. https://support.illumina.com/content/dam/illuminasupport/documents/documentation/soft *ware_documentation/bcl2fastq/bcl2fastq2-v2-20-software-guide15051736-03.pdf* Adapters masking.

Adapter sequences are masked with Cutadapt v1.11 from raw fastq data using the following parameters: --anywhere (on both adapter sequences) --overlap 5 --times 2 --minimum-length 35 --mask-adapter (http://dx.doi.org/10.14806/ej.17.1.200).

Folder “raw_reads” contains files with raw reads (R1: first read sequence; R2: second read sequence) and multiqc_report.html file, which aggregates results from primary bioinformatic analysis into a single report files with parameters that give insight into overall processing and sequencing quality.

### Transcript reconstruction and analysis

Paired-end Illumina sequencing was performed on RNA isolated from TGS1 mutant clones (M1, M2), SMN mutant clones (C1-C2), and parental HeLa cells (CTRL, two replicates) at a depth of 160 million reads/sample. Mutant samples were clustered based on both transcript and gene expression profiles (Supplementary Figure S7). We mapped reads to the human reference genome (Gencode annotation v33) using STAR v2.7.1a (53) and ran Scallop (54) on the resulting BAM files to find all novel transcripts in the data. We combined the Scallop-detected novel transcripts with the reference transcripts and used Kallisto (55) to quantify transcript expression levels. Differentially expressed novel and annotated transcripts were found using Sleuth v0.30.0 (55) (Wald test, FDR adjusted p-value < 0.05, beta value > 2). We compared the 3’ side of each novel transcript to its closest annotated transcript to determine whether the novel transcript has a shorter, extended, or the same 3’ side as compared to the annotated transcript. To investigate the extent of global splicing changes induced by TGS1 and SMN knockouts, we performed differential splicing analysis. We extracted alternative splicing events from the GTF file containing all novel and annotated transcripts and then used SUPPA2 (56) to find significantly changed alternative splicing events. We considered all 7 major categories of alternative splicing events and found the percentage of splicing-inclusion (PSI) for each extracted splicing event. A differentially spliced event between mutant and control samples should have a difference in PSI of at least 0.1 and an adjusted p-value smaller than 0.05.

*Methodological approach used for quantification of intron retention (IR) levels* (Figure 7D): we performed a transcriptome-wide analysis of the expression levels for the major transcript isoforms (i.e., the transcript without the intron) in the intron retention events. IR is quantified by Percentage of Splicing Inclusion (PSI), which is computed as the percentage of reads mapping to the intron-inclusion isoform, relative to total reads mapping to both intron-inclusion and intron-exclusion isoforms. Differential expression of the spliced isoforms (intron exclusion) was computed as the difference in TPM (or transcript per million) values (dTPM) between mutant and control cells (plotted on the X-axis in (Figure 7E, F). The expression of the major transcripts in most of the significant IR events does not change considerably (small difference in TPM) between control and mutant samples (Figure 7E, F). As an independent pipeline for detecting differentially expressed transcripts, we also used a combination of Salmon (57) for transcript quantification and DESeq2 for differential analysis. Statistical analyses on individual transcripts were made with the test implemented in the software.

### Nanopore Sequencing and analyses

#### Data generation

Total RNA was extracted with Trizol reagent (Ambion), treated with Ambion™ DNase I (RNase-free) extracted with phenol/chloroform. All samples have been checked for RIN score above 8 by Agilent 2100 Bioanalyzer RNA assay (Agilent technologies, Santa Clara, CA). The same extracted RNA was used both for Illumina and Nanopore libraries preparation. Nanopore libraries were generated according to PCR-cDNA barcoding (SQK-PCB109) protocol and sequenced on FLO-MIN106 flow cells for 48-72h with -180 mV starting voltage. (See Supplementary Figure S9).

#### Basecalling and Mapping

Samples were sequenced using Oxford Nanopore MinION on FLO-MIN106 flow-cells. Primary data acquisition was performed by MinKNOW. Supplementary Figure S9 reports information about each run. Basecalling and demultiplexing of Fast5 files obtained by sequencing were performed using Guppy v4.2.2. Reads with Qscore > 7 were mapped to hg38 genome using Minimap2 v2.17 (58) with parameters -ax splice.

#### Differential Splicing analysis

To assemble and quantify transcripts from Nanopore reads we used FLAIR v1.4 (59). BAM files obtained with Minimap2 were converted into BED12 files using the bam2Bed12.py utility included in FLAIR. After this, splice junctions of reads were corrected using FLAIR correct module using both the GENCODE v33 GTF file (gencode.v33.chr_patch_hapl_scaff.annotation.gtf) (60) and short Illumina reads of the corresponding samples. BED12 files were then filtered to maintain only reads with at least one splice junction. Filtered BED12 files from all samples were used to assemble the transcriptome using the FLAIR *collapse* module, providing the GENCODE GTF file to annotate the results and requiring at least 10 supporting reads to identify novel transcripts. Transcripts from both the GENCODE and FLAIR GTF files were quantified using the FLAIR *quantify* module. The TPM values were thus calculated and provided to SUPPA2 to identify significantly changed splicing events, following the same procedure used for Illumina short reads.

#### Analysis of transcript readthrough

To study readthrough transcription in WT and mutant samples, we first assigned multi-exonic corrected reads to their corresponding genes starting from the GTF of assembled transcripts obtained by running another run of FLAIR collapse, with minimum number of supporting reads set to 3 to identify all the possible splicing junctions for each gene. Each read was assigned to the gene with which it shared the highest number of splicing junctions. For the identification of the last exon of each gene, we first retained only those transcripts from the reference GTF that have an average TPM of higher than one in at least one condition (WT and TGS1 mutant in WT vs TGS1 comparison, WT and SMN mutant in WT vs SMN comparison); then, for each gene, we selected the last exon of the transcript with the most distal termination site. BEDTools v2.29.1 *intersect* module (61) was employed to select, among the reads assigned to each gene, those mapping to the last exon. Such reads, each one representing a different transcript, were classified as readthrough if their last mapping position was more than 10 nucleotides downstream the termination of the last exon. Differential readthrough analysis was performed by comparing each mutant sample against the pooled WT sample, obtained by combining the reads of both WT samples. For each last exon having at least 10 reads in both WT and mutant samples under analysis, the proportion of readthrough reads in both samples was compared by performing a Fisher’s exact test. This way, we identified Differential Readthrough Events (DREs) as those having a p-value < 0.05. DREs in which the proportion of readthrough reads was higher in the mutant sample and those in which such proportion was higher in the control samples were defined as “Preferential Readthrough in Mutant” and “Preferential Readthrough in CTR” events, respectively. The reads mapping to exons undergoing “Preferential Readthrough in Mutant” events were used to determine the association between readthrough and differential splicing. Readthrough reads from mutant samples were classified as aberrant if their splice junction chains were not found among the non-readthrough reads from the pooled WT sample. The same was done for the non-readthrough reads from mutant samples. The proportions of aberrant reads among readthrough and non-readthrough reads were compared using the chi-square test. To identify the alternative splicing events carried by aberrant reads, we generated several GTF files, each one containing an aberrant read and the non-readthrough reads from the pooled WT sample,and provided such files to SUPPA2 generateEvents module.

To identify fusion transcripts of adjacent genes, we focused on readthrough reads with one or more exons after the termination site and searched for overlaps with downstream genes, defined as those genes for which the first splicing junction is located after the last splicing junction of the gene to which the readthrough reads were assigned. We also evaluated whether at least one splice junction of the overlapping readthrough reads coincided with a splice junction of the downstream gene.

### Phylogenetic alignment

TGS1 homologs were identified by searching in the UniProt knowledgebase (https://www.uniprot.org/). The amino acid alignment was built using Geneious prime using Clustal Omega. To build the phylogenetic tree, pairwise distances were calculated based on the multiple sequence alignment using the Jukes-Cantor distance model. The tree was built using the UPGMA method.

### Statistics

Statistical analyses were performed in GraphPad Prism 6. Type of test, sample size, data representation and p-values are indicated in figure legends. p ≤0.0001 is indicated as ****; p> 0.05 was considered not significant.

## RESULTS

### The *C. elegans* TGS1 ortholog is expressed in motoneurons and required for neuron survival and worm locomotion

To investigate the role of TGS1 in the nervous system and its relationships with SMN biology, we exploited the *C. elegans* model (23). We first analysed the expression of the *C.elegans* ortholog of *TGS1*, *T08G11.4 1* (Supplementary Figure S1) (62), hereafter referred to as *cetgs-1*, using a GFP-reporter approach (24) in which the promoter regions of the two annotated *cetgs-1* isoforms were fused to GFP (p*cetgs-1*::GFP *a* and *b*). Both isoforms are expressed in ventral cord motoneurons (MNs), 19 of which are D-type gamma-aminobutyric acid (GABA) MNs, which are specifically marked by the expression of p*unc-47*::RFP (Figure 1A) (63, 64).

**Figure 1.**
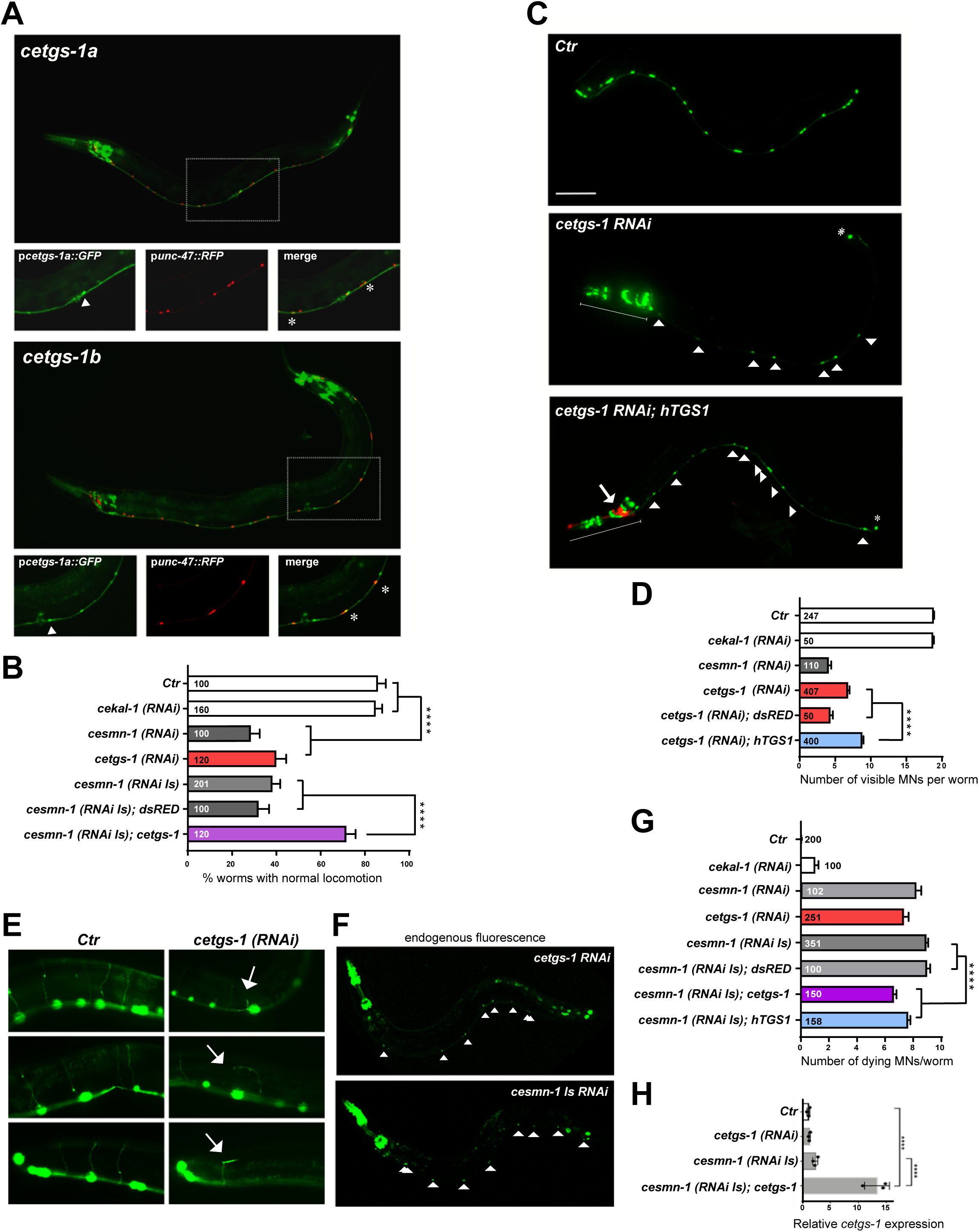
Knockdown of *cetgs-1 or cesmn-1* in D-type MNs results in similar phenotypes that are rescued by *cetgs-1* and *hTGS1* overexpression. **(A)** The promoters of the p*cetgs-1a* and p*cetgs-1b* isoforms drive the expression of GFP in MNs of the ventral cord. D-type (GABA) MNs express RFP under the control of the MN specific promoter (p*unc-47*). In merged images, the D-type MNs expressing both *cetgs-1 (GFP) and* p*unc-47 (RFP)* are marked by asterisks. The cells expressing only *cetgs-1(GFP)* are MNs other than D-type (arrowheads). Anterior is left and ventral is down in all images. Scale bar, 50 µM. **(B)** Knockdown of *cetgs-1 or cesmn-1* in D-type MNs leads to similar locomotion defects. Wild type (Ctr) and ce*kal-1* silenced animals are controls. The locomotion defect elicited by *cesmn-1* silencing is rescued by *cetgs-1* (driven by the pan-neuronal punc-119 promoter) but not by *dsRED* expression. Bars represent the percentage of animals with normal backward locomotion ± SEM, from at least two independent lines/clones. Numbers within bars are the animals tested. **** p<0.0001; (one-way ANOVA). No statistically significant differences were observed between *cesmn-1(RNAi)* and *cetgs-1(RNAi)* (p=0.41) or *cesmn-1(RNAi)* and *cesmn-1(RNAi Is)* (p=0.50). **(C)** Transgenic worms expressing GFP [*oxIs12* (p*unc-47::GFP*)] show 19 D-type MNs of the ventral cord (Ctr)*. cetgs-1* (*RNAi*) worms display fewer GFP expressing MNs in the ventral cord compared to controls. This phenotype is partially rescued by hTGS1 expression. MNs are indicated by arrowheads; the RNAi construct is not expressed in the tail, where one GFP positive cell is always detectable (asterisk); the heads (underlined) express the p*chs-2::GFP* injection marker in both [*cetgs-1* (*RNAi*)] and [*cetgs-1* (*RNAi*); *hTGS1*] worms; the latter were also injected with the p*odr-1::*RFP marker (arrow). Scale bar, 75 µM. **(D)** Quantification of ventral cord D-type MNs. Neuron loss caused by *cetgs-1* knockdown is partially rescued by pan-neuronal expression of human *TGS1* (*hTGS1*) but not of *dsRED*. Each bar represents the mean number of visible MNs from at least two independent lines/clones ± SEM. Numbers within the bars are the animals tested. **** P<0.0001; (One-way ANOVA) **(E)** In *oxIs12* [*punc-47::GFP*] transgenic animals, both MN cell bodies and axons are visible. In control (Ctr) and ce*kal-1(RNAi)* animals, commissures appear as single, straight axons directed to the upper side. *cetgs-1* knockdown worms exhibit commissures with extra branching and guidance defects (arrows). **(F)** Apoptotic autofluorescence signals (arrowheads) in dying MNs of *cetgs-1* (*RNAi)* and *cesmn-1(RNAi Is*) animals. **(G)** Quantification of dying MNs. Bars represent the average number of dying MNs in transgenic animals from at least two independent lines/clones ± SEM. Numbers within and next to bars are the animals tested. **** p<0.0001; (One-way ANOVA). No statistically significant differences were observed by comparing: *cesmn-1(RNAi)* and *cetgs-1(RNAi)* (p=0.27) or *cesmn-1(RNAi)* and *cesmn-1(RNAi Is)* (p=0.08). **(H)** RT-qPCR showing the overexpression of *cetgs-1* in *cesmn-1(RNA Is); cetgs-1* animals, driven by the pan-neuronal punc-119 promoter (>10 fold higher than in wild type). The reduction in cetgs-1 expression could not be detected in cetgs-1(RNAi) animals, as silencing occurs in 19 neurons that represent just 0.6% of total cells. Data are from three biological replicates, are normalized to *ceact-1* and are relative to wild-type animals (Ctr). **** p<0.0001; (One-way ANOVA).

To selectively silence *cetgs-1* in the D-type MNs, we generated a strain carrying the [p*unc-25*::*cetgs-1(RNAi)*] transgene. This strain is viable, fertile and has a normal development. As a control we used a transgenic RNAi line that targets ce*kal-1*, a neurodevelopmental gene not expressed in the D-type MNs (65). We also exploited a *cesmn-1*(*RNAi*) line expressing an extrachromosomal RNAi construct (23). D-type MNs control *C. elegans* backward movement (66). 86% of wild type controls (*wt*) and 85% *cekal-1*-silenced animals, showed normal backward locomotion; this proportion was reduced to 40% and 29% in *cetgs-1*(*RNAi*) and *cesmn-1*(*RNAi*) worms, respectively (Figure 1B), indicating that both *cetgs-1* and *cesmn-1* are required for proper MN function and locomotor behavior.

We next asked whether the *cetgs-1-*dependent locomotion defect is due to an alteration in MN morphology and survival. We introduced in *cetgs-1*(*RNAi*) animals the *oxIs12*[p*unc-47*::GFP] transgene that expresses GFP in D-type MNs, allowing their visualization in living animals. *wt* or *cekal-1*-silenced worms invariably showed 19 D-type MNs, each extending a circumferential commissure to the dorsal side (Figure 1C-E) (66). In *cetgs-1*(*RNAi*) animals the number of visible D-type MNs was in average 7 per worm, while *cesmn-1*(*RNAi*) animals displayed only 4 neurons/worm, as previously reported (23) (Figure 1C, D). The D-type MNs of *cetgs-1*(*RNAi*) animals (n = 606 commissures) showed a defect in axonal morphology, consisting of extra branching and guidance defects (Figure 1E). This defect was present in 14% of the commissures of *cetgs-1*(*RNAi*) worms but was never observed in control (n = 586) or *cekal-1*(*RNAi*) (n = 410) animals (p<0.0001; non-parametric *z*-test.). Remarkably, a similar defect was found in *cesmn-1* knockdown worms (23).

Next, we expressed in D-type MNs either human *TGS1* (*punc-119*::*hTGS1,* referred to as *hTGS1*) or the red fluorescent protein *dsRED* that served as a control (p*unc-119*::*dsRED*). *hTGS1*, but not *dsRED,* partially rescued the MN loss caused by *cetgs-1*(*RNAi*) (from 7 to 9 neurons/animal; Figure 1C, D), indicating that this phenotype is specifically caused by *cetgs-1* silencing and suggesting that the *TGS1* requirement for neuron survival is conserved from worms to humans.

Finally, we asked whether the loss of D-type MNs observed in *cetgs-1*(*RNAi*) worms is caused by cell death. Apoptotic MN death in *cesmn-1* knocked down animals is revealed by the accumulation of an endogenous auto-fluorescent marker (23). *wt* worms showed no dying MNs, while *cekal-1*(*RNAi*) worms showed an average 1 dying MN per animal. In contrast, *cetgs-1*(*RNAi*) and *cesmn-1*(*RNAi*) worms displayed 7 and 8 dying MNs per worm, respectively (Figure 1F, G). Thus, loss of either *cesmn-1* or *cetgs-1* causes D-type MN death.

### *cetgs-1* overexpression mitigates the defects elicited by *cesmn-1* depletion

To determine whether *cetgs-1* interacts genetically with *cesmn-1*, we expressed *cetgs-1* under the control of a pan-neuronal promoter (p*unc-119*::*cetgs-1*, referred to as *cetgs-1*) in worms that also express an integrated *cesmn-1* RNAi construct (*cesmn-1(RNAi Is)* (23). The overexpression of *cetgs-1* was confirmed by RT-qPCR (Figure 1H). 62% of the *cesmn-1*(*RNAi Is*) worms showed abnormal locomotion. Expression of *cetgs-1* in these worms lowered the proportion of animals with abnormal locomotion to 28% (Figure 1B), while *dsRED* expression did not change the frequency of worms with locomotory defects (68%, p = 0.9) (Figure 1B). In addition, while in *cesmn-1*(*RNAi Is*) or *cesmn-1*(*RNAi Is*); dsRED worms, we observed an average of 9 dying MNs/worm, in the presence of the *cetgs-1* or *hTGS1* transgenes, the average number of dying MNs was slightly but significantly lowered to 7 and 8, respectively (Figure 1F, G). Thus, *cetgs-1* overexpression partially suppresses locomotion defects and neuronal death caused by *cesmn-1* silencing.

### Overexpression of dTgs1 ameliorates the Smn loss of function phenotype in Drosophila

We next assayed the effects of TGS1 overexpression in a fly SMA model. It has been reported that *Drosophila* Tgs1 (dTgs1) depletion affects motor behavior (40,67,68). We have previously shown that RNAi against *Smn* in neurons perturbs the circuits that control post-eclosion events, leading to defective wing expansion (69). A similar phenotype has been observed in hypomorphic *dTgs1* mutants (40). We generated flies expressing a *UAS-Smn RNAi* construct in neurons, using the *nsyb-GAL4* driver. 16% of these *Smn RNAi* flies displayed unexpanded wings (Figure 2A, B), and this percentage was increased to 44% when RNAi was performed in flies carrying only one copy of *Smn+* (Δ*Smn/+;* Figure 2B). We then examined flies co-expressing *UAS-Smn-RNAi* and *UAS-dTgs1*, both driven by *nsyb-GAL4*. As control, we used flies carrying a *UAS-CTRL* construct encoding the unrelated Mst protein (38). An increase in the dosage of dTgs1, but not of Mst, significantly lowered the frequency of flies with unexpanded wings (from 16% to 9%) (Figure 2B). A human *TGS1* transgene (*UAS-hTGS1),* which fully rescues the lethality associated with null *dTgs1* mutations (40), decreased the proportion of flies with unexpanded wings from 16% to 10% in *UAS-Smn-RNAi*; *Smn^+^/Smn^+^* flies, and from 44% to 30% in *UAS-Smn-RNAi* flies bearing a single copy of *Smn* (Figure 2B). These results indicate that dTgs1 overexpression partially suppresses the defects caused by *Smn* deficiency in at least a subset of fly neurons, and that human TGS1 can substitute for the neuronal function of its fly ortholog.

**Figure 2.**
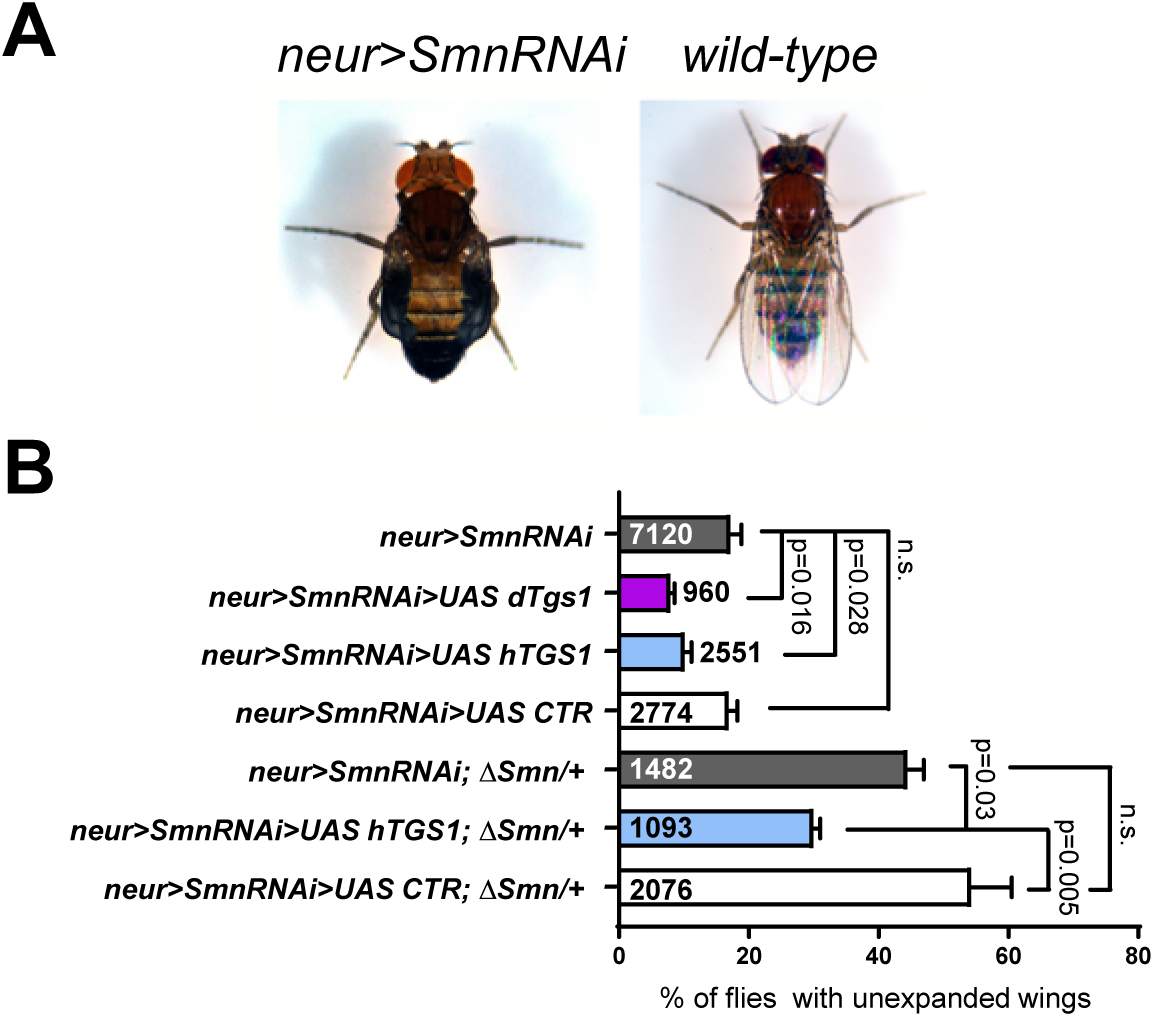
Tgs1 overexpression ameliorates the wing-expansion defects caused by neuronal knockdown of *Smn* in *Drosophila*. **(A)** Examples of flies showing wing expansion failure upon nsyb-GAL4 driven (neur>) *Smn* RNAi. **(B)** Frequencies of the defective wing expansion phenotype in flies co-expressing a *UAS-Smn RNAi* (*Smn RNAi*) construct and the indicated UAS-constructs driven by *nsyb-GAL4* (*neur>*). *UAS dTgs1* encodes GFP-tagged dTgs1; *UAS hTGS1* encodes GFP-tagged human TGS1; *UAS CTR* is a control construct expressing the unrelated Mst protein. *ΔSmn/+* flies are heterozygous for a deficiency of the *Smn* locus. Error bars: ± SEM. Numbers within and next to bars are the animals tested. p values: One-way ANOVA with Tukey’s multiple comparisons test.

### TGS1 downregulation in zebrafish causes motor axon defects

To determine the consequences of TGS1 loss in a vertebrate model, we exploited zebrafish. To downregulate *tgs1*, we injected larvae with two antisense morpholino oligonucleotides (MOs): a translation blocking MO (*tgs1* ATG-MO) and a splicing-blocking MO (*tgs1* Sp-MO) (Supplementary Figure S2A, B). Each MO was injected at an optimized dosage and caused little or no effect on the overall larval morphology and development (Supplementary Figure S2C).

Injection of 2 ng *tgs1* ATG-MO resulted in defects in caudal primary motoneuron (CaP-MN), including axonal truncations and increased terminal branching. 10% of the CaP-MNs exhibited truncated axonal projections and 25% showed increased terminal branching (Figure 3A, B). Negligible CaP-MN defects were observed in uninjected larvae and in larvae injected with a non-targeting MO (Figure 3A, B). 1.5 ng *tgs1* Sp-MO injection led to 15% CaP-MNs with axonal truncations and 35% with increased terminal branching (Figure 3B). Notably, these results recapitulate the motoneuron defects observed in *smn* morphants (36,70,71).

**Figure 3.**
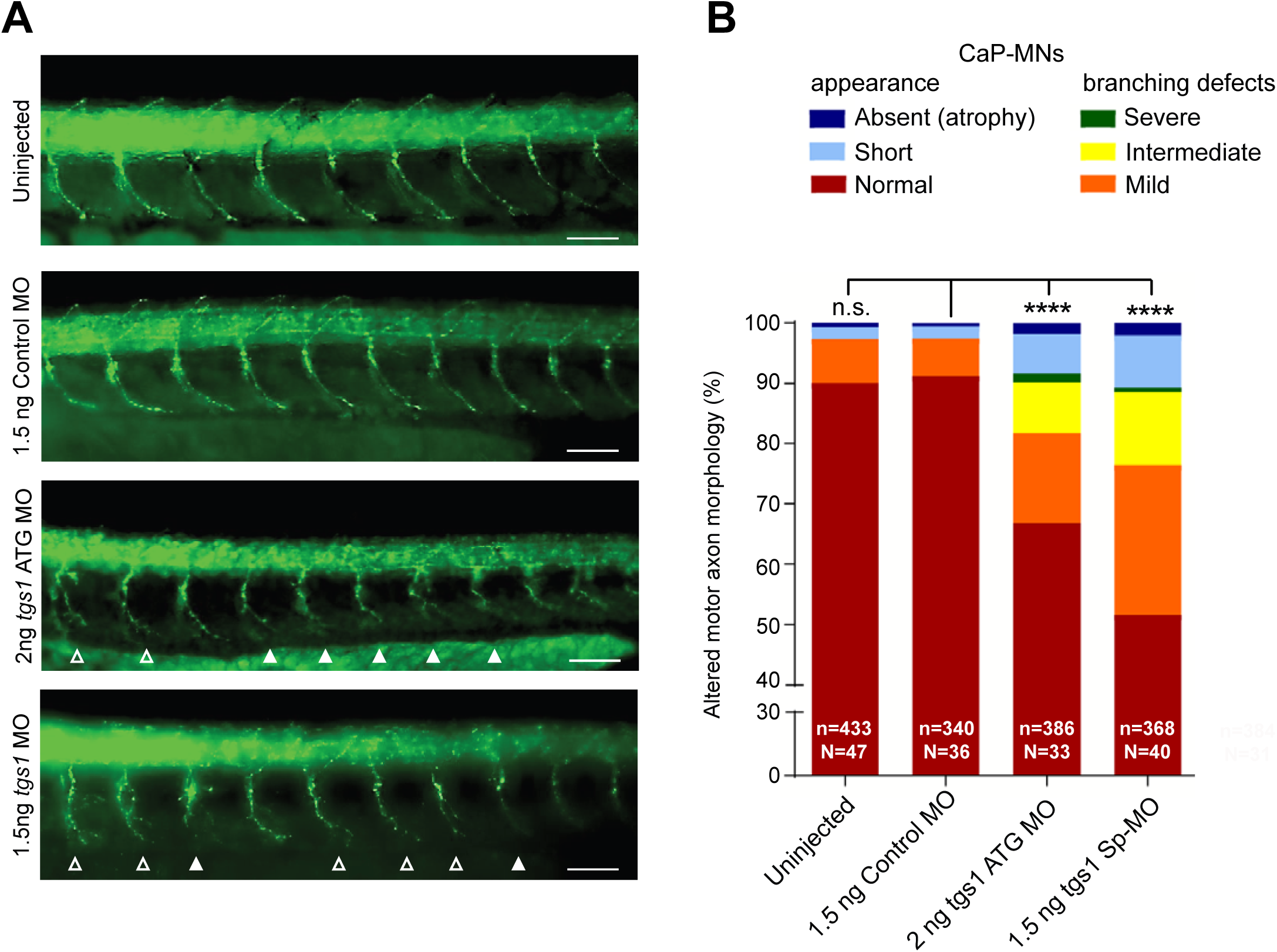
*Tgs1* downregulation in zebrafish leads to caudal primary motor neurons defects. **(A)** Lateral views of whole-mount embryos immunostained with a synaptotagmin antibody (Znp1) that labels the caudal primary motor neurons (CaP-MN). Embryos were untreated or injected with the indicated morpholinos (MO). Injection of either *tgs1* ATG-MO or *tgs1* Sp-MO results in truncated or absent motor axons (solid arrowheads) and terminally-branched axons (open arrowheads). Scale bar, 25 µm. See also Supplementary Figure S2. **(B)** Based on overall appearance, CaP-MNs were classified as: normal, short (truncated axonal projection) or absent (axonal atrophy). Based on terminal branching, axons were classified as normal, mild (branching ventral from midline), intermediate (2-3 or more branches at ventral or midline) or severe (>3 branches ventral or dorsal from midline). Zebrafish larvae injected with 1.5 ng of *tgs1 Sp-*MO or 2 ng *tgs1* ATG*-*MO displayed CaP-MN defects compared to control (non-targeting) MO and uninjected fish. Results are percentages from 3 independent experiments (n = axons analysed; N= animals tested. ****, p<0.0001 Chi-square test); n.s. not significant. See also Supplementary Figure S2D.

We also attempted to perform rescue experiments by co-injecting FLAG-tagged human *TGS1* mRNA and the *tgs1* Sp-MO. Co-injection of *tgs1* Sp-MO and 100 pg of *FLAG-hTGS1* mRNA resulted in a significant rescue of the neurological phenotype (Supplementary Figure S2 D). However, injection in embryos of *FLAG-hTGS1* mRNA (100 to 400 pg) caused developmental defects in larvae (Supplementary Figure S2 C), preventing a firm conclusion about the rescue capacity of human *TGS1.* Collectively, these results show that TGS1 plays a role in motoneuron development in zebrafish.

### TGS1 deficiency impairs 3’ processing of snRNAs in human cells

Given the similarity between the phenotypes elicited by TGS1 and SMN depletion, we asked whether TGS1 loss affects snRNP biogenesis (72). To address this, we used two HeLa clones carrying CRISPR/Cas9-induced mutations in *TGS1* (*TGS1 M1*, *M2*) (45), which decrease TGS1 expression to less than 10% of the normal level but do not affect the expression of the SMN protein (Figure 4A and Supplementary Figure S3C). We have previously shown that both mutant lines exhibit a rather diffuse distribution of the Cajal body (CB) marker coilin (45) and fail to accumulate SMN in CBs. This defect is rescued by the expression of FLAG-TGS1 (*TGS1 M1R* cells, Figure 4A and Supplementary Figure S3 A, B). To ascertain whether *TGS1* deficiency affects snRNA expression, we performed RT-qPCR on total RNA from control cells, *TGS1* mutant cells, and *TGS1* mutant cells bearing a FLAG-*TGS1* rescue construct. There were no significant differences in the abundance of different snRNAs, with minor changes likely reflecting clonal variability (Figure 4B).

**Figure 4.**
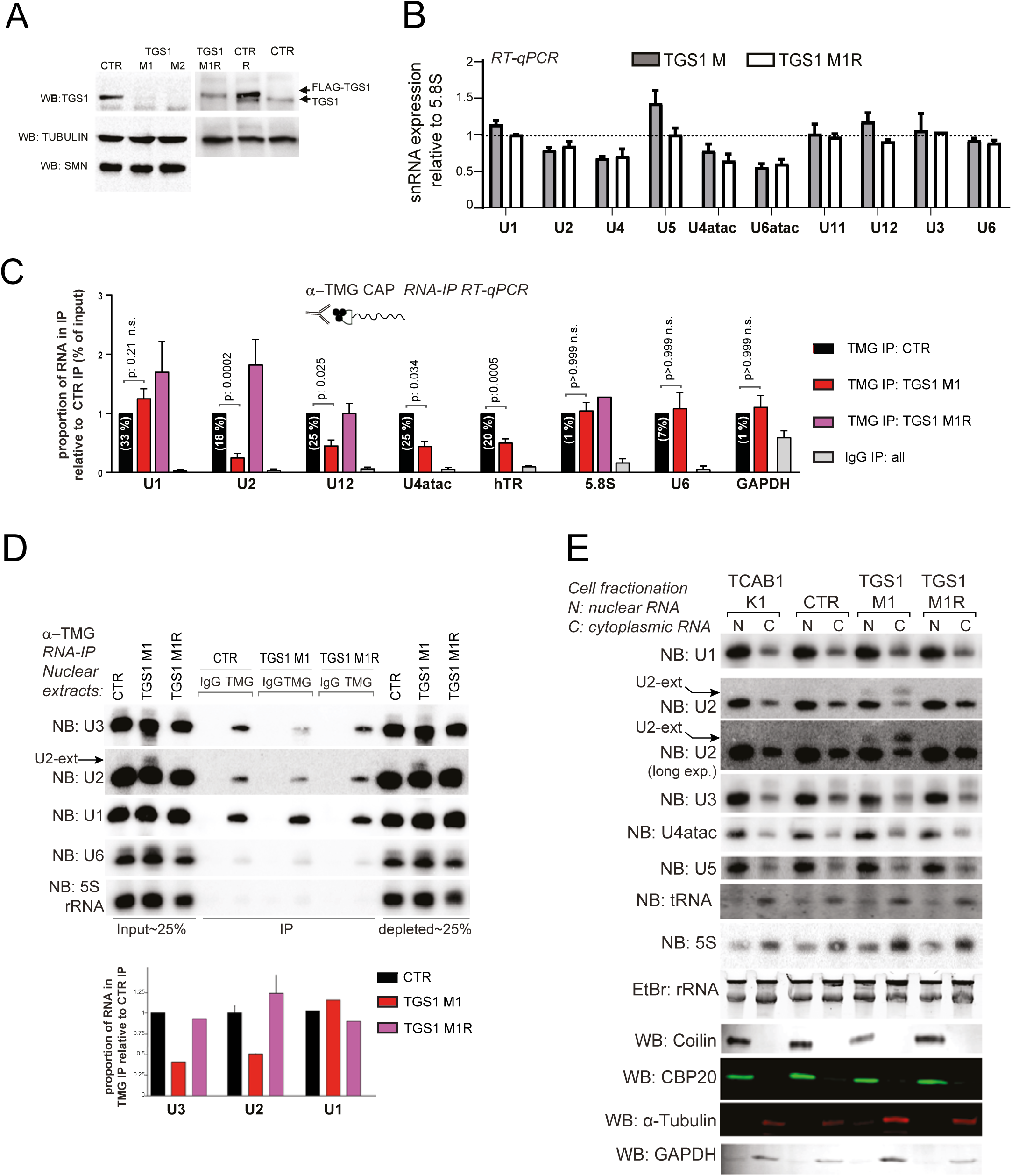
TGS1 deficiency affects maturation of U2 snRNAs in human cells. **(A)** Western blotting (WB) showing TGS1 expression in two independent CRISPR clones of HeLa cells (M1, M2). CTR, parental cell line; *TGS1 M1R* and *CTR R*, are a *TGS1 M1* and a parental line stably expressing FLAG-TGS1. Tubulin is a loading control. The abundance of the SMN protein is not affected by mutations in *TGS1* (see also Supplementary Figures S3C and S6B). **(B)** RT-qPCR showing the abundances of the indicated snRNAs in *TGS1 M1*, *TGS1 M2* and *TGS1 M1R* cells. Data from three independent experiments are relative to parental HeLa cells (set to 1) and normalized to 5.8S rRNA. The U3 snoRNA (U3) and the uncapped U6 and U6atac snRNAs are controls. No statistically significant differences were observed between *TGS1 M* and *TGS1 M1R* cells (error bars: ± SEM; p values: all >0.05; two-way ANOVA with Sidak’s multiple comparisons test) **(C)** Total RNA from control (CTR), *TGS1 M1* and *TGS1 M1R* cells was precipitated with the R1131 anti-TMG CAP antibody (50,73,74) (TMG IP) or control IgGs (IgG IP). Histogram bars represent the fold enrichment of the indicated transcripts in RNA IP (RIP) eluates (relative to input), determined by RT-qPCR and normalized to TMG IP from control cells (set to 1). Numbers in parentheses represent the percentage of RNA in IP eluates relative to input. IgG IP bar: values of IgG IPs for the three cell types were pooled into a single bar. MMG-capped GAPDH mRNA, the uncapped 5.8S and U6 snRNA are negative controls. Data are from three to five independent experiments. p value: two-way ANOVA with Sidak’s multiple comparisons test. **(D)** RIP was performed with the R1131 anti-TMG antibody (TMG) or control IgG on nuclear extracts from control (CTR), *TGS1 M1* and *TGS1 M1R* cells. Membranes were probed for U3 snoRNA, U2 and U1 snRNAs. 5.8S and U6 snRNA that lack a TMG cap, are negative controls. Histogram bars represent the quantification of the U2, U1 and U3 RNAs, by densitometric analysis. Data are from three biological replicates. Note the presence of U2 snRNA precursors (arrow) in nuclear extracts from *TGS1* M1 cells, which display reduced hypermethylation of U2 snRNA and U3snoRNA, but not of U1 snRNA. **(E)** Representative Northern blots showing U2 snRNA precursors (arrow) in nuclear and cytoplasmic fractions. RNA was purified from nuclear and cytoplasmic extracts from: *TCAB1 K1* HeLa mutant cells (this cell line, described in (48) has reduced CBs and was used as a control); CTR, parental HeLa cells; *TGS1 M1* and *TGS1* M1R cells. Membranes were probed for the U1, U2, U4atac, U5 snRNA and the U3 snoRNA. 5S rRNA and tRNA-Arg were used as loading controls. Et Br rRNA staining is a loading control. Western blot with Coilin, CBP20, alpha-tubulin and GAPDH were used as loading controls, indicating absence of nuclear contamination in cytoplasmic fractions and viceversa.

Next, to determine the effects of *TGS1* deficiency on cap hypermethylation of different snRNAs, we performed RNA-IP on total RNA, with the anti TMG antibody R1131 that specifically recognizes the TMG cap (50,73,74). Quantitative RT-PCR (RT-qPCR) on RNA precipitated with the TMG antibody showed that in the U2, U12 and U4atac snRNAs, but not in the U1 snRNA, the proportion of TMG-capped molecules was more abundant in control cells than in TGS1 mutant cells (Figure 4C). The U2 snRNA was the most affected by TGS1 deficiency, as TMG capped U2 snRNA molecules in *TGS1 M1* cells were reduced 4-fold compared to control. TMG-capped U4atac and U12 snRNA molecules were reduced by 50% in *TGS1 M1* cells. In contrast, the abundance of TMG capped U1 snRNA was not reduced in *TGS1 M1* cells. As a positive control for these experiments, we used telomerase RNA (hTR). The abundance of TMG-capped hTR was reduced by 50% in *TGS1 M1* cells (Figure 4C), consistent with our previous work showing that TGS1 hypermethylates hTR and downregulates its abundance (45). The GAPDH, 5.8S and U6 RNAs were used as negative controls as GAPDH mRNA has an MMG cap that is not recognized by the anti-TMG antibody, the 5.8S rRNA is normally uncapped and the U6 snRNA has a γ-monomethyl cap. As expected, these RNAs crossreacted very poorly with the anti-TMG antibody and were precipitated in similar proportions from *TGS1 M1* and control cells (Figure 4C). RNA-IP with the anti TMG antibody, performed on nuclear RNA fractions and followed by Northern Blotting (NB), confirmed that, compared to control and *TGS1 M1R* cells, *TGS1 M1* cells exhibit a reduction in the TMG capped U2 snRNA and U3 snoRNA, but not in the U1 snRNA (Figure 4D). This experiment also showed that nuclear extracts of *TGS1* M1 cells are enriched in slower migrating U2 species (Figure 4D), which are likely to correspond to U2 precursors carrying genome-encoded extensions at their 3’ ends (11, 21).

Next, we analyzed by NB the distribution of the extended snRNA species in nuclear and cytoplasmic fractions of both control and *TGS1* mutant cells. Extended RNA precursors were detected for U2 snRNAs but not for U1, U3, U4atac, and U5 snRNAs. U2 snRNA precursors were clearly enriched in both nuclear and cytoplasmic RNA of *TGS1 M1* and M2 cells, compared to TGS1-proficient cells (Figure 4E and Supplementary Figure S4A, B). Nuclear or cytoplasmic accumulations of extended U2 precursors were not enriched in cells carrying mutations in *TCAB1* (48) that disrupt Cajal body stability (Figure 4E). Immunostaining of *TGS1 M1* cells with antibodies against the TMG cap and the SmB protein showed a strong reduction in the speckled nuclear distribution of snRNPs compared to control or *TGS1 M1R* cells (Supplementary Figure S4C). In addition, after immunostaining with these antibodies, *TGS1 M1* cells showed a diffuse cytoplasmic halo that was not observed in control or *TGS1 M1R* cells, suggesting that TGS1 deficiency leads to retention of some snRNPs in the cytoplasm. Consistent with this finding, single molecule FISH detected cytoplasmic retention of U2 species in *TGS1 M1* cells but not in TGS1-proficient cells (Supplementary Figure S4D). In contrast, FISH analysis showed that the nuclear localization of U1 snRNA is unaffected by TGS1 deficiency (Supplementary Figure S4D). These results indicate that TGS1 deficiency causes aberrant accumulation of unprocessed U2 precursors that might interfere with spliceosome activity, as occurs in yeast (75).

We characterized the 3’ ends of the pre-U2 molecules in total RNA by 3’RACE and Next-Gen sequencing as previously described (47, 76). In control cells, 90% of the reads mapped to the mature forms of U2, most of which contained 188 nt plus a 3’ adenosine added post-transcriptionally (≤189 nt), in agreement with previous results (3, 77). The remaining 10% of the U2 molecules had extended tails including genome-encoded nucleotides at their 3’ ends (Figure 5A, B). *TGS1* mutants displayed a 4-fold higher proportion of 3’ extended U2 molecules (41%) compared to control (10%), with an average tail length of 12 nt (Figure 5A, B). In *TGS1 M1R* cells, the abundance of the extended U2 molecules was reduced from 41% to 20%, showing rescue of the mutant phenotype (Figure 5A). 3’RACE and sequencing performed on nuclear and cytoplasmic RNA confirmed the data obtained by NB (Figures 4D, E, 5C, D). In the nuclear RNA fractions, the abundance of the U2 extended species was 18% in *TGS1* mutant cells, 7% in control cells and 8% in *TGS1M1R* cells (Figure 5C). In the cytoplasmic fractions, the extended U2 molecules were 52% in *TGS1* mutant cells, 13% in control cells, and 20% in *TGS1M1R* cells (Figure 5C). *TGS1* mutant cells showed no variations in the proportion of mature U2 molecules carrying a 3’ untemplated adenosine compared to controls; in both cases the 189/188 ratio was approximately 80%. Importantly, the finding that the nuclei of *TGS1* mutant cells contain more extended U2 molecules than control nuclei (Figures 4D, E and 5C) suggests that a fraction of these unprocessed species completes the cytoplasmic assembly pathway and is imported in the nucleus.

**Figure 5.**
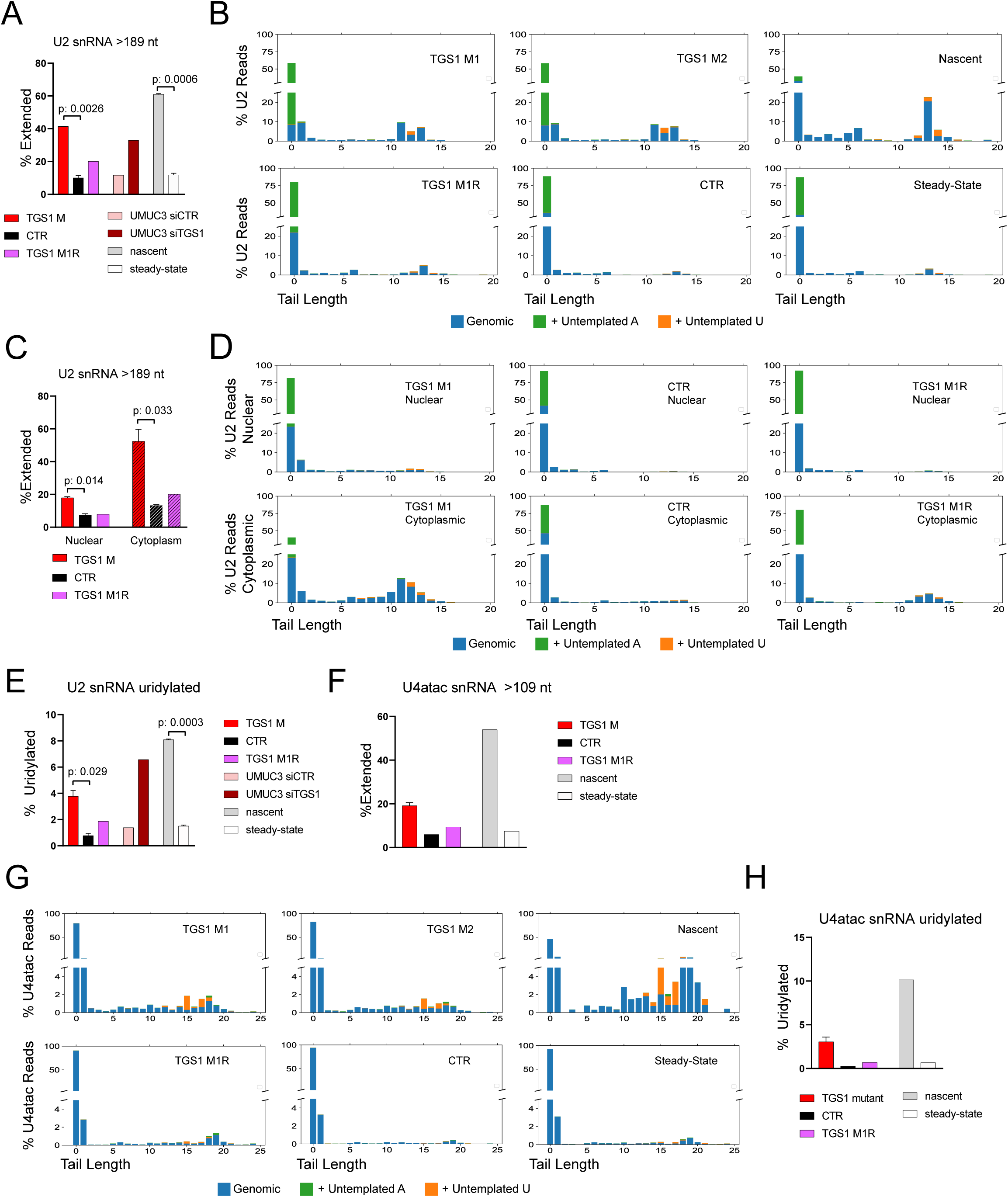
TGS1 loss leads to an accumulation of 3’extended U2 and U4atac snRNAs in human cells. **(A)** 3’ RACE and sequencing on total RNA showing that *TGS1* mutant HeLa cells accumulate more U2-extended snRNA molecules than either parental (CTR) or *TGS1 M1R* cells. UMUC3 cells treated with TGS1 siRNAs (siTGS1) also accumulate more U2- extended snRNAs than cells exposed to non-targeting siRNAs (siSCR). Nascent and steady-state RNA fractions were purified from control HeLa cells (see Methods). Extended U2 molecules carry extra 3’ sequences of templated and /or untemplated nucleotides. Average reads per sample > 78k ± 41k. Data are mean values of two independent clones + SEM. p values: two-sided Student’s t test. **(B)** Plots showing the percentage of the different sequence reads of U2 snRNAs in the same HeLa cell samples as in A. Numbers on the x-axis: additional nucleotides beyond the 189 nt of the mature form. Position 0 includes mature U2 snRNA species of 188 nt or 188 nt plus a post-transcriptionally added A. y-axis: percent of total reads. Blue: genomic-templated nucleotides; green: untemplated adenosine; orange: untemplated uridine. **(C)** Characterization of U2 snRNA molecules from nuclear and cytoplasmic fractions from *TGS1 M1*, *TGS1 M2*, *TGS1 M1R* and control (*CTR*) cells. **(D)** Tail length and composition U2 snRNAs in the samples shown in (C). y-axis: percent of total reads. **(E)** Percentages of U2 molecules with tails ending with post-transcriptionally added uridines in the HeLa cell RNA samples described in A. Total reads per sample >75k. **(F)** Percentages of 3’ extended U4atac snRNAs in the RNA samples described in A. Average reads per sample > 106k ± 21k (Nascent:>11k). **(G)** Tail lengths and composition of the 3’ ends of U4atac snRNAs in *TGS1 M1*, *TGS1 M2*, *TGS1 M1R* and control (*CTR*) cells. Numbers on the x-axis: additional nucleotides beyond the 130nt of mature U4atac (position 0). y-axis: percent of total reads. **(H)** Percentages of U4atac snRNA molecules with extended tails ending with untemplated uridines. Total reads per sample >75k.

To assess whether TGS1 deficiency affects processing of the U2 precursors in a different cell type, we knocked down *TGS1* by RNAi in UMUC3 cells, which showed a strong reduction in the TGS1 protein and defective CBs (Supplementary Figure S3D, E). 3’ RACE and sequencing on total RNA revealed that in *TGS1*-RNAi cells the proportion of extended molecules is 3-fold higher than in cells treated with control siRNA (Figure 5A).

To confirm that the extended species are precursors of U2 molecules, we performed metabolic labeling of newly transcribed RNA with 4-thiouridine for 4 hours in control HeLa cells, and affinity-purified this nascent RNA population with thiol-specific biotinylation and streptavidin-coated magnetic beads (47). Unlabeled RNAs (steady-state RNA) recovered after this treatment, are enriched in mature RNA species while labelled RNAs are enriched in short-lived RNA precursors. A comparison of the sequence profiles showed that extended-U2 molecules are 5-fold more abundant in nascent RNA samples (59% of total U2 reads) than in steady-state RNA (12%; Figure 5A, B), indicating that the extended U2 species are immature U2 precursors, not yet processed at their 3’ ends.

In total RNA from control cells, a small fraction of the U2 reads are extended molecules carrying non-templated adenosines (0.2%) or uridines (0.6%) added post-transcriptionally at their 3’ ends. The acquisition of an extra A or U is specific for the extended molecules with a mean tail length of 12 nucleotides. In *TGS1* mutant cells, the fraction of extended U2 molecules incorporating extra uridines is 6 times and 2 times higher than in control and *TGS1 M1R* rescued cells, respectively (Figure 5B, E). A 5- fold increase in uridylated molecules is also observed in *TGS1* RNAi UMUC3 cells compared to cells treated with nontargeting dsRNA (Figure 5E). These data suggest that TGS1 affects not only 3’ trimming but also the profile of post-transcriptional modifications on the U2 precursors. Interestingly, oligo-U extended molecules are 5-fold more abundant in nascent RNA than in mature RNA, indicating that these modifications represent an intermediate step in the 3’ processing of U2 precursors (Figure 5E).

We also analyzed the 3’ sequences of the U1and U4atac snRNAs. In both control and *TGS1* mutant cells 98% of U1 snRNA molecules were 164 nt long and less than 1% of these molecules carried untemplated adenosine or uridine residues at their ends, indicating that in the *TGS1* mutant clones, neither cap hypermethylation of U1 snRNAs, nor their biogenesis is affected (Supplementary Figure S5A). In contrast, *TGS1* mutant cells were enriched in U4atac extended forms showing a mean tail length of 10 nt; these forms were 19% of total reads in *TGS1* mutants, 6% in control cells, and 9% in rescued cells (Figure 5F-G). Moreover, in *TGS1* mutant cells 3% of U4atac molecules incorporated additional uridines, compared to 0.25% in control cells and 0.7% in rescued cells (Figure 5H).

Collectively, these results indicate that TGS1 depletion results in defective hypermethylation of some snRNA species, which largely correlates with the accumulation of extended U2 and U4atac snRNA molecules. These snRNAs differ in the type and proportion of 3’ end extensions, most likely reflecting diverse dynamics in their 3′ end processing. The involvement of TGS1 and cap hypermethylation in 3′ maturation of snRNAs is further supported by the observation that in our mutant background both cap hypermethylation and 3′ end processing of U1 snRNA are unaffected by TGS1 deficiency.

### Tgs1 controls 3’ processing of snRNAs in Drosophila

To ask whether the role of TGS1 in snRNA 3’ processing is conserved in flies, we exploited *dTgs1^R1^*, a null CRISPR/Cas9-induced allele, which in homozygosity causes death in late second instar larvae (40). We also used *dTgs1^R1^*/*dTgs1^CB0^* heterozygous flies that die as third instar larvae. The lethality of both mutants is rescued by ubiquitous expression of either a *dTgs1* or a human *TGS1* transgene (40, 42).

We characterized the 3’ ends of U2 snRNAs by 3’ RACE and RNA sequencing on total RNA from wild type larvae (*CTR*), *dTgs1^R1^* homozygous larvae and *dTgs1^R1^*/*dTgs1^CB0^* larvae; as control, we also examined mutant larvae constitutively expressing *dTgs*1. *dTgs1* mutant larvae showed an increase in 3’ extended U2 species compared to either wild type larvae or larvae bearing the rescue construct (Figure 6A). 95% of these extended U2 species displayed one additional uridine at the 3’ end (Figure 6B). The remaining 5% of the extended molecules included species containing longer tails incorporating both genome-templated and untemplated A or U nucleotides. Thus, TGS1 is required for proper 3’ processing of U2 snRNAs also in flies.

**Figure 6.**
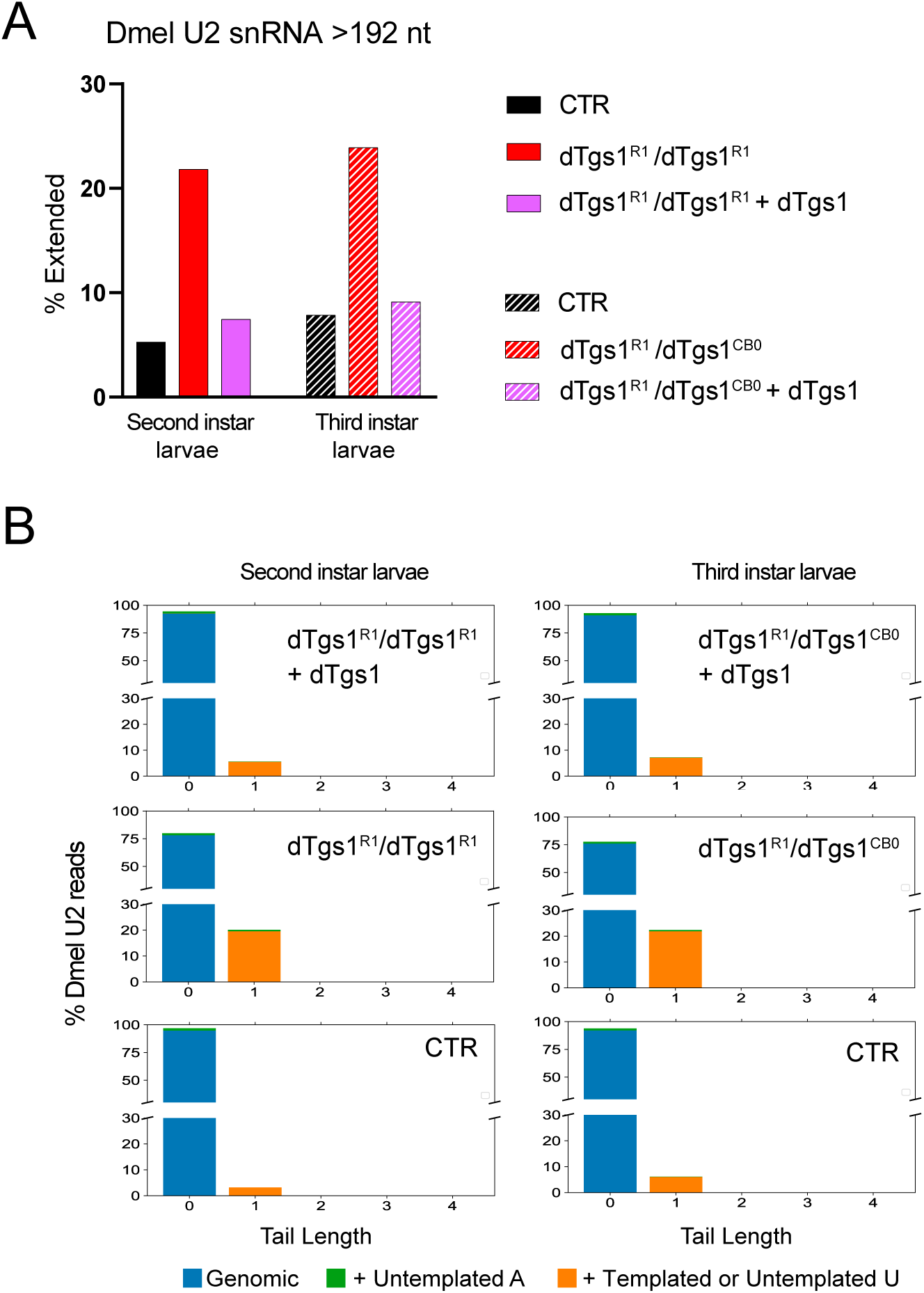
3’ extended U2 snRNA molecules accumulate in *Drosophila dTgs1* mutants. **(A)** Percentage of extended U2 snRNA molecules determined by 3’ RACE and sequencing on RNA from equally aged second or third instar mutant larvae, rescued mutant larvae (bearing a *dTgs1* construct), and wild type larvae (CTR). The most abundant population of U2 snRNAs is 192 nt long. Extended U2 snRNAs are longer than 192 nt and incorporate templated or untemplated nt at their 3’ ends. Total reads > 2900. **(B)** Tail length and composition of the 3’ ends of U2 snRNAs for the second and third instar larvae described in A. Numbers on the x-axis: additional nucleotides beyond the mature form of 192 nt (position 0). y-axis: percent of total reads.

In contrast, an analysis of the U1 snRNAs showed that they are substantially unaffected by *dTgs1* mutations, consistent with the results obtained with human cells. In mutant and wild type larvae, extended species represented about 1,5% and 0.5% of total U1 RNAs, respectively.

### Aberrant transcripts accumulate in both *TGS1* and *SMN* mutant cells

Studies conducted in different systems demonstrated that SMN deficiency causes profound perturbations in the transcriptome (78–83). To explore the consequences of TGS1 loss on mRNA splicing and gene expression, we performed deep sequencing on total RNA extracted from *TGS1* mutant HeLa cells, and from two independent *SMN* mutant clones derived from the same HeLa cell line used to generate *TGS1* mutant cells. These lines, designated as *SMN C1* and *SMN C2*, carry CRISPR-induced mutations in the sixth exon of the gene and produce mutated versions of the SMN protein, which are expressed at reduced levels, compared to full length SMN (Supplementary Figure S6A, B). These lines have normal amounts of the TGS1 protein (Supplementary Figure S6B) and do not exhibit significant changes in viability, compared to the HeLa parental cell line (CTR) and *TGS1 M1* mutant cells (Supplementary Figure S6C). *SMN C1* has a small deletion in the SMN proline rich domain and *SMN C2* carries frameshift mutations (see Supplementary Figure S6A). The SMN proteins encoded by the *SMN C2* cells have a lower MW compared to the wild type protein. The *SMN C1* and *SMN C2* mutant cells are defective in CB formation and exhibit fewer SMN-enriched CBs compared to control cells (Supplementary Figure S6D, E).

Paired-ended Illumina sequencing (160 million reads/sample) was performed on RNA isolated from *TGS1* mutant clones (*M1, M2*), *SMN* mutant clones (*C1, C2*), and parental HeLa cells (*CTR*). Independent mutant clones and controls were clustered based on both transcript and gene expression profiles (Supplementary Figure S7A-C). We used Kallisto (84) and Sleuth (55) for transcript quantification and differential analysis, respectively (see Methods). *TGS1* and *SMN* mutant cells showed a differential expression of both annotated and unannotated transcripts compared to controls (reconstructed by Scallop (54); Figure 7A, B; and Supplementary Table S1). Our analysis revealed significant changes in the expression levels of 3084 and 351 transcript isoforms in TGS1 and SMN mutant cells, respectively (Figure 7A and B; Supplementary Table S1). We found changes in the expression levels of many unannotated transcripts (1080/3084 for TGS1 and 158/351 for SMN mutants); 48 of these novel isoforms were found to be affected in both mutant cell types (Figure 7C; Table S1). The number of differentially expressed (DE) transcripts was higher in *TGS1* mutant cells than in *SMN* mutant cells (Figure 7A-C), likely reflecting the higher degree of functional inactivation of *TGS1* compared to *SMN* (Figure 4A and Supplementary Figure S6A, B).

**Figure 7.**
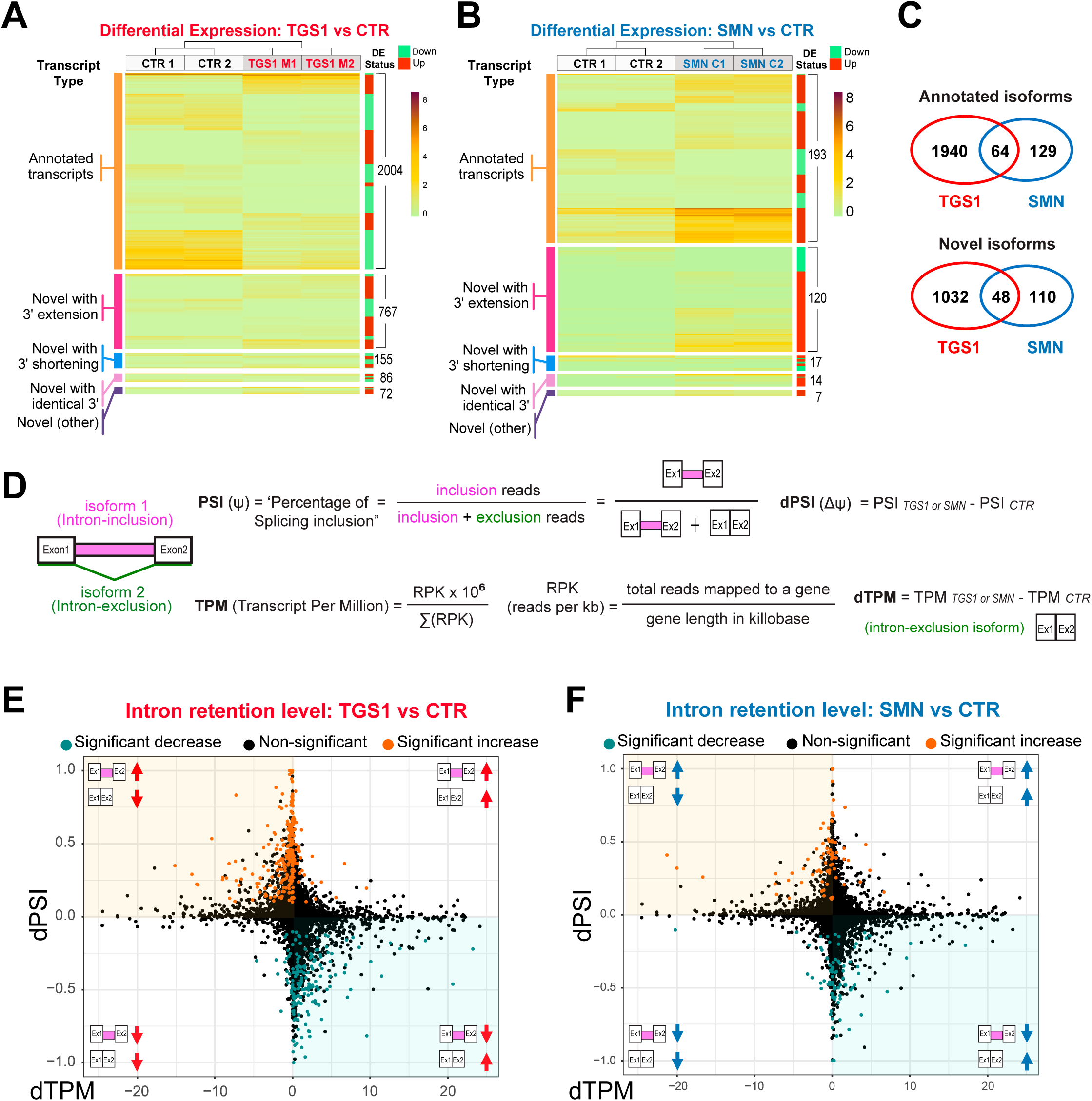
Mutations in *TGS1* and *SMN* cause global changes in RNA expression and splicing. **(A, B)** Heatmaps depicting the expression levels (TPM values estimated by Kallisto) for differentially expressed (DE) transcripts in *TGS1 M*1 and *TGS1 M2* (A), or *SMN C1* and *SMN C2* (B) mutant cells, compared to parental HeLa cells (CTR). Transcripts are classified into 5 types: annotated transcripts per GENCODE (orange); novel transcripts with extended 3’ (magenta) or shortened 3’ (blue); novel transcripts with annotated identical 3’ (pink); and other (intergenic and opposite-strand) novel transcripts (purple). Transcripts within each group are ranked by unsupervised clustering. Total transcript number for each group, and the DE status for each transcript (either up- or down- regulated in mutant cells) are annotated by the green/red sidebar to the right (see also Supplementary Table S1). **(C)** Venn diagram from data in A and B, showing the significant number of shared differentially expressed transcript isoforms (both annotated and unannotated) between *TGS1* and *SMN* mutant cells. See also Supplementary Table S1. **(D)** Methodological approach used for quantification of intron retention (IR) levels (see also Methods). **(E, F)** Scatter plots showing for each IR event between mutant (*TGS1 M1* and *M2* or *SMN C1* and *C2*) and CTR cells the differential PSI (dPSI) against the differential expression level (dTPM) of the major transcript (intron exclusion). The significant IR events are color-coded in the plots, with significantly upregulated events colored in orange and significantly downregulated in teal.

The number of annotated and unannotated transcripts that were found to be differentially expressed in both *SMN* and *TGS1* mutant cells were significantly greater than would be expected by chance (p-value < 10e-22, hypergeometric test, 110400 total annotated transcripts, 28100 total unannotated transcripts with computed p-value by Sleuth) (Figure 7C; Supplementary Table S1, see Methods). Notably, 71% and 76% of the differentially expressed unannotated transcripts respectively found in *TGS1* and *SMN* mutant cells, showed an increased number of reads for intergenic regions compared to controls (Figure 7A, B; Supplementary Table S1). These 3’ elongated transcripts extended beyond the 3’ end of the gene and sometimes included the coding region of the next gene, likely reflecting an altered transcription termination. Most of the DE transcripts were also identified using an independent analysis workflow (54, 57) (Supplementary Tables S3-S5). The mutant cells also exhibited changes in splicing behavior (Supplementary Figure S8A), with intron retention being the most frequent defect in both *TGS1* (3.5% and 4.5% of annotated and unannotated IR events, respectively) and *SMN* mutants (1.2% and 2.1% of annotated and unannotated IR events, respectively); Figure 7D-F; Supplementary Figure S8A, B and Table S2.

Long read Oxford Nanopore sequencing was performed on the six RNA samples, described above, to gain additional information on the transcripts that accumulate in *TGS1* and *SMN* mutant cells, and to assess the association of splicing defects and 3′ extensions on the same transcript isoform. 72 million total reads were generated, preprocessed and mapped to the human genome. The FLAIR pipeline (59), was employed for splicing junction correction, yielding 24,7 million corrected multi-exonic reads that were used for transcriptome reconstruction (see methods and Supplementary Figure S9). 55,815 transcripts, 41,565 of which not present in the reference annotation, were identified and quantified using FLAIR; differential splicing between mutant and control samples was assessed using SUPPA2 (Figure 8 A-C, Supplementary Figure S10A). The number of shared annotated and unannotated differential splicing events between *SMN* and *TGS1* mutant cells is greater than expected by chance (Figure 8 D). Furthermore, there is a significant overlap between the differential splicing events identified by Illumina and Nanopore sequencing (Figure 8E and Supplementary Table S8), providing further support for the reproducibility of our findings on the global splicing defects caused by TGS1 and SMN depletion. Multi-exonic reads intersecting the last exon of each gene were used to identify loci in which the proportion of readthrough transcripts was significantly different in *TGS1* and *SMN* mutant samples compared to control (Figure 8F and Supplementary Table S9). Interestingly, in both comparisons most of the differential readthrough events consist in a higher proportion of readthrough reads in the mutant, indicating a defective termination leading to the production of longer transcripts. Many of these longer transcripts extended into the downstream gene. For the genes, showing such Preferential Readthrough in Mutant samples, SUPPA-2 was used to determine the number of readthrough and non-readthrough reads harboring an alternative spicing event in the mutant samples. These analyses (Figure 8G and Supplementary Table S10) indicate that in both *TGS1* and *SMN* mutant cells, readthrough reads have a higher probability of containing also an alternative splicing event compared to non-readthrough reads.

**Figure 8.**
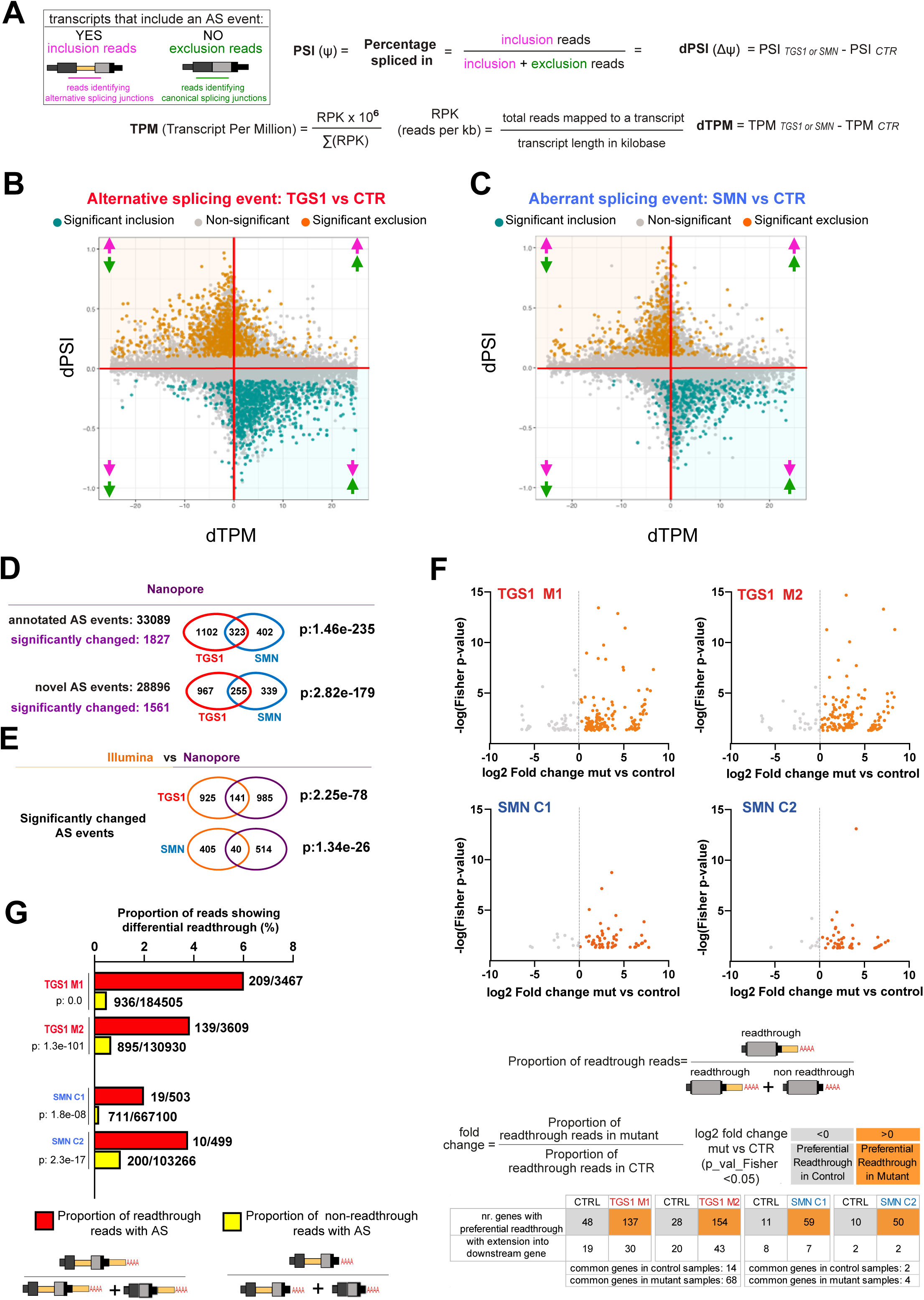
Analysis of the aberrant transcripts that accumulate in *TGS1* and *SMN* mutant cells by Oxford Nanopore sequencing. **(A)** Methodological approach used for transcript quantification and evaluation of transcript differential splicing. **(B,C)** Scatter plots showing, for each AS event between mutant (*TGS1 M1, M2* and *SMN* C1, C2) and parental HeLa cells (CTR), the differential PSI (dPSI) against the differential expression level (dTPM) of the major transcript (that does not contain the AS event). The significant differential splicing events are color-coded in the plots, with significantly included events colored in orange and significantly excluded in teal. Non-significant events are in grey (see also Supplementary Figure S10A for a classification of the AS events and Table S6). The arrows indicate up or downregulation of AS inclusion transcripts (pink) and AS exclusion transcripts (green), as depicted in (A), respectively. **(D)** Venn diagrams showing the number of annotated and novel differential splicing events found in *TGS1 M1, M2* and *SMN C1, C2* mutant cells by Nanopore sequencing. The statistical significance of the intersection was determined via Fisher’s exact test. See also Supplementary Table S7). **(E)** Venn diagram showing the number of the differential splicing events identified by Illumina and Nanopore analysis in *TGS1* and *SMN* mutant cells. The statistical significance of the intersection was determined via Fisher’s exact test. See also Supplementary Table S8. **(F)** Dot plots showing differential readthrough events in mutant cells (orange dots) vs control cells (grey dots). For each readthrough event, the value of the log2(fold change) (mutant vs control) is reported on the x-axis. Statistical significance of the fold enrichment for each event was determined by Fisher’s exact test (p-values on the y axis; p<0.05 was considered significant). The numbers of genes with “Preferential Readthrough in Mutant” (orange) or “Preferential Readthrough in CTR” (grey) are reported below the graphs. See also Supplementary Table S9; see methods. **(G)** Bar plot showing that, in mutant samples, the percentage of reads that carry an aberrant splicing event is higher among readthrough reads compared to non-readthrough reads. Only reads mapping to last exons undergoing “Preferential Readthrough in Mutant” events were used for this analysis. Aberrant splicing was assessed by comparing reads from mutant samples against non-readthrough reads from control samples. The significance of the difference in the proportions of aberrant splicing reads between readthrough and non-readthrough reads was assessed using the chi-square test. See also Supplementary Table S10.

To confirm the results obtained using high-throughput data analysis, we performed targeted validation for 4 transcripts by RT-PCR or RT-qPCR on RNA samples from CTR, *TGS1* M and *TGS1* M1R cells. Amplification products consistent with the predicted novel transcript isoforms were enriched in the two *TGS1* mutant clones compared to both the parental cell line and *TGS1 M1R* cells (Supplementary Figures S11 and S12). Two of these validated novel transcripts were found to be enriched in *TGS1* mutant clones also by Nanopore analysis and are representative examples of readthrough transcripts that extend into the downstream gene. (Supplementary Figures S11B, E).

## DISCUSSION

We have shown that the highly conserved TGS1 hypermethylase localizes to motor neurons in *C. elegans*, and that its deficiency results in morphological and functional abnormalities in neurons from both *C. elegans* and *D. rerio*. Consistent with these defects, *TGS1* deficiency impairs locomotion in worms and wing expansion in flies, which is governed by bursicon-expressing neurons (85). Previous work has also shown that mutations in *Drosophila Tgs1* cause abnormal larval locomotion (40,67,68). The phenotypes elicited by *TGS1* inhibition in flies, worm and zebrafish are highly reminiscent of those caused by loss of function of the *SMN* gene. Importantly, we showed that *TGS1* overexpression ameliorates the neurological phenotypes elicited by the impairment of the *SMN* function in both fly and worm model systems. These findings suggest commonalities in the molecular and cellular effects induced by either *TGS1* or *SMN* loss of function.

Our investigation on the consequences of TGS1 loss in snRNA biogenesis uncovers the involvement of this hypermethylase in 3′ end processing. The HeLa cell model was instrumental to compare the molecular consequences of SMN and TGS1 loss in the same isogenic background. Additionally, we demonstrate that 3′ extended mRNA transcripts accumulate in both *TGS1* and *SMN* mutant cells. Lastly, our analyses of different animal models provide robust evidence that TGS1 together with SMN plays conserved, important activities for neuronal function.

Several studies have linked neurodegeneration to defective snRNA processing, snRNP biogenesis and/or pre-mRNA splicing (3,72,82,86,87). It has been reported that SMN-depleted cells exhibit an enrichment of 3’ unprocessed snRNA precursors resulting from their impaired assembly into snRNPs (11). In a mouse model of SMA, motor neurons display a much greater reduction in the snRNPs levels relative to unaffected spinal cells (88), and their selective death is the result of converging mechanisms of p53 activation driven by dysregulation of distinct mRNA splicing pathways (89–91). It has been also reported that mutations in a single gene of the U2 multicopy cluster (Rnu2-8), result in defective splicing and cause ataxia and neurodegeneration in mice (86). Unprocessed snRNAs accumulate in cells from patients affected by pontocerebellar hypoplasia (PCH7), which is also characterized by motor neuron loss (3). Interestingly, we show that *TGS1* mutant cells accumulate extended snRNA species with post-transcriptionally added adenosines or uridines similar to cells from PCH7 patients (3,92,93).

Our 3′ RACE and sequencing experiments show that TGS1 loss results in the accumulation of untrimmed 3’ extended U2 snRNA molecules, but has no effect on U1 snRNA in both human cells and *Drosophila* lethal mutants. TGS1-deficient human cells also accumulate untrimmed U4atac snRNAs. Interestingly, viable hypomorphic *DrosophilaTgs1* mutants that cause male sterility exhibit a reduction in TMG-capped U1 and U2 snRNAs in testes and a concomitant accumulation of long precursors for most snRNAs (94). Collectively, these results indicate that TGS1-mediated cap hypermethylation affects 3′ end processing of snRNAs.

Immature snRNA precursors produced by different snRNA genes, differ in length (11,21,94–97). In human cells deficient for either TGS1 (this study) or TOE1 (3, 93), the tails of the extended molecules generated by different U snRNA subclasses vary in length, number of templated nucleotides, and number/type of untemplated residues. We found that in TGS1-deficient human cells and flies the U2 snRNAs acquire different 3′ extended structures. While the majority of extended human U2 snRNAs has templated tails with or without an untemplated U residue, *Drosophila* U2 molecules preferentially gain untemplated uridines. Uridylation is thought to recognize misprocessed and/or low quality snRNPs and target them for degradation (93,98–100). The variation in the 3’ ends of snRNAs may reflect differences in promoter sequences (101), gene bodies, or 3′ regions, which are crucial determinants for the cleavage and maturation of the snRNA precursor transcripts (102). The different susceptibilities of individual snRNAs to loss of maturation factors might also depend on the specificity of these factors in 3′ end processing and to their residual levels. It is possible that TGS1 hypermethylates different snRNAs with different efficiencies and that our experimental conditions did not allow detection of differences in the hypermethylation level of U1 snRNA.

There is also evidence suggesting that the U1 snRNAs behave differently from the other U snRNA subclasses. Recent work has revealed that 3′ end processing of some snRNAs is a biphasic process that takes place both before and after the assembly with the SMN complex (93). Human U1 has a peculiar Sm core assembly pathway and differs from the other snRNAs in the affinity for SMN-Gemin5 (11, 103). These features suggest a specific regulation of the biogenesis of the U1 snRNP, which is the most abundant snRNP, playing roles in both splicing and polyA site selection (telescripting) (104). Thus, the finding that the U1 snRNAs are not affected by loss of TGS1 might depend on several factors, including the specificity of the TGS1 enzymatic activity and the peculiar maturation pathway of the U1 snRNPs.

TGS1 deficiency results in extensive alterations in the human transcriptome, including changes in the efficiency of intron removal and accumulation of transcripts with 3’ extensions spanning intergenic regions, and often incorporating exons of adjacent genes. Similar alterations are also observed in the transcriptome of *SMN* mutant cells. Long-read ONT sequencing confirmed that *TGS1*- and *SMN*- deficient cells accumulate mRNAs that carry both splicing and 3′ end cleavage defects. Many of these are chimeric transcripts, in which the exon of a gene is fused to exons of the downstream gene. Recent work has suggested that SMN directly mediates proper transcription termination by favoring polymerase II release (105), and SMN deficiency has been linked to accumulation of R-loops and DNA damage (80, 106). Accumulation of aberrant read-through transcripts is consistent with these results and may contribute to the neurodegenerative defects observed in *TGS1* and *SMN* mutant models.

Previous work has shown that U2 snRNP alterations caused by deficiency of the SF3B1 protein result in splicing defects and transcription proceeding beyond the canonical polyadenylation (poly-A) sites, leading to severe pathological consequences (59, 107). In addition, it has been shown that U1 snRNPs prevent premature transcription termination before the last exon by inhibiting 3′ end cleavage at cryptic, early polyadenylation sites (104). The results of our Long-read ONT sequencing of transcripts from TGS1- and SMN- deficient cells support the hypothesis that at least a fraction of the aberrant readthrough mRNAs observed in *TGS1* and *SMN*-mutant cells are caused by a primary defect in splicing, and consolidate the notion that reduced efficiency of co-transcriptional splicing impairs correct transcription termination (108, 109) However, our observations do not exclude the possibility that TGS1 cooperates with SMN in transcription termination and that its loss directly contributes to the formation of long readthrough transcripts.

The characterization of transcriptional defects in *TGS1*- and *SMN*-deficient cells was carried out on steady-state poly-A transcripts and therefore the occurrence of non-polyadenylated and unstable species that are subjected to clearance by the RNA surveillance machineries may have been overlooked. It has been reported that inverted Alu repeats promote the expression of circular RNAs from the *SMN* locus (110, 111). In our sequencing datasets we did not find aberrant transcripts arising from the *SMN* loci, however, it is possible that transcriptional readthrough of the *SMN* gene might enhance the expression of SMN circular RNAs (110). Further analyses with methods that allow the isolation of non-polyadenylated transcripts as in (112) and (113) and by long-read sequencing of nascent RNA will likely expand the repertoire of RNAs affected by TGS1 and SMN depletion.

Collectively, our data support a model where TGS1 and SMN play closely related functions in preventing neurodegeneration (Figure 9A, B). TGS1 plays functions in snRNA 3′ maturation and its loss affects the accuracy of both pre-mRNA splicing and transcription termination. SMN promotes assembly of snRNA into snRNPs and is essential for splicing and efficient transcription termination (82, 105). Thus, loss of either protein leads to substantial transcriptome alterations. These alterations may be neurotoxic by directly affecting specific transcripts important for neuronal survival and function, a scenario supported by increasing experimental evidence (89-91,114). In addition, reduced efficiency of splicing and transcription termination may be a source of transcriptional stress that results in genome-wide accumulation of R-loops and DNA damage, which may also affect neuronal survival (115). Future studies on ultra-fine characterization of aberrant mRNA molecules and their effects in neurons are needed to support this possibility, which has strong relevance not only for SMA but also for other neurodegenerative disorders. Importantly, we found that TGS1 overexpression can partially rescue the neurological phenotypes caused by SMN depletion in the fly and worm models. Moreover, previous work has shown that TGS1 directly interacts with the SMN protein in human cells (14), and that *Drosophila* Tgs1 physically associates with most components of the SMN complex *in vivo* (40). Thus, the physical interaction between TGS1 and SMN and the similar phenotypes elicited by their deficiency suggest a functional interaction between these proteins. We hypothesize that TGS1 overexpression in *SMN*-depleted cells might enhance snRNP biogenesis and possibly improve the fidelity of splicing and transcription termination, functionally compensating at least in part for defects caused by SMN deficiency (Figure 9).

**Figure 9.**
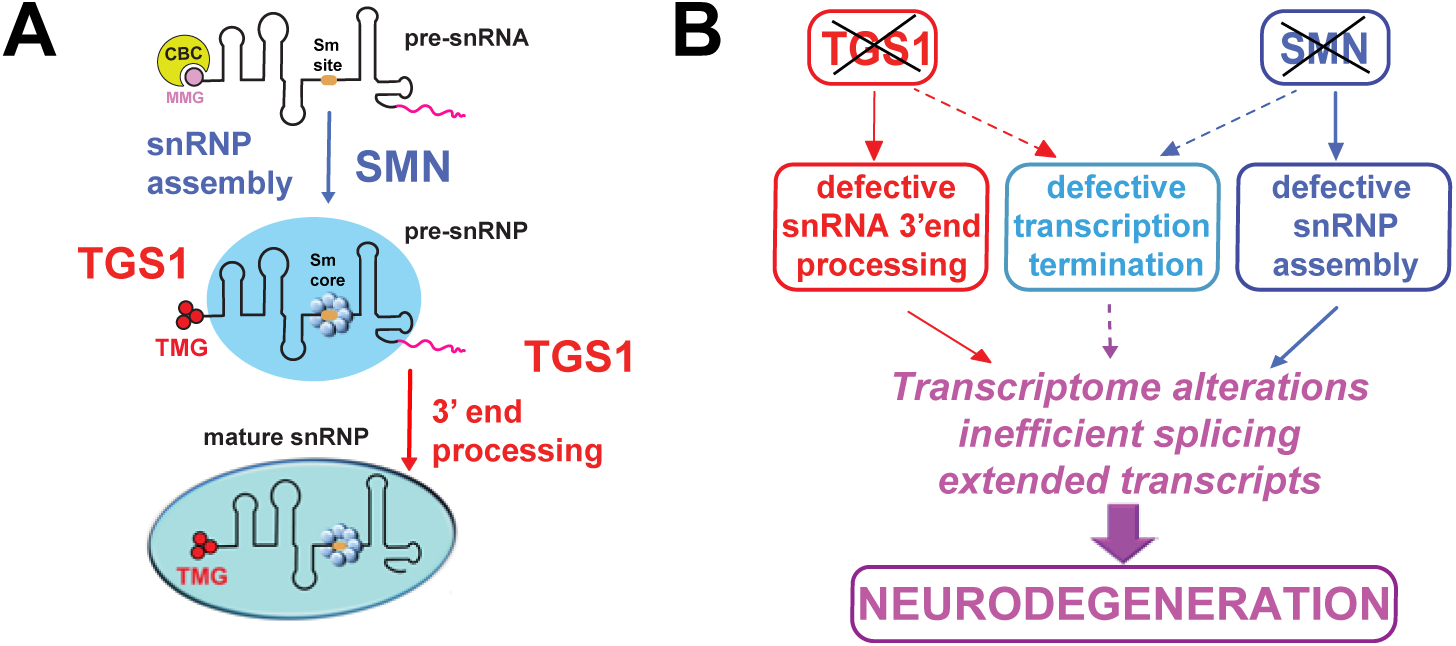
SMN and TGS1 protect against neurodegeneration through both common and specific routes. **(A)** Roles of SMN and TGS1 in snRNA biogenesis. **(B)** A model for the roles of TGS1 and SMN in prevention of neurodegeneration. The effects of TGS1 or SMN deficiency on snRNA biogenesis and general transcription suggest that these proteins play interconnected roles in prevention of neurodegeneration.

Recent remarkable advances in the field have led to the approval of disease modifying therapies for SMA patients, which include SMN replacement via gene therapy as well as increased production of full-length SMN by splicing modulation with antisense oligonucleotides or small molecules (116–119). However, it is evident that these treatments do not represent a cure for the disease and expansion of therapeutic options for SMA patient is needed. This will likely require the development of SMN-independent strategies targeting disease-relevant downstream pathways (116,117,120–125).

Candidate targets have been emerging from studies in model organisms (71,78,90,91,126). Here, we have shown that TGS1 overexpression can compensate, at least in part, for SMN loss in animal models of neurodegeneration. Although further research is needed to analyze the functional relationships between SMN and TGS1, our results indicate that modulation of TGS1 activity may represent a new potential strategy to integrate SMA therapies.

## Data Availability

Custom python scripts for 3’-RACE: https://cmroake.people.stanford.edu/links-python-scripts

RNAend-Seq data: SRA BioProject ID PRJNA628085 Transcriptome analyses: https://github.com/roozbehdn/TGS1_SMN

Illumina and Nanopore RNA sequencing data available at:

https://sales.bio.unipd.it/bulk/e5606676c2072ade71c85745c6ea25c8d8c00aca235ca9cf 6aa5f04079883c06/

GEO accession number:

## Acknowledgements

We thank P. Bazzicalupo for critical reading of the manuscript, G. Zampi and F. Sola for technical support, S. Schneider for support in zebrafish procedures, P. De Piante for data analysis. For strains and plasmids we thank A. Fire, K. Shen, O. Hobert, C. Bargmann and the Caenorhabditis Genetics Center (CGC), funded by NIH Office of Research Infrastructure Programs (P40 OD010440). We thank Reinhard.Luehrmann for sharing the R1131 TMG antibody.

## SUPPLEMENTARY FIGURE LEGENDS

**Figure S1.**
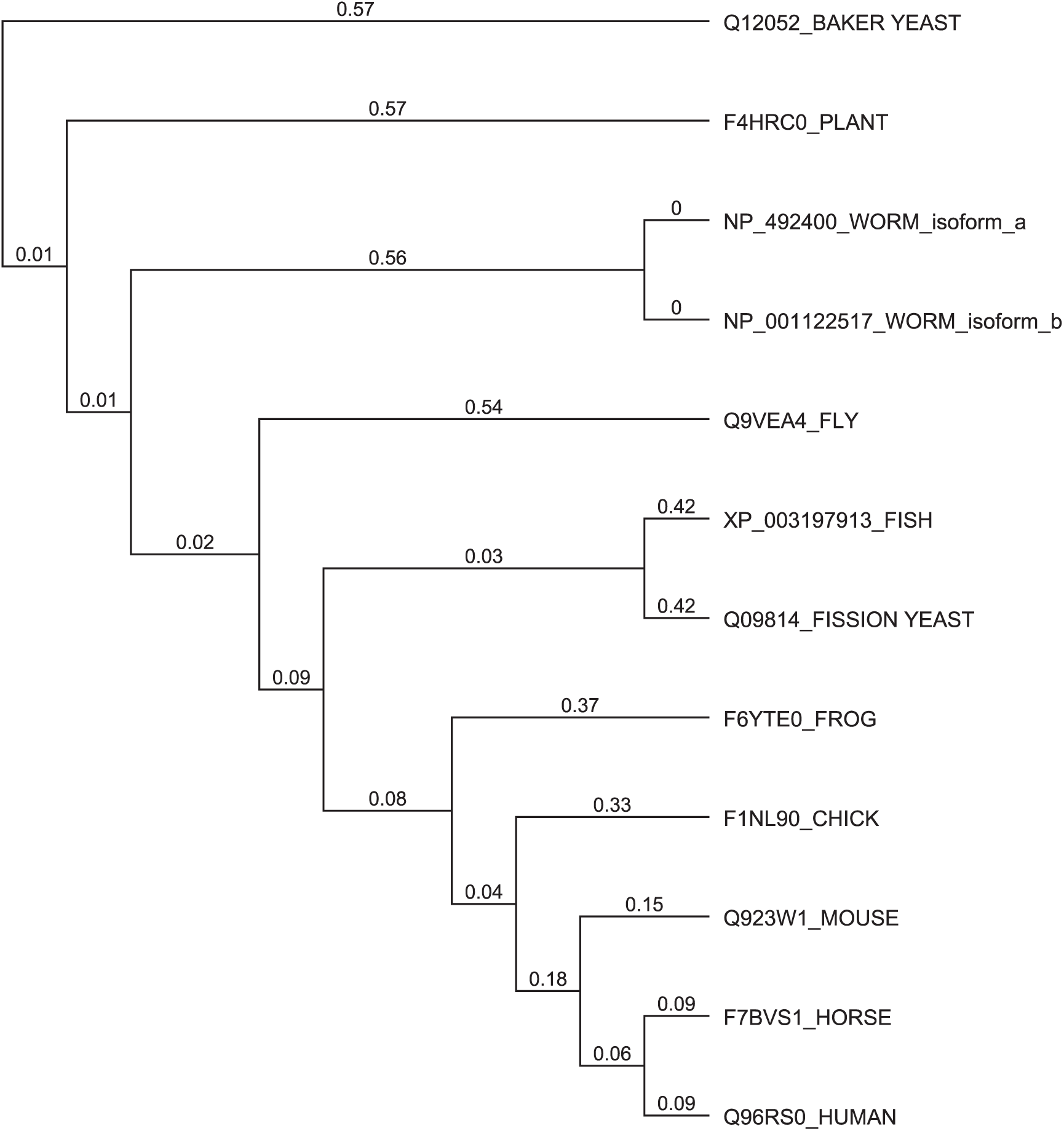
TGS1 orthologs. Evolutionary tree showing the phylogenetic relationship among TGS1 proteins from diverse organisms.

**Figure S2.**
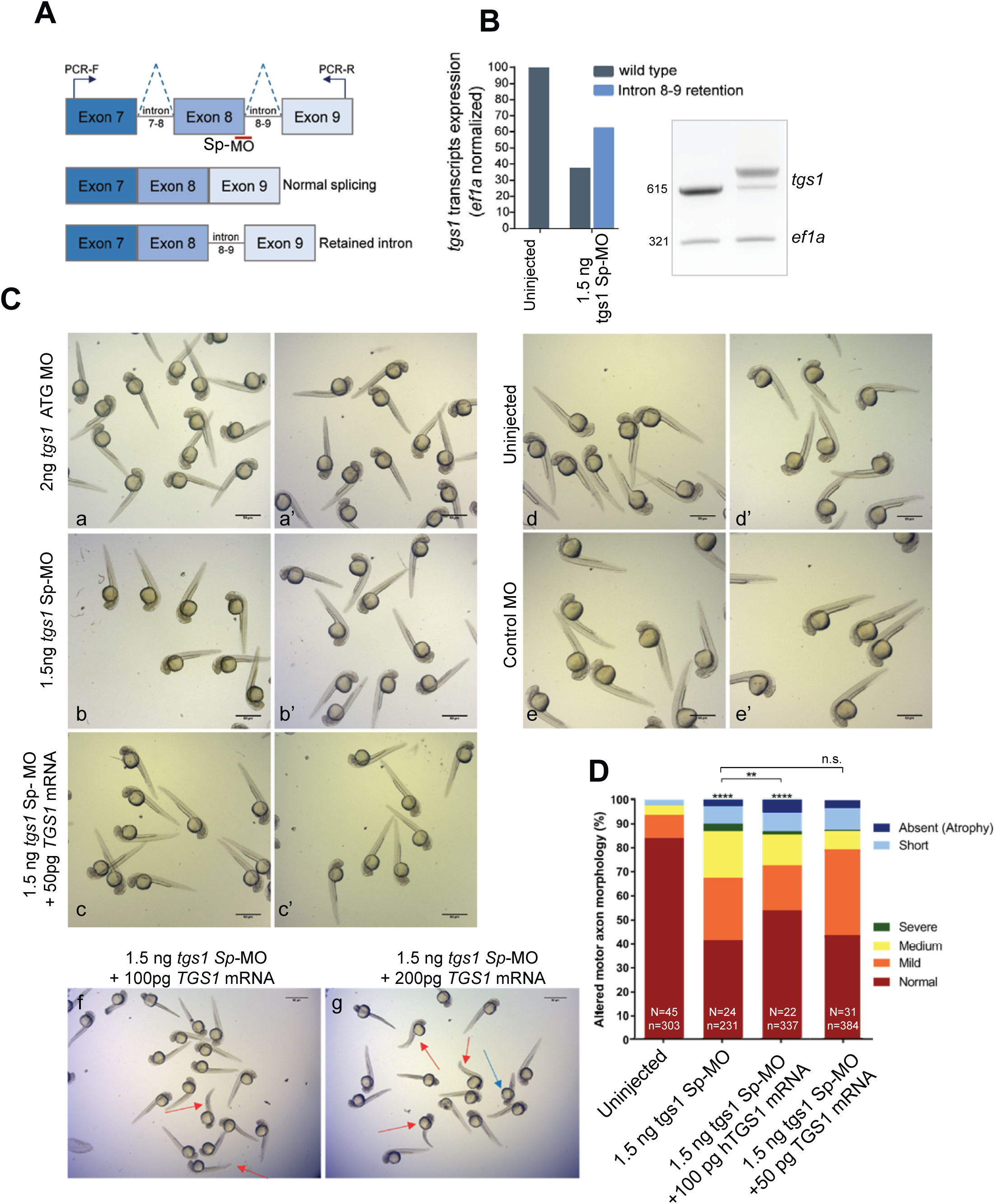
Injection of a *tgs1* Sp-MO induces intron 8 retention and causes CaP-MN defects. **(A)** Diagram showing the exon/introns structure of the *tgs1* gene. The location of the *tgs1* Sp-MO at the intron-exon boundary is indicated with a red line. The dotted line indicates normal splicing of introns 7 and 8. Arrows: primers used for RT-PCR to verify that the *tgs1* Sp-MO injection induces accumulation of mis-spliced *tgs1* transcripts retaining intron 8. **(B)** Representative agarose gel showing *tgs1* mRNA products as measured by semi- quantitative RT-PCR in uninjected (control) and *tgs1* Sp-MO injected embryos. As compared with control fish, injection of the *tgs1* Sp-MO reduced the amount of correctly spliced transcripts (represented by the wildtype amplicon of 615 bp) and induced an intron 8-retained splice transcript of 750 bp. This transcript contains a stop codon upstream the sequence encoding the *tgs1* catalytic domain and is therefore predicted to produce a non- functional protein. Densitometric analysis of *tgs1* transcripts normalized against the *ef1a* housekeeping gene shows that *tgs1 Sp-*MO reduces WT *tgs1* transcripts by approximately 65%. **(C)** Morphology of zebrafish larvae after *tgs1* MOs and *TGS1* mRNA injection. Bright field images of fish larvae injected with *tgs1* MOs (alone or in combination with an RNA encoding FLAG-tagged human TGS1 (*hTGS1*) and control groups (control MO and uninjected). Gross morphological defects are not observed upon injection of 2 ng *tgs1* ATG MO (panels a, a’), 1.5 ng *tgs1* Sp-MO (alone or in combination with 50 pg *TGS1* mRNA; panels b, b’, c, c’), and 1.5 ng control-non targeting – MO (panels d, d’) or in uninjected animals (e, e’). In contrast, injection of 1.5 ng *tgs1* Sp-MO in combination with 100 pg or 200 pg h*TGS1* RNA resulted in cardiac edema, tail bending and body axis malformations (arrows in panels f, g). Scale bar: 50 µm. **(D)** Based on overall appearance, CaP-MNs were classified as described in Fig. 4B. Zebrafish larvae were injected with 1.5 ng of *tgs1 Sp-*MO alone or in combination with 100 pg or 50 pg h*TGS1* RNA. Results are presented in percentages from 3 independent experiments (n = axons analysed; N= animals tested. ****, p<0.0001 Chi-square test); n.s. not significant. The capacity of 100 pg h*TGS1* RNA to rescue the neuronal defects induced by injection of *tgs1 Sp-*MO cannot be unequivocally assessed, as larvae injected with this dosage of h*TGS1* RNA display developmental abnormalities (shown in image f of panel C).

**Figure S3.**
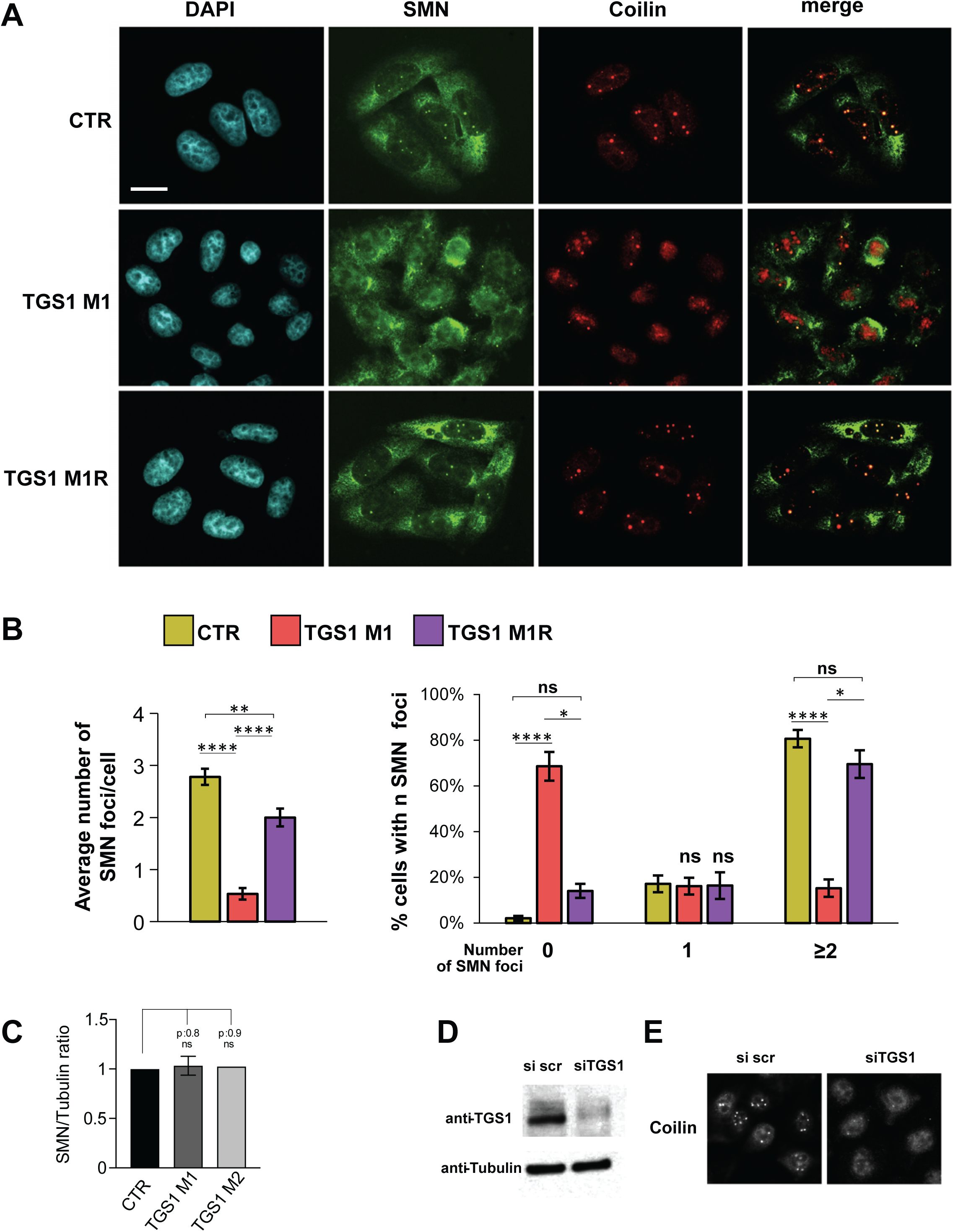
*TGS1* knockdown disrupts Cajal body organization in HeLa and UMUC3 cells. **(A)** CB organization in *TGS1*-proficient (CTR), *TGS1 M1* and *TGS1 M1R* HeLa cells. Cells were stained with anti-SMN and anti-Coilin antibodies and with DAPI (DNA). SMN accumulates in the CBs and colocalizes with the CB marker Coilin in 99% of control cells. Depletion of TGS1 leads to a reduction in the coilin-rich CBs, loss of SMN nuclear localization, and ectopic SMN granules in the cytoplasm. These defects are ameliorated by stable expression of a *FLAG*-*TGS1* transgene (M1R). Scale bar, 20 µm. **(B)** Quantification of the number of SMN/Coilin positive foci in the nuclei of HeLa cells with the indicated genotypes. At least 200 cells counted per sample. Error bars, S.E.M. *, **, **** p < 0.05, 0.01 and 0.0001, respectively; ns, not significant, two-way ANOVA). **(C)** Quantification of the SMN protein in western blots performed on protein extracts from TGS1 mutant HeLa cell clones, probed with the anti-SMN and anti-Tubulin antibodies. Data from three replicates are relative to tubulin. p values: one-way ANOVA. See also Figure 4A and Supplementary Figure S6B. **(D)** Western blotting showing TGS1 abundance in UMUC3 cells treated with *TGS1* siRNA or non-targeting dsRNA (si scr) for 6 days. Tubulin is a loading control. **(E)** UMUC3 cells treated with *TGS1* siRNA or non-targeting dsRNA (si scr) stained with anti-Coilin antibodies and DAPI. Note that *TGS1* knockdown disrupts CB organization.

**Figure S4.**
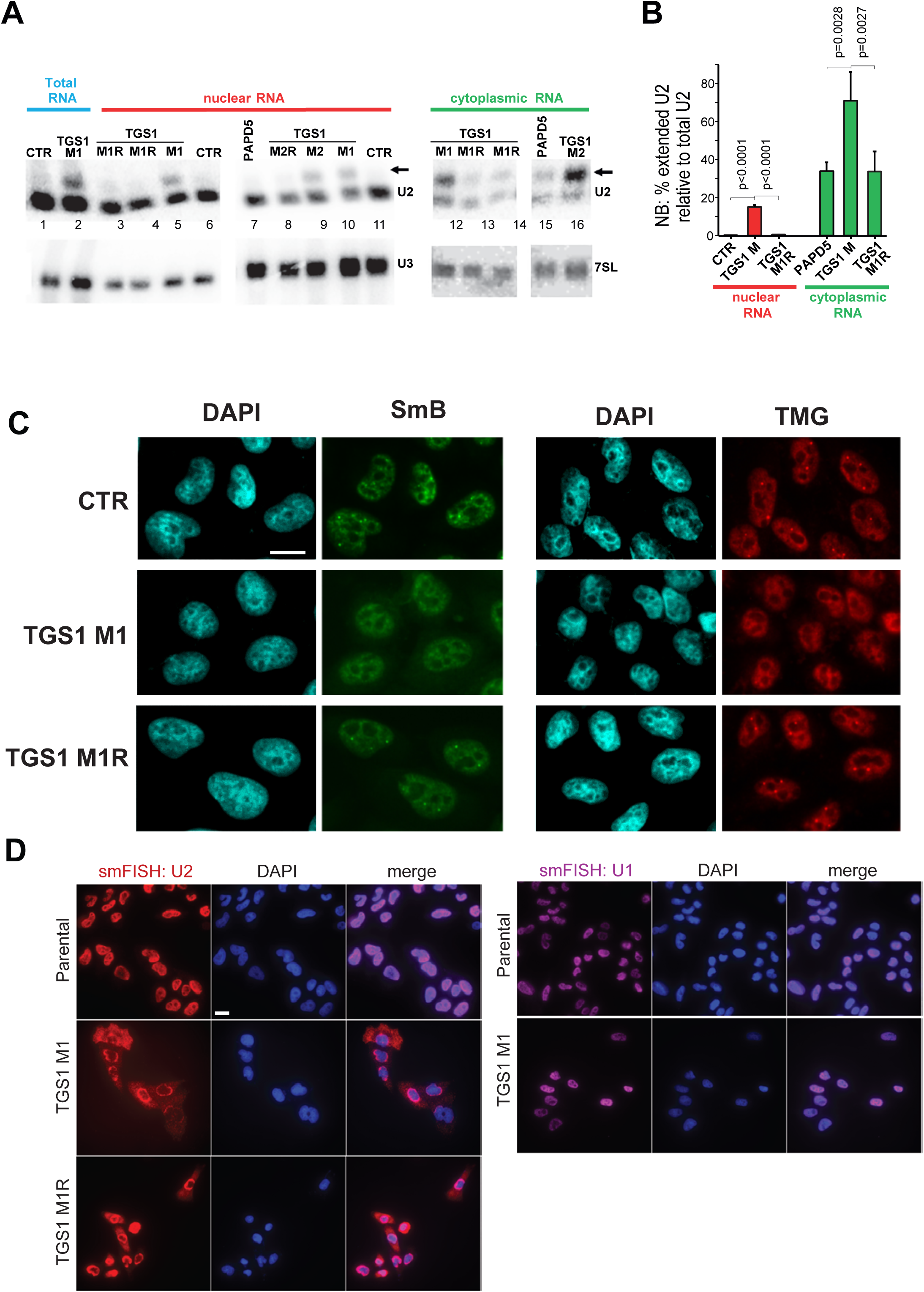
TGS1 affects 3′ end processing of U2 snRNA and alters subcellular snRNPs localization. **(A)** Northern blots showing U2 snRNA precursors (arrow) in nuclear and cytoplasmic fractions from HeLa cells of the indicated genotypes; the U3 and 7SL RNAs are loading controls for the nuclear and cytoplasmic fractions, respectively. CTR, parental cell line; M1 and M2, *TGS1* mutant cells. *M1R* and *M2R*, *TGS1 M1* and *M2* rescued cells. *PAPD5*, HeLa clone carrying mutations in *PAPD5*, used as negative control. **(B)** Quantification of the blots in (A) by densitometric analysis. Bars: percentage of extended U2 molecules relative to total U2 (± SEM). p values: One-way ANOVA with post- hoc Tukey’s test. **(C)** Localization of snRNPs in *TGS1 M1, TGS1 M1R* and *CTR* cells. snRNP localization was detected by immunostaining with anti-SmB (green) and anti-TMG (red) antibodies in the cells described in (A). Note the loss of the punctate nuclear foci and the presence of a halo in the cytoplasm of *TGS1 M1* mutant cells. DNA was stained with DAPI (blue). **(D)** Single-molecule Fluorescent *in situ* Hybridization (smFISH, see methods) showing the cellular localization of U2 and U1 snRNA in *TGS1 M1*, *TGS1 M1R* and control (CTR) cells. Note the cytoplasmic accumulation of U2 snRNA in *TGS1 M1* cells, compared to TGS1- proficient cells, in which U2 is mainly nuclear. In contrast, U1 snRNA localization is unaffected in *TGS1 M1* cells. Scale bar, 20 µm.

**Figure S5.**
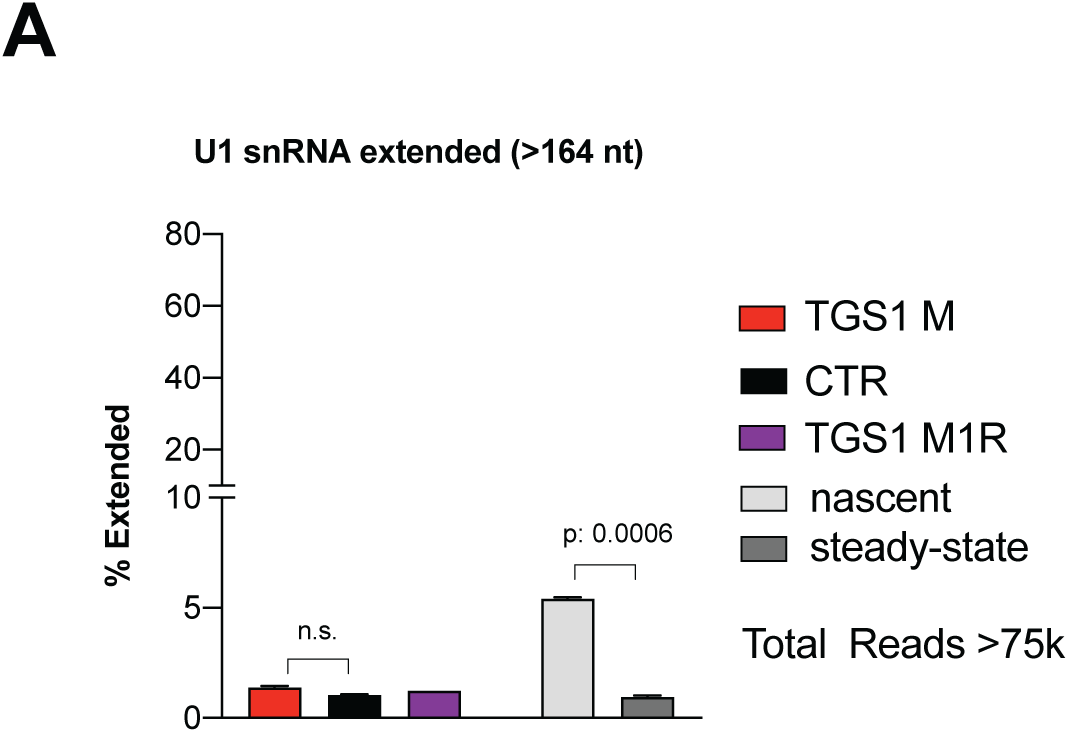
Characterization of the 3′ ends of U1 snRNA in TGS1-deficient cells. **(A)** 3’ RACE and sequencing on total RNA showing that the U1snRNA from *TGS1 M1*, *TGS1 M1R* and control (CTR) HeLa cells exhibit similar 3′ ends. Nascent and steady-state RNA fractions were purified from control HeLa cells (see Methods). Data are mean values of two independent clones + SEM. p values: two-sided Student’s t test. Average reads per sample > 161k ± 40k.

**Figure S6.**
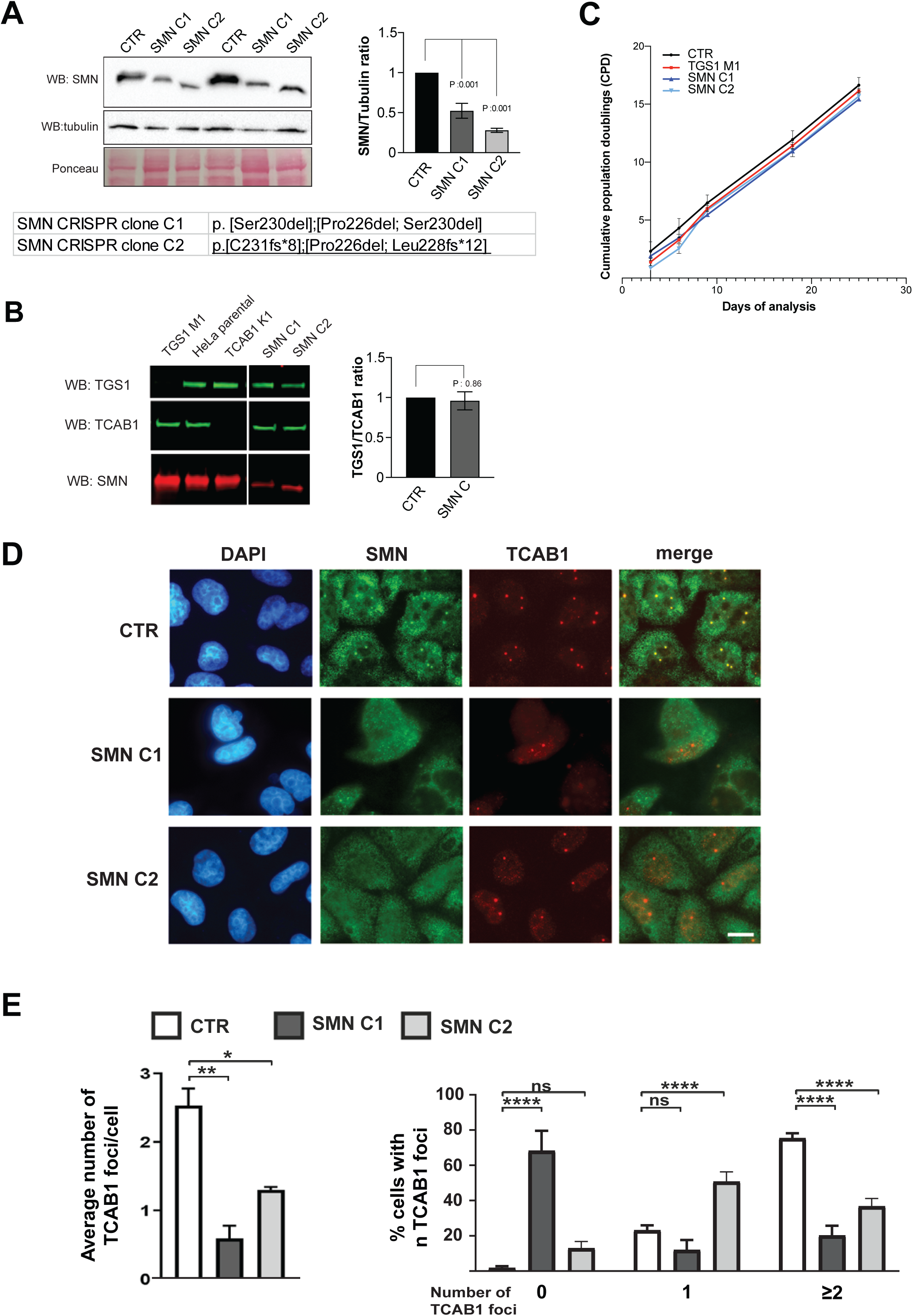
Characterization of CRISPR-induced *SMN* mutant Hela cells. **(A)** Left panel, SMN expression in two independent CRISPR-derived *SMN* mutant HeLa cell clones (C1, C2) revealed by Western blotting (WB). CTR, parental cell line; tubulin is a loading control. The mutations in the *SMN* gene found in *SMN C1* and *SMN C2* mutant cells are indicated below the WB. *SMN C1* mutant cells carry a heterozygous deletion of Pro226 (the last residue in the polyproline helix) and a homozygous deletion of Ser230 (a missense mutation in this residue is found in patients affected by type II SMA (1); *SMN C2* mutant cells carry a heterozygous deletion of Pro226 and heterozygous frameshift mutations: Leu228ter, a mutation frequently found in Chinese SMA families (2) and C231ter. Note that the proteins encoded by the *SMN C2* alleles lack the C-terminal YG box, which controls SMN dimerization and higher order oligomerization (3); they also have a lower MW compared to the wild type protein. Right panel, quantification of SMN protein level in the *SMN C1* and *SMN C2* mutant clones relative to tubulin and normalized to to *SMN*-proficient HeLa cells; p values: one-way ANOVA. **(B)** Left panel, Western blots probed with anti-TGS1 and anti-SMN antibodies, showing that the abundance of TGS1 is unaffected in *SMN C1* and *SMN C2* mutant lines, compared to the parental HeLa cells and *TCAB1 K1* cells (used as a loading control) that lack the CB component TCAB1. **(C)** Cumulative population doubling (CPD) analysis showing similar growth rates in control (CTR), *TGS1 M1*, *SMN C1* and *SMN C2* HeLa cell lines. Population doublings were measured at each passage until day 25. Data are expressed as means ± SD (n = 2). **(D)** CB organization in *SMN*-proficient (CTR), *SMN C1* and *SMN C2* mutant HeLa cells. Cells were stained with anti-SMN and anti-TCAB1 antibodies and counterstained with DAPI (DNA). In control cells, the SMN and TCAB1 proteins are enriched in CBs (SMN and TCAB1 colocalize in 99% of cells). In both *SMN* mutant clones, CB organization is altered, as indicated by loss of the nuclear accumulation of SMN mutant proteins and a reduced numbers of TCAB1-enriched nuclear foci (quantified in (E). **(E)** Quantification of the TCAB1-positive foci in the nuclei of cells of the indicated genotypes. At least 200 cells scored per sample. Error bars, S.E.M. Statistical significance was determined by two-way ANOVA. Scale bar, 5 μm.

**Figure S7.**
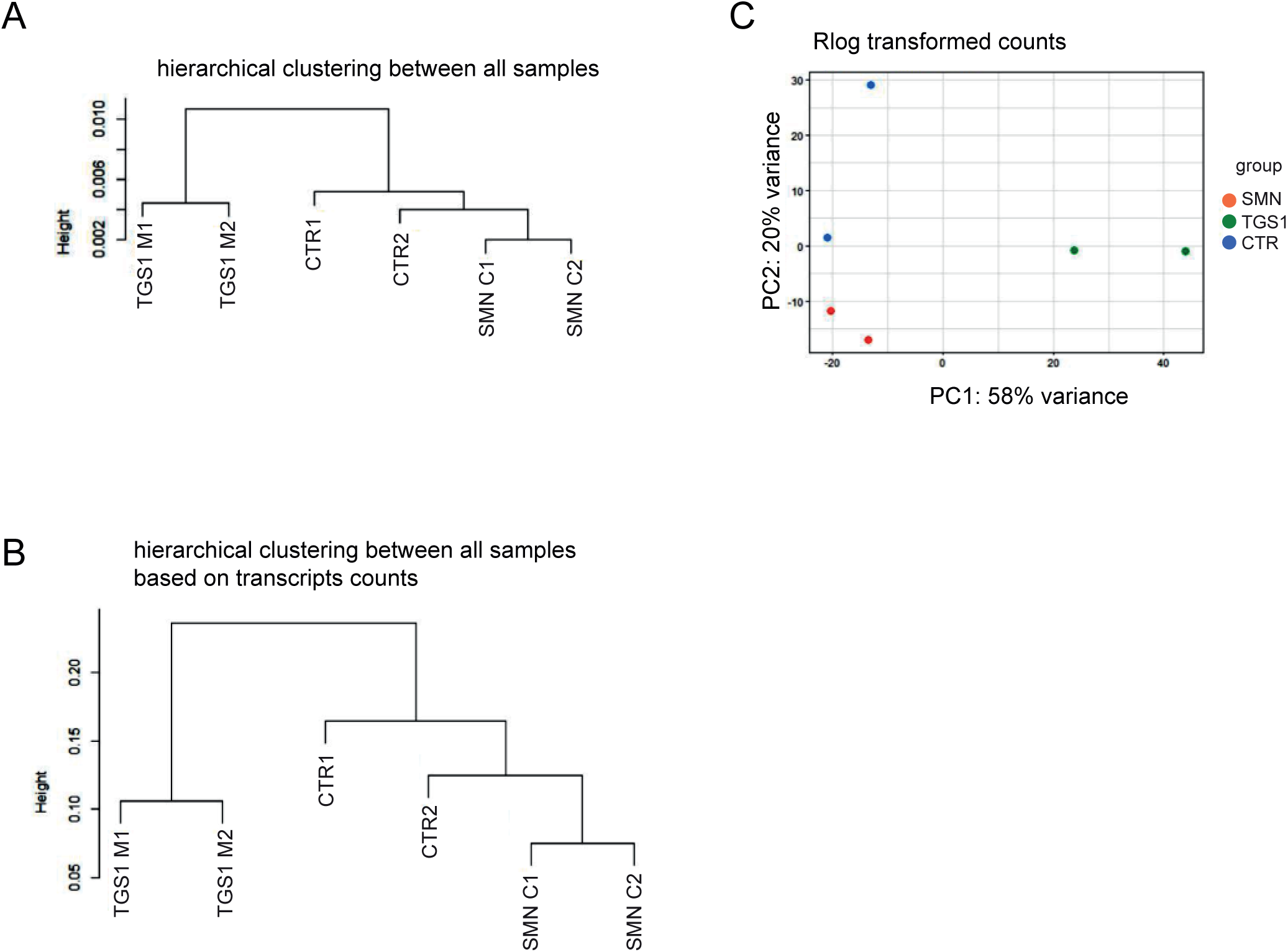
Hierarchical clustering and PCA analysis of RNA samples from CTR, *TGS1* and *SMN* mutant cells. (A, B) Hierarchical clustering and principal component analysis based on gene expression (A, C) and transcript expression (B) show similarity among samples within the same group.

**Figure S8.**
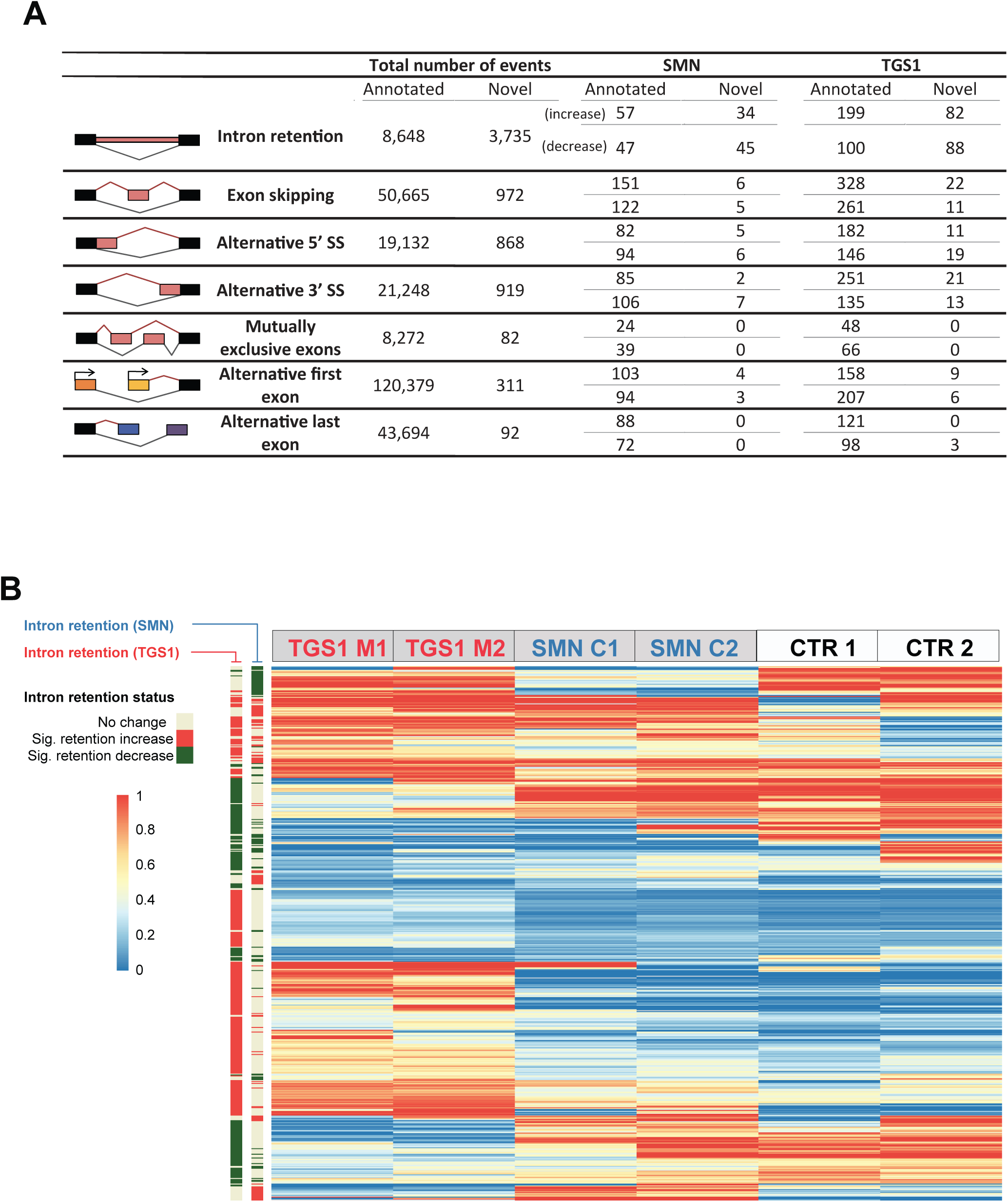
Patterns of alternative splicing in TGS1 and SMN mutant cells (Illumina sequencing) (A) Number of significantly changed alternative splicing (AS) events across 7 major AS types. Annotated AS events are extracted from the GENCODE annotation and novel AS events are obtained from the assembled transcriptome by Scallop (4). The heading “Total number of events” indicates the total number of alternative splicing events in each category and “SMN” and “TGS1” refer to the number of differentially spliced events in each mutant category, where the first and second rows in each category show the number of events with increased and decreased alternative isoforms, respectively. (B) Heatmap of the Intron Retention (IR) events in *SMN* and *TGS1* mutant cells as annotated by two sidebars to the left (increased IR marked in Magenta and decreased in Green). Heatmap shows the PSI value of a given IR event across all six RNA-seq datasets.

**Figure S9.**
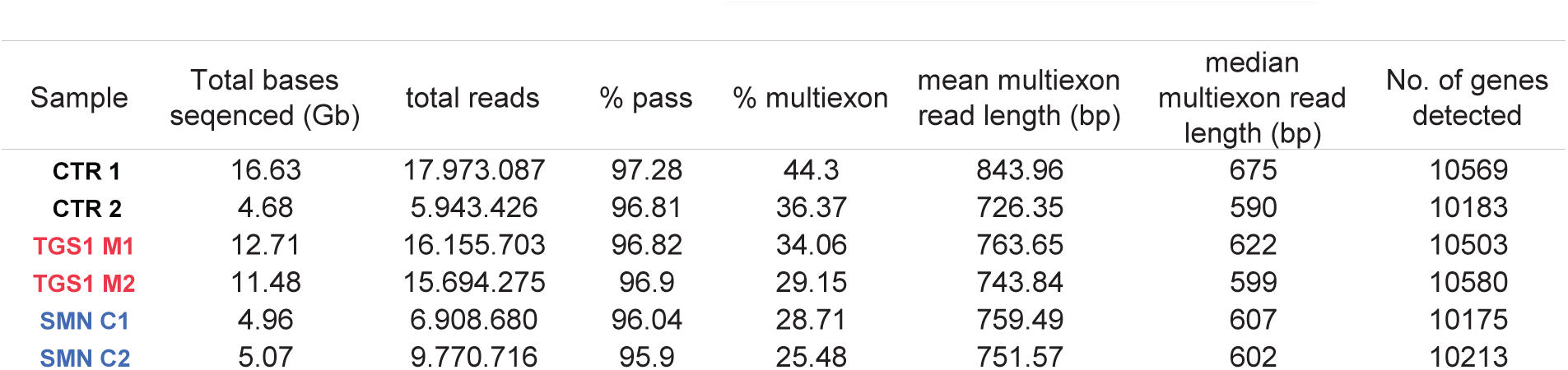
Reads produced by Oxford Nanopore sequencing runs. Statistics for each run: estimated bases (measured in gigabases); % pass: indicates barcoded reads with an average quality score ≥ 7 (found using Guppy). % multiexon: reads that map across at least one splicing junction. Number of detected genes is the number of genes with at least one read. Data acquisition was performed with MinKNOW Core 4.1.2 for samples *CTR1, CTR2, TGS1 M1, TGS1 M2* and MinKNOW Core 4.2.5 for *SMN C1* and *SMN C2*.

**Figure S10.**
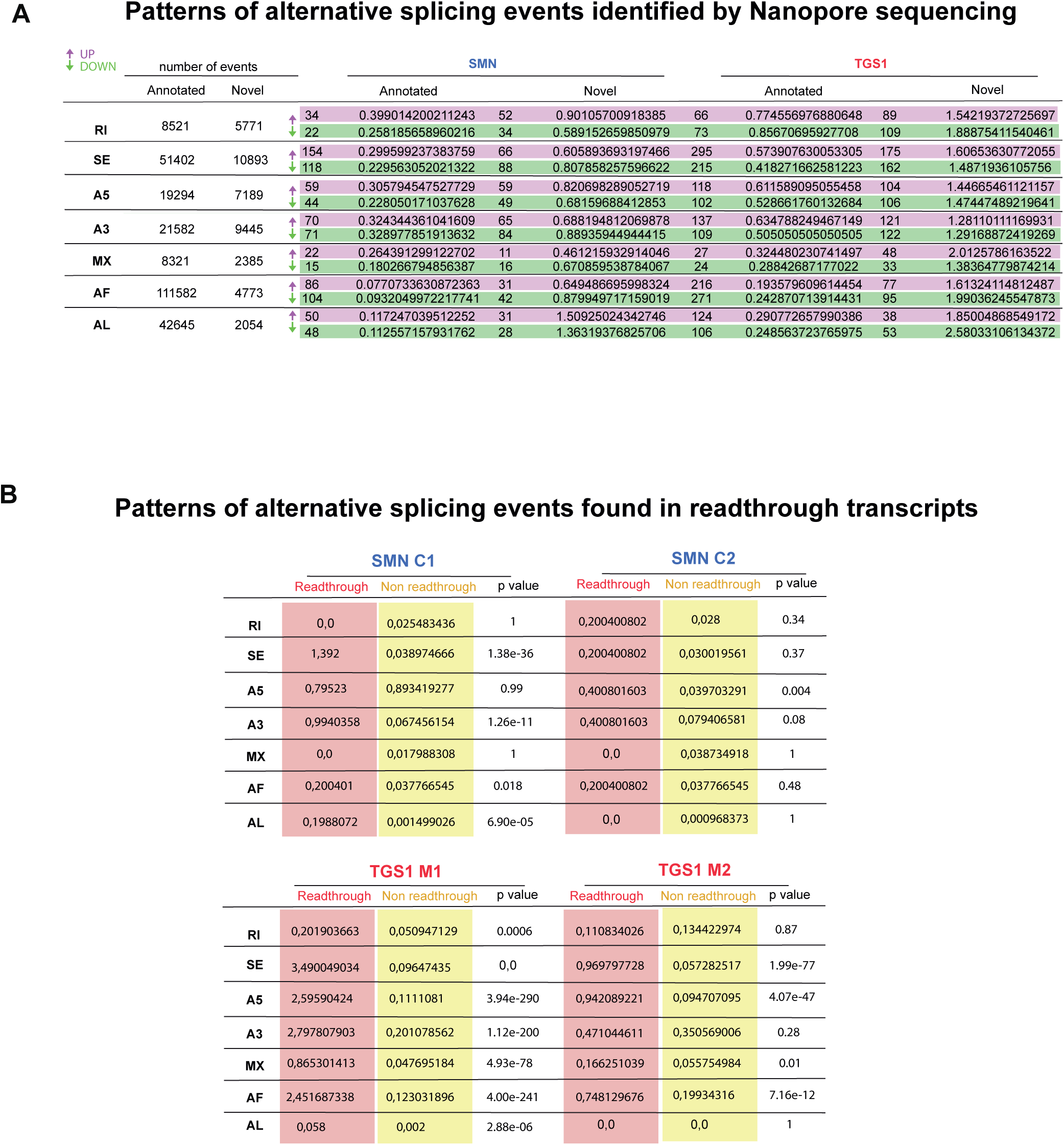
Patterns of alternative splicing in *TGS1* and *SMN* mutant cells (Nanopore sequencing) (A) Numbers of annotated and novel Alternative Splicing (AS) events across seven major AS types (RI: intron retention; SE: exon skipping; A5: alternative 5′ splice site; A3:alternative 3′ splice site; MX: mutually exclusive exon; AF: alternative first exon; AL: alternative last exon). Annotated AS events are those described in the GENCODE annotation, while novel AS events are those present only in the transcriptome assembled by FLAIR (5). “SMN” and “TGS1” columns show the number of annotated and novel differential splicing events for each AS category, with a preferential inclusion of the events in the mutant (dPSI >0.1; up arrow) or in the WT (dPSI< -0.1; down arrow). PSI, dPSI and significance of the differential splicing were calculated using SUPPA2. See also Figures 8B-C and Supplementary table S6. (B) Patterns of alternative splicing events, across the 7 major AS types, identified in readthrough and non-readthrough transcripts produced in mutant samples from genes with “Preferential Readthrough in Mutant”. The percentages of readthrough and non-readthrough reads with AS were calculated as described in Figure 8G, considering each AS type independently. The significance of the difference in the proportions of reads with aberrant splicing was calculated using chi-square test. See also Supplementary table S10.

**Figure S11.**
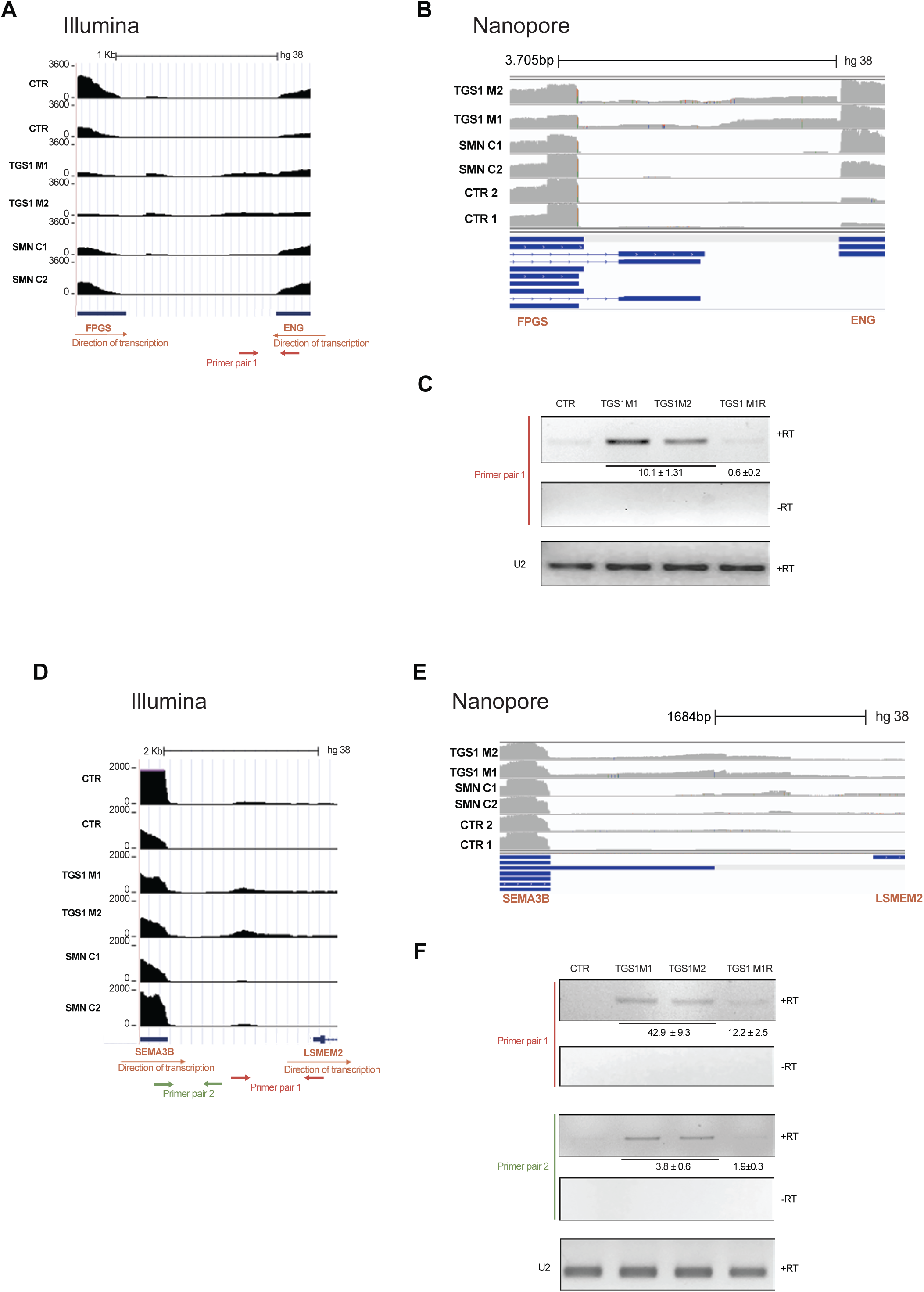
Validation of transcriptional events spanning intergenic regions identified by Illumina and Nanopore sequencing of *TGS1* mutant cells. (A) The densities of Illumina reads in the *FPGS*-*ENG* region are reported in the diagrams from the UCSC Genome browser. The positions of the primer pairs designed to perform RT-PCR analyses (red arrows) are indicated below the diagram (A). (B) Diagram illustrating the densities of Nanopore long reads in the FPGS-ENG region from Integrative genomic viewer (IGV). Blue lines represent annotated transcripts. (C) RT-PCR analysis performed on the indicated samples with the primer pair shown in (A); densitometric quantitation of the amplification products is relative to the U2 amplification product. (D) The densities of Illumina reads in the SEMA3B-LSMEM2 region are reported in the diagrams from the UCSC Genome browser. The positions of the primer pairs designed to perform RT-PCR analyses (red and green arrows) are indicated below the diagram (E) Diagram illustrating the densities of Nanopore long reads in the SEMA3B-LSMEM2 region from Integrative genomic viewer (IGV). Blue lines represent annotated transcripts. (F) RT-qPCR analysis performed on the indicated samples with the primer pair shown in (D); densitometric quantitation of the amplification products is relative to the U2 amplification product. – RT indicates absence of reverse transcriptase; see text for definition of HeLa cell genotypes.

**Figure S12.**
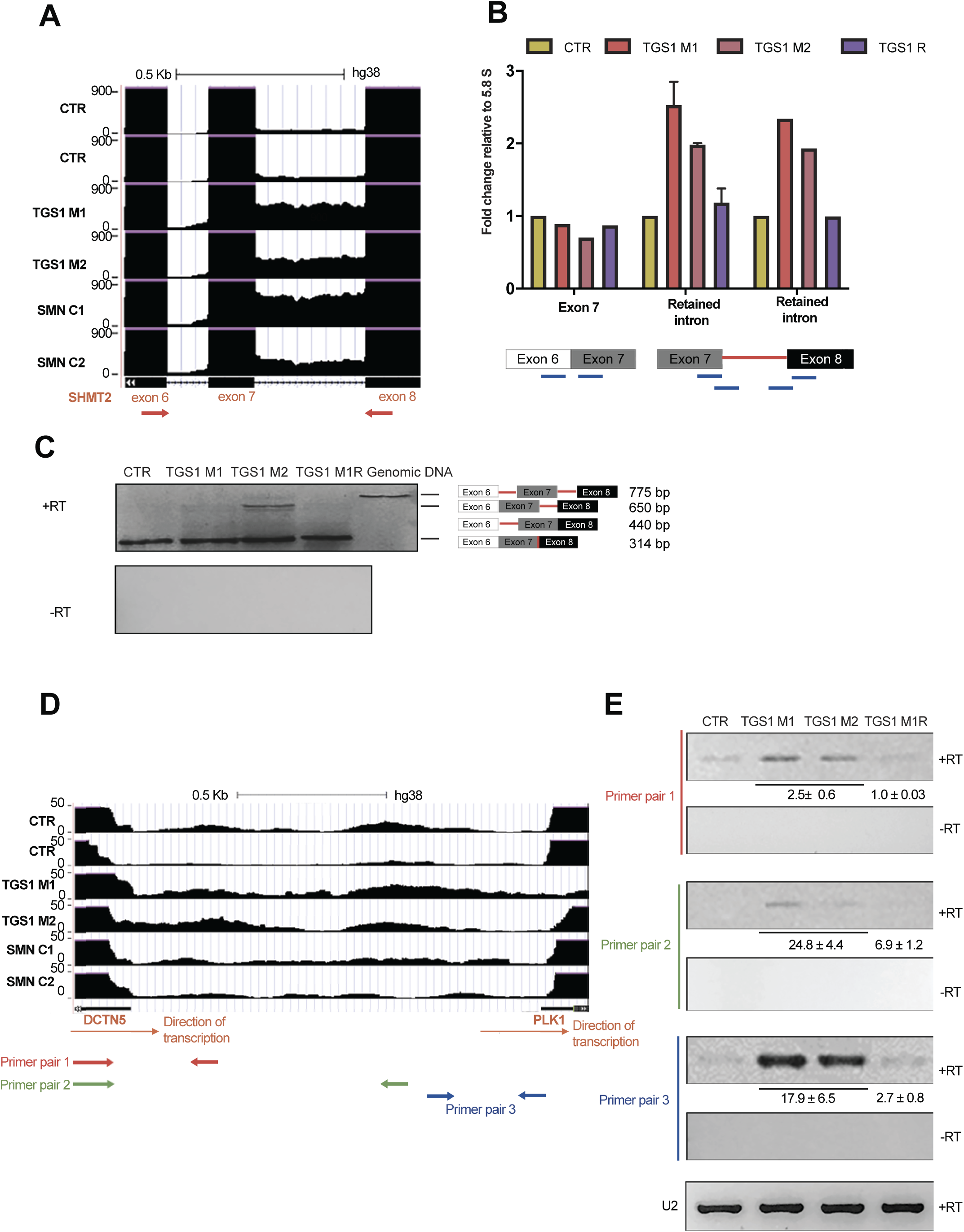
Validation of aberrant transcript structures from in silico analyses in *TGS1* mutant cells (identified in Illumina sequencing) (A-C): Detection of intron 7 retention in the SHMT2 gene. (A) Diagram from USCS Genome browser showing the reads mapping to intron 7 in *TGS1 M1*, *TGS1 M2* and control (CTR) cells. Transcripts containing this intron are also enriched in SMN mutant cells. See text for definition of HeLa cell genotypes. (B) RTq-PCR comparative quantification of the retention level of intron 7 assayed using the indicated primer pairs (blue lines shown below the histograms) and the abundance of the adjacent exon 7. RT-qPCR was performed on cDNA samples from *TGS1 M1*, *TGS1 M2*, *TGS1 M1R* and Control (CTR) HeLa cell lines. Bars represent means between three replicates, are relative to the parental cell line (CTR) and are normalized to 5.8S rRNA. (C) RT-qPCR analysis performed on cDNA samples from *TGS1 M1*, *TGS1 M2*, *TGS1 M1R* and Control (CTR) HeLa cell lines to detect the retention of intron 7. The positions of the primer pairs used for RT-PCR spanning exons 6 to 8 are reported below panel A (red arrows). In negative control samples (-RT), the reverse transcriptase was omitted from the reaction to rule out genomic DNA contamination. Genomic DNA amplified by PCR is a control. The numbers below the panels show the densitometric quantitation of the amplification products relative to the U2 amplification product (+/- SEM, three replicates) (D-E). Validation of transcriptional events spanning the intergenic region between the DCTN5 and PLK1 genes. (D) Diagram from USCS Genome browser shows reads mapping to the region of interest in the sequenced samples. The blue, red and green arrows indicate the positions of the primer pairs designed to perform RT-qPCR analysis shown in panel E. (E) RT-qPCR analysis of the genomic region indicated in panel D. U2 is a positive control.

## SUPPLEMENTARY TABLES available at: https://figshare.com/ s/13572f55424106336cac

**Table S1.** Differentially expressed transcripts in TGS1 and SMN HeLa mutant cells compared to control cells obtained by running the Kallisto+Sleuth pipeline on Illumina reads. List of the transcripts differentially expressed in TGS1 (sig_transcripts_WT_vs_TGS1) or SMN (sig_transcripts_WT_vs_SMN) mutant cells compared to control cells (WT), identified and quantified using Kallisto + Sleuth. Annotated and unannotated transcript isoforms are denoted by ENST- and gene- prefixes, respectively. Transcripts with Beta values less than -2 or greater than 2 and with q values < 0.05 were considered to be significantly differentially expressed (positive values: downregulated; negative values: upregulated)

**Table S2.** List of differential alternative splicing events in TGS1 and SMN HeLa mutant cells compared to control cells obtained by running SUPPA2 on Illumina reads

**Table S3.** List of the transcripts differentially expressed in TGS1 or SMN mutant cells compared to control cells obtained by running the Salmon + DESeq2 on illumine reads. List of the transcripts differentially expressed in TGS1 (TGS1 vs CTR filter) or SMN (SMN vs CTR filter) mutant cells compared to control cells (WT), identified and quantified using the combination of the Salmon, Scallop and DESeq2 softwares. Annotated and unannotated transcript isoforms are denoted by ENST- and gene- prefixes, respectively. Transcripts with log_2_ fold change values less than -2 or greater than 2 and with adjusted p values < 0.05 were considered to be significantly differentially expressed. Sheet *locations*: genomic coordinates for unannotated transcripts are reported.

**Table S4.** Summary of the total number of DE transcripts identified using two independent quantification pipelines

**Table S5.** Classification of novel transcripts

The unannotated transcripts were classified according to the length of their 5′ or 3′ UTR and for the presence of retained introns. For those transcripts carrying unspliced introns (labeled as TRUE), the genomic location of the retained sequence is indicated

**Table S6.** dPSI and dTPM all AS events.

**Table S7.** TGS1 and SMN AS events intersection.

**Table S8.** Common AS events Illumina vs Nanopore.

**Table S9.** Transcript readthrough in TGS1 and SMN.

**Table S10.** Readthrough transcripts and aberrant splicing.

**Table S11.** List of the oligonucleotides used in this work.

## REFERENCES

1. Nussbacher, J.K., Tabet, R., Yeo, G.W. and Lagier-Tourenne, C. (2019) Disruption of RNA Metabolism in Neurological Diseases and Emerging Therapeutic Interventions. Neuron, 102, 294–320.

2. Wirth, B., Karakaya, M., Kye, M.J. and Mendoza-Ferreira, N. (2020) Twenty-Five Years of Spinal Muscular Atrophy Research: From Phenotype to Genotype to Therapy, and What Comes Next. Annu Rev Genomics Hum Genet, 21, 231–261.

3. Lardelli, R.M., Schaffer, A.E., Eggens, V.R., Zaki, M.S., Grainger, S., Sathe, S., Van Nostrand, E.L., Schlachetzki, Z., Rosti, B., Akizu, N. et al. (2017) Biallelic mutations in the 3’ exonuclease TOE1 cause pontocerebellar hypoplasia and uncover a role in snRNA processing. Nat Genet, 49, 457–464.

4. de Vegvar, H.E., Lund, E. and Dahlberg, J.E. (1986) 3’ end formation of U1 snRNA precursors is coupled to transcription from snRNA promoters. Cell, 47, 259–266.

5. Hernandez, N. and Weiner, A.M. (1986) Formation of the 3’ end of U1 snRNA requires compatible snRNA promoter elements. Cell, 47, 249–258.

6. Egloff, S., Zaborowska, J., Laitem, C., Kiss, T. and Murphy, S. (2012) Ser7 phosphorylation of the CTD recruits the RPAP2 Ser5 phosphatase to snRNA genes. Mol Cell, 45, 111–122.

7. O’Reilly, D., Kuznetsova, O.V., Laitem, C., Zaborowska, J., Dienstbier, M. and Murphy, S. (2014) Human snRNA genes use polyadenylation factors to promote efficient transcription termination. Nucleic Acids Res, 42, 264–275.

8. Baillat, D., Hakimi, M.A., Naar, A.M., Shilatifard, A., Cooch, N. and Shiekhattar, R. (2005) Integrator, a multiprotein mediator of small nuclear RNA processing, associates with the C-terminal repeat of RNA polymerase II. Cell, 123, 265–276.

9. Ohno, M., Segref, A., Bachi, A., Wilm, M. and Mattaj, I.W. (2000) PHAX, a mediator of U snRNA nuclear export whose activity is regulated by phosphorylation. Cell, 101, 187–198.

10. Kitao, S., Segref, A., Kast, J., Wilm, M., Mattaj, I.W. and Ohno, M. (2008) A compartmentalized phosphorylation/dephosphorylation system that regulates U snRNA export from the nucleus. Mol Cell Biol, 28, 487–497.

11. Yong, J., Kasim, M., Bachorik, J.L., Wan, L. and Dreyfuss, G. (2010) Gemin5 delivers snRNA precursors to the SMN complex for snRNP biogenesis. Mol Cell, 38, 551–562.

12. Pellizzoni, L., Yong, J. and Dreyfuss, G. (2002) Essential role for the SMN complex in the specificity of snRNP assembly. Science, 298, 1775–1779.

13. Mouaikel, J., Verheggen, C., Bertrand, E., Tazi, J. and Bordonne, R. (2002) Hypermethylation of the cap structure of both yeast snRNAs and snoRNAs requires a conserved methyltransferase that is localized to the nucleolus. Mol Cell, 9, 891–901.

14. Mouaikel, J., Narayanan, U., Verheggen, C., Matera, A.G., Bertrand, E., Tazi, J. and Bordonne, R. (2003) Interaction between the small-nuclear-RNA cap hypermethylase and the spinal muscular atrophy protein, survival of motor neuron. EMBO Rep, 4, 616–622.

15. Narayanan, U., Achsel, T., Luhrmann, R. and Matera, A.G. (2004) Coupled in vitro import of U snRNPs and SMN, the spinal muscular atrophy protein. Mol Cell, 16, 223–234.

16. Strasser, A., Dickmanns, A., Luhrmann, R. and Ficner, R. (2005) Structural basis for m3G-cap-mediated nuclear import of spliceosomal UsnRNPs by snurportin1. EMBO J, 24, 2235–2243.

17. Fischer, U., Heinrich, J., van Zee, K., Fanning, E. and Luhrmann, R. (1994) Nuclear transport of U1 snRNP in somatic cells: differences in signal requirement compared with Xenopus laevis oocytes. J Cell Biol, 125, 971–980.

18. Marshallsay, C. and Luhrmann, R. (1994) In vitro nuclear import of snRNPs: cytosolic factors mediate m3G-cap dependence of U1 and U2 snRNP transport. EMBO J, 13, 222–231.

19. Huber, J., Cronshagen, U., Kadokura, M., Marshallsay, C., Wada, T., Sekine, M. and Luhrmann, R. (1998) Snurportin1, an m3G-cap-specific nuclear import receptor with a novel domain structure. Embo J, 17, 4114–4126.

20. Stanek, D. (2017) Cajal bodies and snRNPs - friends with benefits. RNA biology, 14, 671–679.

21. Wieben, E.D., Nenninger, J.M. and Pederson, T. (1985) Ribonucleoprotein organization of eukaryotic RNA. XXXII. U2 small nuclear RNA precursors and their accurate 3’ processing in vitro as ribonucleoprotein particles. Journal of molecular biology, 183, 69–78.

22. Brenner, S. (1974) The genetics of Caenorhabditis elegans. Genetics, 77, 71–94.

23. Gallotta, I., Mazzarella, N., Donato, A., Esposito, A., Chaplin, J.C., Castro, S., Zampi, G., Battaglia, G.S., Hilliard, M.A., Bazzicalupo, P. et al. (2016) Neuron-specific knock-down of SMN1 causes neuron degeneration and death through an apoptotic mechanism. Hum Mol Genet, 25, 2564–2577.

24. Hobert, O. (2002) PCR fusion-based approach to create reporter Gene constructs for expression analysis in transgenic C. elegans. BioTechniques, 32, 728–730.

25. Esposito, G., Di Schiavi, E., Bergamasco, C. and Bazzicalupo, P. (2007) Efficient and cell specific knock-down of gene function in targeted C. elegans neurons. Gene, 395, 170–176.

26. Eastman, C., Horvitz, H.R. and Jin, Y. (1999) Coordinated transcriptional regulation of the unc-25 glutamic acid decarboxylase and the unc-47 GABA vesicular transporter by the Caenorhabditis elegans UNC-30 homeodomain protein. The Journal of neuroscience : the official journal of the Society for Neuroscience, 19, 6225–6234.

27. Kamath, R.S., Fraser, A.G., Dong, Y., Poulin, G., Durbin, R., Gotta, M., Kanapin, A., Le Bot, N., Moreno, S., Sohrmann, M. et al. (2003) Systematic functional analysis of the Caenorhabditis elegans genome using RNAi. Nature, 421, 231–237.

28. Mello, C.C., Kramer, J.M., Stinchcomb, D. and Ambros, V. (1991) Efficient gene transfer in C.elegans: extrachromosomal maintenance and integration of transforming sequences. The EMBO journal, 10, 3959–3970.

29. Veronico, P., Gray, L.J., Jones, J.T., Bazzicalupo, P., Arbucci, S., Cortese, M.R., Di Vito, M. and De Giorgi, C. (2001) Nematode chitin synthases: Gene structure, expression and function in Caenorhabditis elegans and the plant parasitic nematode Meloidogyne artiellia. Molecular Genetics and Genomics, 266, 28–34.

30. Kimble, J.E. and White, J.G. (1981) On the control of germ cell development in Caenorhabditis elegans. Dev Biol, 81, 208–219.

31. Stinchcomb, D.T., Shaw, J.E., Carr, S.H. and Hirsh, D. (1985) Extrachromosomal DNA transformation of Caenorhabditis elegans. Mol Cell Biol, 5, 3484–3496.

32. McIntire, S.L., Reimer, R.J., Schuske, K., Edwards, R.H. and Jorgensen, E.M. (1997) Identification and characterization of the vesicular GABA transporter. Nature, 389, 870–876.

33. Riessland, M., Kaczmarek, A., Schneider, S., Swoboda, K.J., Lohr, H., Bradler, C., Grysko, V., Dimitriadi, M., Hosseinibarkooie, S., Torres-Benito, L. et al. (2017) Neurocalcin Delta Suppression Protects against Spinal Muscular Atrophy in Humans and across Species by Restoring Impaired Endocytosis. American journal of human genetics, 100, 297–315.

34. Mendoza-Ferreira, N., Coutelier, M., Janzen, E., Hosseinibarkooie, S., Lohr, H., Schneider, S., Milbradt, J., Karakaya, M., Riessland, M., Pichlo, C. et al. (2018) Biallelic CHP1 mutation causes human autosomal recessive ataxia by impairing NHE1 function. *Neurology*. Genetics, 4, e209.

35. Lan, C.C., Tang, R., Un San Leong, I. and Love, >D.R. (2009) Quantitative real-time RT-PCR (qRT-PCR) of zebrafish transcripts: optimization of RNA extraction, quality control considerations, and data analysis. Cold Spring Harbor protocols, 2009, pdb prot5314.

36. Oprea, G.E., Krober, S., McWhorter, M.L., Rossoll, W., Muller, S., Krawczak, M., Bassell, G.J., Beattie, C.E. and Wirth, B. (2008) Plastin 3 is a protective modifier of autosomal recessive spinal muscular atrophy. Science, 320, 524–527.

37. Chang, H.C., Dimlich, D.N., Yokokura, T., Mukherjee, A., Kankel, M.W., Sen, A., Sridhar, V., Fulga, T.A., Hart, A.C., Van Vactor, D. et al. (2008) Modeling spinal muscular atrophy in Drosophila. PLoS One, 3, e3209.

38. Palumbo, V., Pellacani, C., Heesom, K.J., Rogala, K.B., Deane, C.M., Mottier-Pavie, V., Gatti, M., Bonaccorsi, S. and Wakefield, J.G. (2015) Misato Controls Mitotic Microtubule Generation by Stabilizing the TCP-1 Tubulin Chaperone Complex [corrected]. Curr Biol, 25, 1777–1783.

39. Bischof, J., Maeda, R.K., Hediger, M., Karch, F. and Basler, K. (2007) An optimized transgenesis system for Drosophila using germ-line-specific phiC31 integrases. Proc Natl Acad Sci U S A, 104, 3312–3317.

40. Maccallini, P., Bavasso, F., Scatolini, L., Bucciarelli, E., Noviello, G., Lisi, V., Palumbo, V., D’Angeli, S., Cacchione, S., Cenci, G. et al. (2020) Intimate functional interactions between TGS1 and the Smn complex revealed by an analysis of the Drosophila eye development. PLoS Genet, 16, e1008815.

41. Buszczak, M., Paterno, S., Lighthouse, D., Bachman, J., Planck, J., Owen, S., Skora, A.D., Nystul, T.G., Ohlstein, B., Allen, A. et al. (2007) The carnegie protein trap library: a versatile tool for Drosophila developmental studies. Genetics, 175, 1505–1531.

42. Raffa, G.D., Siriaco, G., Cugusi, S., Ciapponi, L., Cenci, G., Wojcik, E. and Gatti, M. (2009) The Drosophila modigliani (moi) gene encodes a HOAP-interacting protein required for telomere protection. Proceedings of the National Academy of Sciences of the United States of America, 106, 2271–2276.

43. Lattao, R., Bonaccorsi, S., Guan, X., Wasserman, S.A. and Gatti, M. (2011) Tubby-tagged balancers for the Drosophila X and second chromosomes. Fly, 5, 369–370.

44. Ran, F.A., Hsu, P.D., Wright, J., Agarwala, V., Scott, D.A. and Zhang, F. (2013) Genome engineering using the CRISPR-Cas9 system. Nat Protoc, 8, 2281–2308.

45. Chen, L., Roake, C.M., Galati, A., Bavasso, F., Micheli, E., Saggio, I., Schoeftner, S., Cacchione, S., Gatti, M., Artandi, S.E. et al. (2020) Loss of Human TGS1 Hypermethylase Promotes Increased Telomerase RNA and Telomere Elongation. Cell Rep, 30, 1358–1372 e1355.

46. Xu, L. and Blackburn, E.H. (2007) Human cancer cells harbor T-stumps, a distinct class of extremely short telomeres. Mol Cell, 28, 315–327.

47. Roake, C.M., Chen, L., Chakravarthy, A.L., Ferrell, J.E., Raffa, G.D. and Artandi, S.E. (2019) Disruption of Telomerase RNA Maturation Kinetics Precipitates Disease. Molecular Cell, 74, 688–700.e683.

48. Chen, L., Roake, C.M., Freund, A., Batista, P.J., Tian, S., Yin, Y.A., Gajera, C.R., Lin, S., Lee, B., Pech, M.F. et al. (2018) An Activity Switch in Human Telomerase Based on RNA Conformation and Shaped by TCAB1. Cell, 174, 218–230 e213.

49. Venteicher, A.S., Abreu, E.B., Meng, Z., McCann, K.E., Terns, R.M., Veenstra, T.D., Terns, M.P. and Artandi, S.E. (2009) A human telomerase holoenzyme protein required for Cajal body localization and telomere synthesis. Science, 323, 644–648.

50. Luhrmann, R., Appel, B., Bringmann, P., Rinke, J., Reuter, R., Rothe, S. and Bald, R. (1982) Isolation and characterization of rabbit anti-m3 2,2,7G antibodies. Nucleic Acids Res, 10, 7103–7113.

51. Livak, K.J. and Schmittgen, T.D. (2001) Analysis of relative gene expression data using real-time quantitative PCR and the 2(-Delta Delta C(T)) Method. Methods, 25, 402–408.

52. Chen, L., Ooi, S.K., Conaway, R.C. and Conaway, J.W. (2014) Generation and purification of human INO80 chromatin remodeling complexes and subcomplexes. J Vis Exp, e51720.

53. Dobin, A., Davis, C.A., Schlesinger, F., Drenkow, J., Zaleski, C., Jha, S., Batut, P., Chaisson, M. and Gingeras, T.R. (2013) STAR: ultrafast universal RNA-seq aligner. Bioinformatics, 29, 15–21.

54. Shao, M. and Kingsford, C. (2017) Accurate assembly of transcripts through phase-preserving graph decomposition. Nature biotechnology, 35, 1167–1169.

55. Pimentel, H., Bray, N.L., Puente, S., Melsted, P. and Pachter, L. (2017) Differential analysis of RNA-seq incorporating quantification uncertainty. Nature methods, 14, 687–690.

56. Trincado, J.L., Entizne, J.C., Hysenaj, G., Singh, B., Skalic, M., Elliott, D.J. and Eyras, E. (2018) SUPPA2: fast, accurate, and uncertainty-aware differential splicing analysis across multiple conditions. Genome biology, 19, 40.

57. Patro, R., Duggal, G., Love, M.I., Irizarry, R.A. and Kingsford, C. (2017) Salmon provides fast and bias-aware quantification of transcript expression. Nature methods, 14, 417–419.

58. Li, H. (2018) Minimap2: pairwise alignment for nucleotide sequences. Bioinformatics, 34, 3094–3100.

59. Tang, A.D., Soulette, C.M., van Baren, M.J., Hart, K., Hrabeta-Robinson, E., Wu, C.J. and Brooks, A.N. (2020) Full-length transcript characterization of SF3B1 mutation in chronic lymphocytic leukemia reveals downregulation of retained introns. Nature communications, 11, 1438.

60. Frankish, A., Diekhans, M., Ferreira, A.M., Johnson, R., Jungreis, I., Loveland, J., Mudge, J.M., Sisu, C., Wright, J., Armstrong, J. et al. (2019) GENCODE reference annotation for the human and mouse genomes. Nucleic Acids Res, 47, D766–D773.

61. Quinlan, A.R. and Hall, I.M. (2010) BEDTools: a flexible suite of utilities for comparing genomic features. Bioinformatics, 26, 841–842.

62. Shaye, D.D. and Greenwald, I. (2011) OrthoList: a compendium of C. elegans genes with human orthologs. PLoS One, 6, e20085.

63. Spencer, W.C., Zeller, G., Watson, J.D., Henz, S.R., Watkins, K.L., McWhirter, R.D., Petersen, S., Sreedharan, V.T., Widmer, C., Jo, J. et al. (2011) A spatial and temporal map of C. elegans gene expression. Genome Research, 21, 325–341.

64. Kaletsky, R., Lakhina, V., Arey, R., Williams, A., Landis, J., Ashraf, J. and Murphy, C.T. (2016) The C. elegans adult neuronal IIS/FOXO transcriptome reveals adult phenotype regulators. Nature, 529, 92–96.

65. Rugarli, E.I., Di Schiavi, E., Hilliard, M.A., Arbucci, S., Ghezzi, C., Facciolli, A., Coppola, G., Ballabio, A. and Bazzicalupo, P. (2002) The Kallmann syndrome gene homolog in C. elegans is involved in epidermal morphogenesis and neurite branching. *Development (Cambridge*, England), 129, 1283–1294.

66. McIntire, S.L., Jorgensen, E., Kaplan, J. and Horvitz, H.R. (1993) The GABAergic nervous system of Caenorhabditis elegans. Nature, 364, 337–341.

67. Komonyi, O., Schauer, T., Papai, G., Deak, P. and Boros, I.M. (2009) A product of the bicistronic Drosophila melanogaster gene CG31241, which also encodes a trimethylguanosine synthase, plays a role in telomere protection. J Cell Sci, 122, 769–774.

68. Borg, R.M., Fenech Salerno, B., Vassallo, N., Bordonne, R. and Cauchi, R.J. (2016) Disruption of snRNP biogenesis factors Tgs1 and pICln induces phenotypes that mirror aspects of SMN-Gemins complex perturbation in Drosophila, providing new insights into spinal muscular atrophy. Neurobiology of disease, 94, 245–258.

69. Di Giorgio, M.L., Esposito, A., Maccallini, P., Micheli, E., Bavasso, F., Gallotta, I., Vernì, F., Feiguin, F., Cacchione, S., McCabe, B.D., et al. (2017) WDR79/TCAB1 plays a conserved role in the control of locomotion and ameliorates phenotypic defects in SMA models. Neurobiology of disease, 105, 42–50.

70. McWhorter, M.L., Monani, U.R., Burghes, A.H. and Beattie, C.E. (2003) Knockdown of the survival motor neuron (Smn) protein in zebrafish causes defects in motor axon outgrowth and pathfinding. J Cell Biol, 162, 919–931.

71. Hosseinibarkooie, S., Peters, M., Torres-Benito, L., Rastetter, R.H., Hupperich, K., Hoffmann, A., Mendoza-Ferreira, N., Kaczmarek, A., Janzen, E., Milbradt, J. et al. (2016) The Power of Human Protective Modifiers: PLS3 and CORO1C Unravel Impaired Endocytosis in Spinal Muscular Atrophy and Rescue SMA Phenotype. American journal of human genetics, 99, 647–665.

72. Tisdale, S. and Pellizzoni, L. (2015) Disease mechanisms and therapeutic approaches in spinal muscular atrophy. J Neurosci, 35, 8691–8700.

73. Tycowski, K.T., Aab, A. and Steitz, J.A. (2004) Guide RNAs with 5’ caps and novel box C/D snoRNA-like domains for modification of snRNAs in metazoa. Curr Biol, 14, 1985–1995.

74. Jia, D., Cai, L., He, H., Skogerbo, G., Li, T., Aftab, M.N. and Chen, R. (2007) Systematic identification of non-coding RNA 2,2,7-trimethylguanosine cap structures in Caenorhabditis elegans. BMC molecular biology, 8, 86.

75. Becker, D., Hirsch, A.G., Bender, L., Lingner, T., Salinas, G. and Krebber, H. (2019) Nuclear Pre-snRNA Export Is an Essential Quality Assurance Mechanism for Functional Spliceosomes. Cell Rep, 27, 3199–3214 e3193.

76. Goldfarb, K.C. and Cech, T.R. (2013) 3’ terminal diversity of MRP RNA and other human noncoding RNAs revealed by deep sequencing. BMC molecular biology, 14, 23.

77. Cho, H.D., Tomita, K., Suzuki, T. and Weiner, A.M. (2002) U2 small nuclear RNA is a substrate for the CCA-adding enzyme (tRNA nucleotidyltransferase). J Biol Chem, 277, 3447–3455.

78. Rizzo, F., Nizzardo, M., Vashisht, S., Molteni, E., Melzi, V., Taiana, M., Salani, S., Santonicola, P., Di Schiavi, E., Bucchia, M. et al. (2019) Key role of SMN/SYNCRIP and RNA-Motif 7 in spinal muscular atrophy: RNA-Seq and motif analysis of human motor neurons. Brain : a journal of neurology, 142, 276–294.

79. Doktor, T.K., Hua, Y., Andersen, H.S., Broner, S., Liu, Y.H., Wieckowska, A., Dembic, M., Bruun, G.H., Krainer, A.R. and Andresen, B.S. (2017) RNA-sequencing of a mouse-model of spinal muscular atrophy reveals tissue-wide changes in splicing of U12-dependent introns. Nucleic Acids Res, 45, 395–416.

80. Jangi, M., Fleet, C., Cullen, P., Gupta, S.V., Mekhoubad, S., Chiao, E., Allaire, N., Bennett, C.F., Rigo, F., Krainer, A.R. et al. (2017) SMN deficiency in severe models of spinal muscular atrophy causes widespread intron retention and DNA damage. Proc Natl Acad Sci U S A, 114, E2347–E2356.

81. Huo, Q., Kayikci, M., Odermatt, P., Meyer, K., Michels, O., Saxena, S., Ule, J. and Schumperli, D. (2014) Splicing changes in SMA mouse motoneurons and SMN-depleted neuroblastoma cells: evidence for involvement of splicing regulatory proteins. RNA biology, 11, 1430–1446.

82. Zhang, Z., Lotti, F., Dittmar, K., Younis, I., Wan, L., Kasim, M. and Dreyfuss, G. (2008) SMN deficiency causes tissue-specific perturbations in the repertoire of snRNAs and widespread defects in splicing. Cell, 133, 585–600.

83. Lotti, F., Imlach, W.L., Saieva, L., Beck, E.S., Hao le, T., Li, D.K., Jiao, W., Mentis, G.Z., Beattie, C.E., McCabe, B.D. et al. (2012) An SMN-dependent U12 splicing event essential for motor circuit function. Cell, 151, 440–454.

84. Bray, N.L., Pimentel, H., Melsted, P. and Pachter, L. (2016) Near-optimal probabilistic RNA-seq quantification. Nature biotechnology, 34, 525–527.

85. Peabody, N.C., Diao, F., Luan, H., Wang, H., Dewey, E.M., Honegger, H.W. and White, B.H. (2008) Bursicon functions within the Drosophila CNS to modulate wing expansion behavior, hormone secretion, and cell death. J Neurosci, 28, 14379–14391.

86. Jia, Y., Mu, J.C. and Ackerman, S.L. (2012) Mutation of a U2 snRNA gene causes global disruption of alternative splicing and neurodegeneration. Cell, 148, 296–308.

87. Tisdale, S., Lotti, F., Saieva, L., Van Meerbeke, J.P., Crawford, T.O., Sumner, C.J., Mentis, G.Z. and Pellizzoni, L. (2013) SMN is essential for the biogenesis of U7 small nuclear ribonucleoprotein and 3’-end formation of histone mRNAs. Cell Rep, 5, 1187–1195.

88. Ruggiu, M., McGovern, V.L., Lotti, F., Saieva, L., Li, D.K., Kariya, S., Monani, U.R., Burghes, A.H. and Pellizzoni, L. (2012) A role for SMN exon 7 splicing in the selective vulnerability of motor neurons in spinal muscular atrophy. Mol Cell Biol, 32, 126–138.

89. Simon, C.M., Dai, Y., Van Alstyne, M., Koutsioumpa, C., Pagiazitis, J.G., Chalif, J.I., Wang, X., Rabinowitz, J.E., Henderson, C.E., Pellizzoni, L. et al. (2017) Converging Mechanisms of p53 Activation Drive Motor Neuron Degeneration in Spinal Muscular Atrophy. Cell Rep, 21, 3767–3780.

90. Simon, C.M., Van Alstyne, M., Lotti, F., Bianchetti, E., Tisdale, S., Watterson, D.M., Mentis, G.Z. and Pellizzoni, L. (2019) Stasimon Contributes to the Loss of Sensory Synapses and Motor Neuron Death in a Mouse Model of Spinal Muscular Atrophy. Cell Rep, 29, 3885–3901 e3885.

91. Van Alstyne, M., Simon, C.M., Sardi, S.P., Shihabuddin, L.S., Mentis, G.Z. and Pellizzoni, L. (2018) Dysregulation of Mdm2 and Mdm4 alternative splicing underlies motor neuron death in spinal muscular atrophy. Genes Dev, 32, 1045–1059.

92. Son, A., Park, J.E. and Kim, V.N. (2018) PARN and TOE1 Constitute a 3’ End Maturation Module for Nuclear Non-coding RNAs. Cell Rep, 23, 888–898.

93. Lardelli, R.M. and Lykke-Andersen, J. (2020) Competition between maturation and degradation drives human snRNA 3’ end quality control. Genes Dev, 34, 989–1001.

94. Cheng, L., Zhang, Y., Zhang, Y., Chen, T., Xu, Y.Z. and Rong, Y.S. (2020) Loss of the RNA trimethylguanosine cap is compatible with nuclear accumulation of spliceosomal snRNAs but not pre-mRNA splicing or snRNA processing during animal development. PLoS Genet, 16, e1009098.

95. Madore, S.J., Wieben, E.D., Kunkel, G.R. and Pederson, T. (1984) Precursors of U4 small nuclear RNA. J Cell Biol, 99, 1140–1144.

96. Madore, S.J., Wieben, E.D. and Pederson, T. (1984) Intracellular site of U1 small nuclear RNA processing and ribonucleoprotein assembly. J Cell Biol, 98, 188–192.

97. Ezzeddine, N., Chen, J., Waltenspiel, B., Burch, B., Albrecht, T., Zhuo, M., Warren, W.D., Marzluff, W.F. and Wagner, E.J. (2011) A subset of Drosophila integrator proteins is essential for efficient U7 snRNA and spliceosomal snRNA 3’-end formation. Mol Cell Biol, 31, 328–341.

98. Shukla, S. and Parker, R. (2014) Quality control of assembly-defective U1 snRNAs by decapping and 5’-to-3’ exonucleolytic digestion. Proc Natl Acad Sci U S A, 111, E3277–3286.

99. Roithova, A., Feketova, Z., Vanacova, S. and Stanek, D. (2020) DIS3L2 and LSm proteins are involved in the surveillance of Sm ring-deficient snRNAs. Nucleic Acids Res, 48, 6184–6197.

100. Reimao-Pinto, M.M., Manzenreither, R.A., Burkard, T.R., Sledz, P., Jinek, M., Mechtler, K. and Ameres, S.L. (2016) Molecular basis for cytoplasmic RNA surveillance by uridylation-triggered decay in Drosophila. EMBO J, 35, 2417–2434.

101. Ramamurthy, L., Ingledue, T.C., Pilch, D.R., Kay, B.K. and Marzluff, W.F. (1996) Increasing the distance between the snRNA promoter and the 3’ box decreases the efficiency of snRNA 3’-end formation. Nucleic Acids Res, 24, 4525–4534.

102. Guiro, J. and Murphy, S. (2017) Regulation of expression of human RNA polymerase II-transcribed snRNA genes. Open biology, 7.

103. So, B.R., Wan, L., Zhang, Z., Li, P., Babiash, E., Duan, J., Younis, I. and Dreyfuss, G. (2016) A U1 snRNP-specific assembly pathway reveals the SMN complex as a versatile hub for RNP exchange. Nat Struct Mol Biol, 23, 225–230.

104. Venters, C.C., Oh, J.M., Di, C., So, B.R. and Dreyfuss, G. (2019) U1 snRNP Telescripting: Suppression of Premature Transcription Termination in Introns as a New Layer of Gene Regulation. Cold Spring Harbor perspectives in biology, 11.

105. Zhao, D.Y., Gish, G., Braunschweig, U., Li, Y., Ni, Z., Schmitges, F.W., Zhong, G., Liu, K., Li, W., Moffat, J. et al. (2016) SMN and symmetric arginine dimethylation of RNA polymerase II C-terminal domain control termination. Nature, 529, 48–53.

106. Kannan, A., Bhatia, K., Branzei, D. and Gangwani, L. (2018) Combined deficiency of Senataxin and DNA-PKcs causes DNA damage accumulation and neurodegeneration in spinal muscular atrophy. Nucleic Acids Res, 46, 8326–8346.

107. Reimer, K.A., Mimoso, C.A., Adelman, K. and Neugebauer, K.M. (2021) Co-transcriptional splicing regulates 3’ end cleavage during mammalian erythropoiesis. Mol Cell, 81, 998–1012 e1017.

108. Rosa-Mercado, N.A., Zimmer, J.T., Apostolidi, M., Rinehart, J., Simon, M.D. and Steitz, J.A. (2021) Hyperosmotic stress alters the RNA polymerase II interactome and induces readthrough transcription despite widespread transcriptional repression. Mol Cell, 81, 502–513 e504.

109. Alpert, T., Straube, K., Carrillo Oesterreich, F., Herzel, L. and Neugebauer, K.M. (2020) Widespread Transcriptional Readthrough Caused by Nab2 Depletion Leads to Chimeric Transcripts with Retained Introns. Cell Rep, 33, 108324.

110. Ottesen, E.W., Luo, D., Seo, J., Singh, N.N. and Singh, R.N. (2019) Human Survival Motor Neuron genes generate a vast repertoire of circular RNAs. Nucleic Acids Res, 47, 2884–2905.

111. Pagliarini, V., Jolly, A., Bielli, P., Di Rosa, V., De la Grange, P. and Sette, C. (2020) Sam68 binds Alu-rich introns in SMN and promotes pre-mRNA circularization. Nucleic Acids Res, 48, 633–645.

112. Arnold, M., Bressin, A., Jasnovidova, O., Meierhofer, D. and Mayer, A. (2021) A BRD4-mediated elongation control point primes transcribing RNA polymerase II for 3’-processing and termination. Mol Cell, 81, 3589–3603 e3513.

113. Drexler, H.L., Choquet, K., Merens, H.E., Tang, P.S., Simpson, J.T. and Churchman, L.S. (2021) Revealing nascent RNA processing dynamics with nano-COP. Nat Protoc, 16, 1343–1375.

114. Osman, E.Y., Van Alstyne, M., Yen, P.F., Lotti, F., Feng, Z., Ling, K.K., Ko, C.P., Pellizzoni, L. and Lorson, C.L. (2020) Minor snRNA gene delivery improves the loss of proprioceptive synapses on SMA motor neurons. JCI Insight, 5.

115. Kannan, A., Jiang, X., He, L., Ahmad, S. and Gangwani, L. (2020) ZPR1 prevents R-loop accumulation, upregulates SMN2 expression and rescues spinal muscular atrophy. Brain : a journal of neurology, 143, 69–93.

116. Mendell, J.R., Al-Zaidy, S., Shell, R., Arnold, W.D., Rodino-Klapac, L.R., Prior, T.W., Lowes, L., Alfano, L., Berry, K., Church, K. et al. (2017) Single-Dose Gene-Replacement Therapy for Spinal Muscular Atrophy. The New England journal of medicine, 377, 1713–1722.

117. Finkel, R.S., Chiriboga, C.A., Vajsar, J., Day, J.W., Montes, J., De Vivo, D.C., Yamashita, M., Rigo, F., Hung, G., Schneider, E. et al. (2016) Treatment of infantile-onset spinal muscular atrophy with nusinersen: a phase 2, open-label, dose-escalation study. Lancet, 388, 3017–3026.

118. Darras, B.T., Masson, R., Mazurkiewicz-Beldzinska, M., Rose, K., Xiong, H., Zanoteli, E., Baranello, G., Bruno, C., Vlodavets, D., Wang, Y. et al. (2021) Risdiplam-Treated Infants with Type 1 Spinal Muscular Atrophy versus Historical Controls. The New England journal of medicine, 385, 427–435.

119. Baranello, G., Darras, B.T., Day, J.W., Deconinck, N., Klein, A., Masson, R., Mercuri, E., Rose, K., El-Khairi, M., Gerber, M. et al. (2021) Risdiplam in Type 1 Spinal Muscular Atrophy. The New England journal of medicine, 384, 915–923.

120. Hosseinibarkooie, S., Schneider, S. and Wirth, B. (2017) Advances in understanding the role of disease-associated proteins in spinal muscular atrophy. Expert review of proteomics, 14, 581–592.

121. Sumner, C.J. and Crawford, T.O. (2018) Two breakthrough gene-targeted treatments for spinal muscular atrophy: challenges remain. The Journal of clinical investigation, 128, 3219–3227.

122. Talbot, K. and Tizzano, E.F. (2017) The clinical landscape for SMA in a new therapeutic era. Gene Ther, 24, 529–533.

123. Costa-Roger, M., Blasco-Perez, L., Cusco, I. and Tizzano, E.F. (2021) The Importance of Digging into the Genetics of SMN Genes in the Therapeutic Scenario of Spinal Muscular Atrophy. Int J Mol Sci, 22.

124. Blatnik, A.J., 3rd, McGovern, V.L. and Burghes, A.H.M. (2021) What Genetics Has Told Us and How It Can Inform Future Experiments for Spinal Muscular Atrophy, a Perspective. Int J Mol Sci, 22.

125. Wirth, B. (2021) Spinal Muscular Atrophy: In the Challenge Lies a Solution. Trends Neurosci, 44, 306–322.

126. Janzen, E., Mendoza-Ferreira, N., Hosseinibarkooie, S., Schneider, S., Hupperich, K., Tschanz, T., Grysko, V., Riessland, M., Hammerschmidt, M., Rigo, F. et al. (2018) CHP1 reduction ameliorates spinal muscular atrophy pathology by restoring calcineurin activity and endocytosis. Brain : a journal of neurology, 141, 2343–2361.

## SUPPLEMENTARY REFERENCES

1. Jedrzejowska, M., Gos, M., Zimowski, J.G., Kostera-Pruszczyk, A., Ryniewicz, B. and Hausmanowa-Petrusewicz, I. (2014) Novel point mutations in survival motor neuron 1 gene expand the spectrum of phenotypes observed in spinal muscular atrophy patients. Neuromuscul Disord, 24, 617–623.

2. Qu, Y.J., Bai, J.L., Cao, Y.Y., Wang, H., Jin, Y.W., Du, J., Ge, X.S., Zhang, W.H., Li, Y., He, S.X. et al. (2016) Mutation Spectrum of the Survival of Motor Neuron 1 and Functional Analysis of Variants in Chinese Spinal Muscular Atrophy. J Mol Diagn, 18, 741–752.

3. Gupta, K., Wen, Y., Ninan, N.S., Raimer, A.C., Sharp, R., Spring, A.M., Sarachan, K.L., Johnson, M.C., Van Duyne, G.D. and Matera, A.G. (2021) Assembly of higher-order SMN oligomers is essential for metazoan viability and requires an exposed structural motif present in the YG zipper dimer. Nucleic Acids Res, 49, 7644–7664.

4. Shao, M. and Kingsford, C. (2017) Accurate assembly of transcripts through phase-preserving graph decomposition. Nature biotechnology, 35, 1167–1169.

5. Tang, A.D., Soulette, C.M., van Baren, M.J., Hart, K., Hrabeta-Robinson, E., Wu, C.J. and Brooks, A.N. (2020) Full-length transcript characterization of SF3B1 mutation in chronic lymphocytic leukemia reveals downregulation of retained introns. Nature communications, 11, 1438.

